# Machine learning-augmented molecular dynamics simulations (MD) reveal insights into the disconnect between affinity and activation of ZTP riboswitch ligands

**DOI:** 10.1101/2024.09.13.612887

**Authors:** Christopher R. Fullenkamp, Shams Mehdi, Christopher P. Jones, Logan Tenney, Patricio Pichling, Peri R. Prestwood, Adrian R. Ferré-D’Amaré, Pratyush Tiwary, John S. Schneekloth

**Affiliations:** Chemical Biology Laboratory, National Cancer Institute, Frederick, MD, USA; Biophysics Program and Institute for Physical Science and Technology, University of Maryland, College Park 20742, USA; Laboratory of Nucleic Acids, National Heart, Lung, and Blood Institute, National Institutes of Health, Bethesda, MD, USA; Department of Chemistry and Biochemistry and Institute for Physical Science and Technology, University of Maryland, College Park 20742, USA; University of Maryland Institute for Health Computing, Bethesda, Maryland 20852, USA

## Abstract

The challenge of targeting RNA with small molecules necessitates a better understanding of RNA-ligand interaction mechanisms. However, the dynamic nature of nucleic acids, their ligand-induced stabilization, and how conformational changes influence gene expression pose significant difficulties for experimental investigation. This work employs a combination of computational and experimental methods to address these challenges. By integrating structure-informed design, crystallography, and machine learning-augmented all-atom molecular dynamics simulations (MD) we synthesized, biophysically and biochemically characterized, and studied the dissociation of a library of small molecule activators of the ZTP riboswitch, a ligand-binding RNA motif that regulates bacterial gene expression. We uncovered key interaction mechanisms, revealing valuable insights into the role of ligand binding kinetics on riboswitch activation. Further, we established that ligand on-rates determine activation potency as opposed to binding affinity and elucidated RNA structural differences, which provide mechanistic insights into the interplay of RNA structure on riboswitch activation.

## INTRODUCTION

RNA is known to form complex secondary and tertiary structures with regulatory roles such as influencing stability^1,2^, splicing^3–5^, and gene expression^6–9^. Furthermore, folded three-dimensional structures of RNA can form hydrophobic pockets^10^ that can be targeted with small molecules^11–13^. As a result, RNA has reemerged as a therapeutic intervention point for diseases currently ‘undrugged’ at the protein level^14–16^. However, our knowledge about RNA-small molecule interactions and how small molecule binding influences RNA structure and function is limited compared to protein-ligand interactions. For example, a disconnect between binding affinity and activity in biochemical/biological assays for RNA ligands can be observed but is often difficult to rationalize, even when high-resolution structures are available. Computational approaches that incorporate RNA dynamics could help address this challenge by successfully dovetailing experimental results with accurate, long-timescale simulations.^17–21^

The regulatory functions of RNA depend on the interplay of metastable states within highly dynamic structural ensembles^22^. Typically, biophysical methods developed for investigating protein-ligand interactions are adopted for studying RNA^23–25^, but these methods are not always compatible with interrogating dynamic RNA-ligand interactions. However, advances in single-molecule FRET assays^26–30^, and computational methods have provided tools that enable the investigation of RNA structural dynamics^31^ and the discovery of small molecules targeting RNA dynamic ensembles^32^. Molecular dynamics (MD) is a widely used computational method that has shown considerable success in studying biophysical problems of significance.^33^ Recent advances in computational hardware, including specialized supercomputers^34^ and GPUs^35^ have facilitated spatial parallelization and enabled the implementation of all-atom MD for large systems. Such approaches can provide atomistic insights about transient phenomena that are elusive to experimental observations and aid in the development of RNA-targeting small molecules. However, studying long time-scale rare events such as ligand binding/unbinding events,^36^ and slow conformational changes in biomolecules^37^ can be challenging due to the sequential nature of time and here we applied machine learning-augmented enhanced sampling to address this problem.^38^

Traditionally, optimization of protein-targeting small molecules has relied heavily on structure-guided design using co-crystal structures of protein-ligand complexes^39,40^. However, compared to proteins, there are few co-crystal structures of RNA-ligand complexes solved,^41^ and this has limited the use of structure-guided design for optimizing RNA-targeting molecules to far fewer instances^12,32,42–44^. Bacterial riboswitches are a well-understood class of structured mRNA motifs that control essential metabolic pathways for bacterial growth and virulence. Riboswitches control gene expression by sensing the intracellular concentration of a cognate ligand and, upon binding, undergo a conformational change that regulates gene expression^45^. Riboswitches often have well-defined three-dimensional structures that contain hydrophobic pockets that can be targeted with drug-like small molecules^11^ and have been implicated as potentially novel antimicrobial targets^46^. Due to the well-characterized functional effect that ligand binding has on gene expression and the availability of co-crystal structures of riboswitch-ligand complexes, riboswitches are ideal model systems for investigating RNA-small molecule binding interactions and structure-guided medicinal chemistry efforts.

In this work, we used structure-informed design to synthesize a focused library of 27 small molecules that bind to and activate the *Fusobacterium ulcerans* ZTP riboswitch *in vitro*.^47^ However, upon further investigation we observed a poor correlation between *in vitro* riboswitch activation and ligand affinity among a library of eight novel ligands. To investigate this disconnect, we first co-crystallized two ligands with the RNA and used machine learning-augmented molecular dynamics simulations^38^ to investigate the diverse dissociation mechanisms of seven synthetic ligands and the cognate ligand ZMP. From these simulations for each ligand, we calculated the rate of ligand binding (k_on_), which is challenging to determine experimentally, and found a strong correlation between k_on_ and *in vitro* riboswitch activation. Furthermore, comparison of ligand dissociation trajectories identified key differences between the dissociation mechanisms of synthetic ligands compared to cognate ligand ZMP. These differences correlated with *in vitro* riboswitch activation and provide mechanistic insights into the role of flexibility, and specific riboswitch residues in the observed differences among ligands.

## RESULTS

### Structure-informed Design of F. ulcerans ZTP riboswitch binders

Previously, isosteric replacement of the phosphate and ribose sugar moiety of ZMP with a pyridine group resulted in the discovery of compound **1** (**Figure 1c**)^48^. Compound **1** possessed a weaker affinity (K_D_ ∼600 nM) but was a stronger riboswitch activator (T_50_ ∼5.8 µM) than cognate ligand ZMP (K_D_ ∼324 nM and T_50_ ∼37 µM) in biochemical assays^48^. Intrigued by the disconnect between ligand affinity and activation, we examined the reported co-crystal structures of **ZMP** (PDB: 60D9) and **1** (PDB: 6WZS) bound to *F. ulcerans* ZTP riboswitch to rationally design new analogs. The binding pose of ZMP and **1** are very similar due to the conserved amino-amidocarboxamide (AICA) core. Analysis of the co-crystal structures indicated that due to the extent of burial of the imidazole core and the limited available volume, extensive modification of the AICA core would not be tolerated (**Figure 1a**). This observation is further supported by the inherent selectivity of the ZTP riboswitch for ZMP and ZTP over inosine, which is a downstream metabolic intermediate, and only a single carbon unit larger. Therefore, we directed our analysis to the solvent-exposed area around the pyridine of **1**. ZMP and **1** each make unique interactions within the binding pocket and could result in the observed disconnect between affinity and activation. The ribose and phosphate moieties of ZMP make hydrogen bonds with the 2’-OH of G63 and N4 of C69 (PDB: 60D9). In contrast, the pyridine moiety of **1** makes a *π*-*π* (black lines) stacking interaction with G63 and a putative hydrogen bond (purple line) with 2’-OH of G63 (**Figure 1b**). The loss of the hydrogen bond interaction with N4 of C69 and gain of the *π*-*π* stacking interaction resulted in a ∼2-fold loss in affinity for compound **1**, yet **1** had greater *in vitro* and *in vivo* activation of the *F. ulcerans* ZTP riboswitch^48^. Additionally, adjacent to the pyridine group is a cavity (**Figure 1a, labeled C1**) that could accommodate larger substituents.

**FIG. 1:**
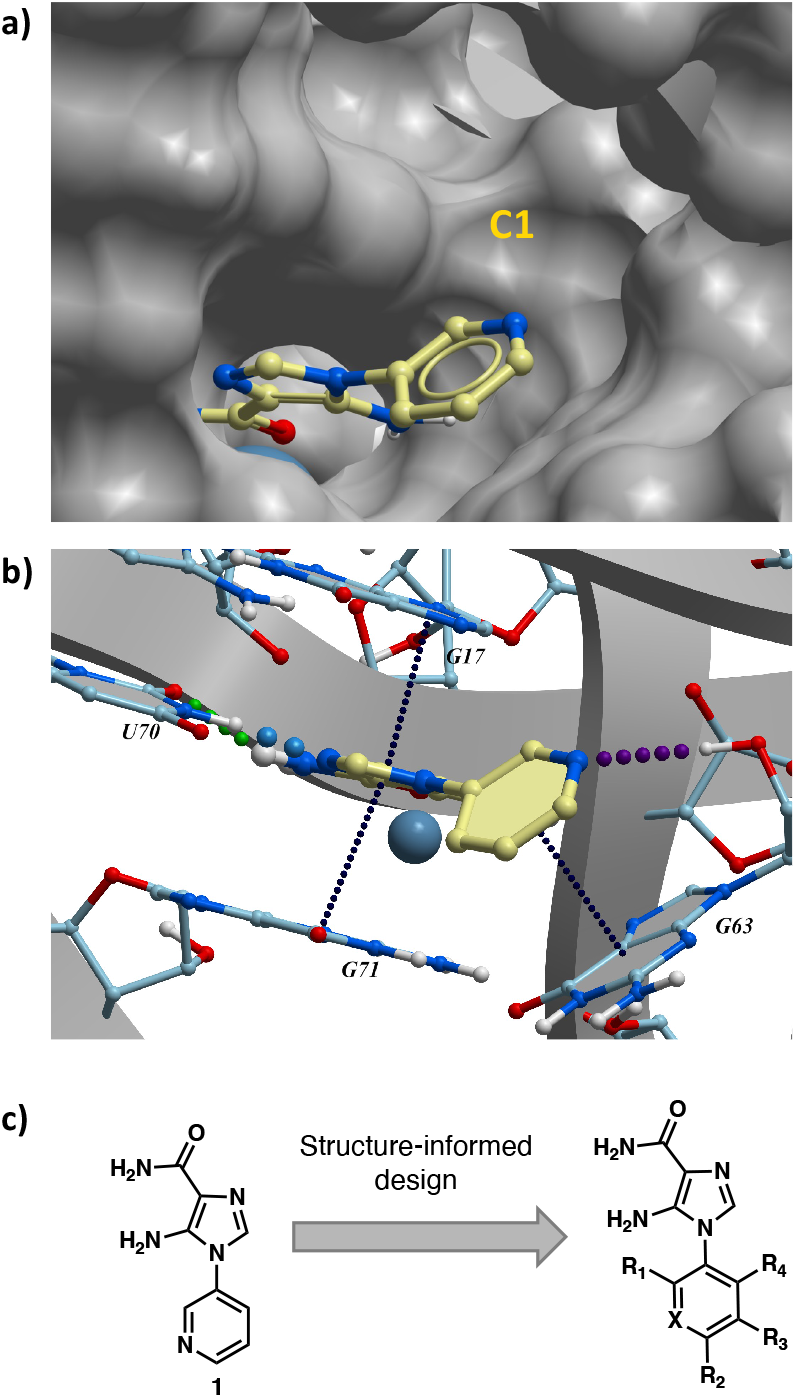
Structural analysis of 1 bound to ZTP riboswitch and overview of the structure-based design of synthetic binders of the *F. ulcerans* ZTP riboswitch. (a) The binding pose of 1 in complex with *F*.*ulcerans* ZTP riboswitch (PDB: 6WZS). Analysis indicates limited volume is available to accommodate additional modifications around the AICA core, and a potential cavity is present adjacent to the pyridine moiety of 1 (labeled C1), which could accommodate larger substituents. (b) The binding mode of **1** highlights the key hydrogen bonding and *π*-*π* stacking interactions (black dotted lines) of **1** with bases U17, G17, G71, and G63. A proposed putative hydrogen bond interaction with the 2’-OH of G63 and the pyridine moiety is highlighted with a purple dotted line. (c) Overview of the structural modification to **1** resulting in novel activators of *F. ulcerans* ZTP riboswitch highlighting the modifications made around the pyridine ring of **1**.

### Synthesis and affinity measurement of designed analogs

Based on the analysis of the co-crystal structures of **ZMP** (PDB: 6OD9) and **1** (PDB: 6WZS) in complex with the *F. ulcerans* ZTP riboswitch, we designed and synthesized a library of 27 synthetic analogs that incorporated minor changes in (1) the AICA core or (2) the pyridine of **1** (**Figure 1c**)and **Table I**, analogs **4-28**). The library of analogs was accessed by one of three synthetic routes. Compound **2** was afforded by cycloaddition of 3-azidopyridine with 2-cyanoacetamide in 30% yield^49^. Compound **3** was accessed by nucleophilic aromatic substitution between 1H-imidazole-4-carboxamide and 4-chloropyridine in modest yields, and compounds **4-28** were synthesized by reaction of 2-amino-cyanoacetamide with triethyl orthoformate followed by the addition of substituted anilines, quinolines, or napthyridines to afford the desired analogs in modest to good yields^48,50^ (**Figure S12**). With our library of analogs in hand, the equilibrium dissociation constant to the aptamer domain of *F. ulcerans* ZTP riboswitch was measured using isothermal titration calorimetry (ITC), following the previously described method^48^.

**TABLE 1:**
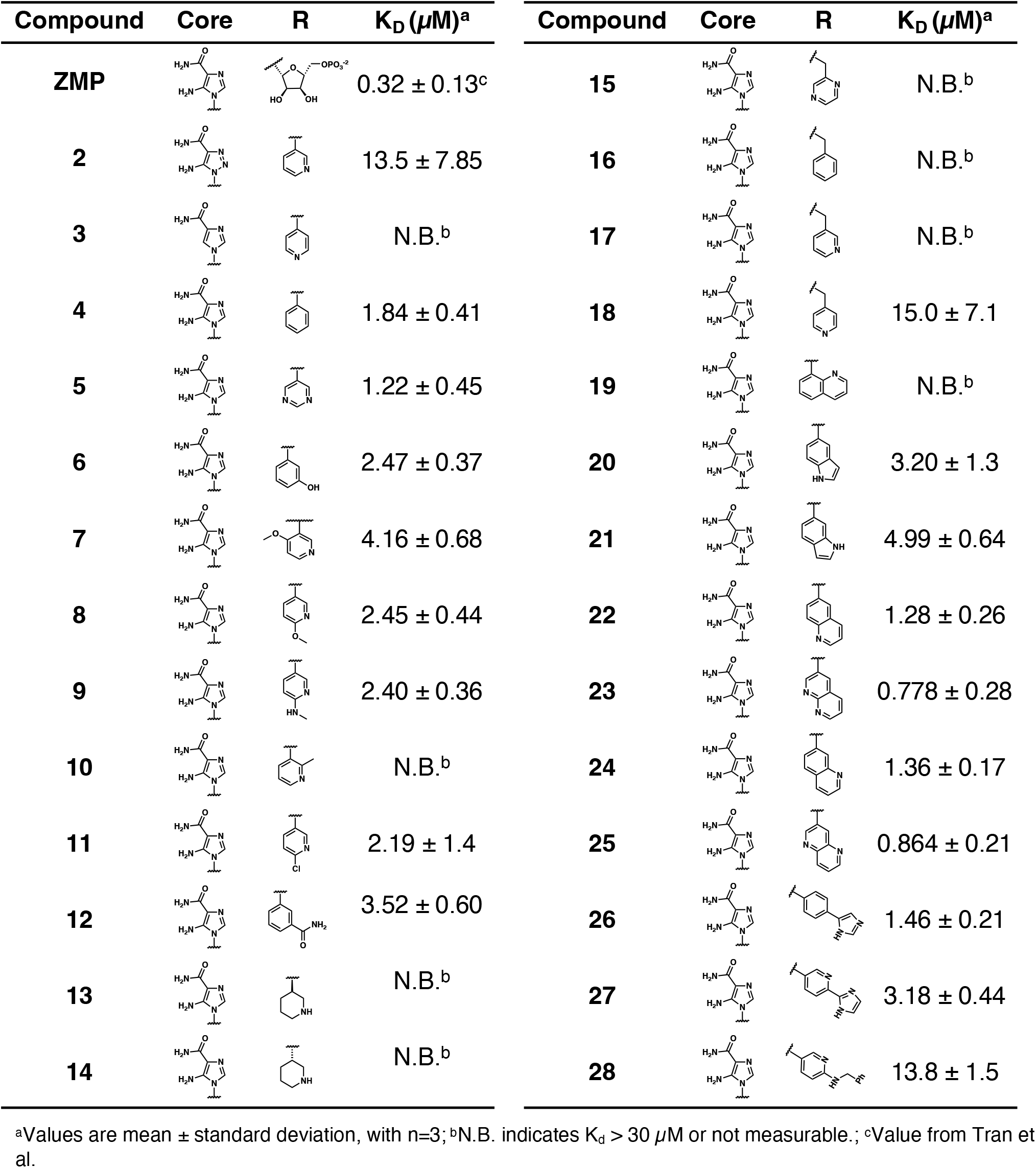
Structure-Informed Synthetic Analogs of m-pyridinyl AICA and ITC *K*_*D*_.

| Compound | Core | R | $K_D$ ($\mu\text{M}$)<sup>a</sup> | Compound | Core | R | $K_D$ ($\mu\text{M}$)<sup>a</sup> |
| --- | --- | --- | --- | --- | --- | --- | --- |
| **ZMP** | | | $0.32 \pm 0.13^c$ | **15** | | | N.B.<sup>b</sup> |
| **2** | | | $13.5 \pm 7.85$ | **16** | | | N.B.<sup>b</sup> |
| **3** |  |  | N.B.<sup>b</sup> | **17** |  |  | N.B.<sup>b</sup> |
| **4** | | | $1.84 \pm 0.41$ | **18** | | | $15.0 \pm 7.1$ |
| **5** | | | $1.22 \pm 0.45$ | **19** | | | N.B.<sup>b</sup> |
| **6** | | | $2.47 \pm 0.37$ | **20** | | | $3.20 \pm 1.3$ |
| **7** | | | $4.16 \pm 0.68$ | **21** | | | $4.99 \pm 0.64$ |
| **8** | | | $2.45 \pm 0.44$ | **22** | | | $1.28 \pm 0.26$ |
| **9** | | | $2.40 \pm 0.36$ | **23** | | | $0.778 \pm 0.28$ |
| **10** | | | N.B.<sup>b</sup> | **24** | | | $1.36 \pm 0.17$ |
| **11** | | | $2.19 \pm 1.4$ | **25** | | | $0.864 \pm 0.21$ |
| **12** | | | $3.52 \pm 0.60$ | **26** | | | $1.46 \pm 0.21$ |
| **13** | | | N.B.<sup>b</sup> | **27** | | | $3.18 \pm 0.44$ |
| **14** | | | N.B.<sup>b</sup> | **28** | | | $13.8 \pm 1.5$ |

<sup>a</sup>Values are mean ± standard deviation, with $n=3$ ; <sup>b</sup>N.B. indicates $K_d > 30 \mu\text{M}$ or not measurable.; <sup>c</sup>Value from Tran et al.

Because the AICA core was deeply buried in the binding pocket(**Figure 1a**), only minor modifications to the core were attempted. Single-atom substitution (C to N) in the imidazole core, compound **2**, resulted in a ∼22-fold loss in binding affinity (K_D_ = 13.7 7.8 µM) compared to **1** (K_D_ = 0.60 0.06 µM). In addition, the removal of the 5-amino group, compound **3**, resulted in a complete loss of binding (K_d_ *>* 30 µM) (**Table I**). These modifications highlight the importance of the AICA core for riboswitch-ligand recognition and binding. AICA core modifications were not tolerated, we directed our efforts to replace the pyridine moiety of compound **1** with different aromatic and saturated ring systems. These efforts resulted in analogs with a range of affinities from ∼800 nM to *>*30 µM ((**Table I, 4-28**).

Replacement of the pyridine in compound **1** for a phenyl group, compound **4**, resulted in a ∼3-fold loss of binding, highlighting the importance of the previously reported putative hydrogen bonding interaction between the nitrogen atom in the pyridine ring of **1** and the 2’-OH of G63^48^. The substitution of pyridine for pyrimidine, compound **5**, had a ∼2-fold loss in affinity. In addition, replacement with piperidine, compounds **13** and **14**, resulted in a total loss of binding (K_D_ *>*30 µM), highlighting the importance of the *π*-*π* stacking interaction between the pyridine ring of **1** and G63. Furthermore, the substitution of the 2-position of the pyridine ring with electron-donating (methoxy (**8**) and N-methylamine (**9**)) or electron-withdrawing (Chloro (**11**)) resulted in a ∼4-fold loss in affinity compared to **1** (**Table I**).

Introduction of a methyl group ortho to the pyridinyl nitrogen atom of compound **1** or a methoxy group at position 4, compound **7**, to promote the ideal torsional angle for hydrogen bond formation with 2’-OH of G63, abolished binding (**Table I, 7** and **10**, K_D_’s *>*30 µM). Further, replacement of the meta-nitrogen of the pyridine with hydroxyl or an amide was tolerated but with a ∼5-6 fold loss in binding affinity (**6** and **12, Table I**). In addition, attempts to extend the ligand into cavity **C1** (**Figure 1b**), with benzylic aryl and quinoline moieties (compounds **15-19**), resulted in a ∼ 24-fold reduction in affinity for **18** and complete loss of binding K_D_*>*30 µM for **15-17** and **19**.

The extension of the pyridine moiety of **1** towards the opening of the binding cavity with quinolines (compounds **22** and **24**) and naphthyridines (compounds **23** and **25**) resulted in analogs with equivalent K_D_’s to **1** (K_D_ values of 0.778-1.38 µM versus 0.610 µM, respectively). In contrast, replacement with an indole, **20** and **21**, resulted in a ∼7-fold loss in affinity compared to **1**. Biaryl analogs (**26, 27**, and **28**) also resulted in a loss of affinity. Overall, we rationally designed and synthesized a library of 27 new ligands and identified two ligands, **23** and **25**, with similar affinity to **1** (K_D_ ∼800 nM for **23** and **25** compared to K_D_ ∼610 nM for **1**). Our library highlights how small modifications to the ligand structure can result in drastic effects on binding affinities to *F. ulcerans* ZTP riboswitch. With our newly identified riboswitch binding analogs in hand, we next evaluated their ability to activate the *F. ulcerans* ZTP riboswitch *in vitro*.

### Synthetic analogs activate ZTP riboswitch in vitro

To evaluate the activation potential of our synthetic ligands, we conducted single-round transcription termination assays with *F. ulcerans*m ZTP riboswitch using the previously reported methods.^48,51^ Based on the regulatory mechanism of the *F. ulcerans* ZTP riboswitch, as an activator of transcription, we expect to see greater read-through with increased binding of the RNA aptamer with ligand. The accumulation of the read-through transcript can be quantified and fitted to determine T_50_, the concentration at which the riboswitch is half activated.

ZMP was first retested and found to have a T_50_ in good agreement with the previously reported value (59 15.8 µM vs. 37 12 µM) (**Figure 2**)^48^. We then chose a subset of the newly designed synthetic ligands (**2, 23, 25, 26, 27** with a range of binding affinities (K_D_ ∼800 nM to 13 µM) to investigate their ability to activate transcriptional read-through. Compounds **2, 23, 25, 26**, and **27** all had lower T_50_ values (i.e., better activation) than ZMP, even though ZMP is a tighter binder. These results are similar to the previously reported observation between **1, 4-piperidinyl AICA**, and **ZMP** (**Figure 2**)^48^, in which **1** and **4-piperidinyl AICA** activate transcription to a greater extent than **ZMP**, even though both bind the riboswitch with a weaker affinity. A potential hypothesis explaining the observed disconnect between affinity and activation is riboswitch activation is driven by the rate of ligand binding (k_on_) and not overall affinity (K_D_). Using a single-molecule FRET-based assay to study riboswitch folding, it was previously demonstrated that ZMP’s activation of the ZTP riboswitch is driven by k_on_^29^; however, it has not been shown that this holds true across a panel of synthetic ligands.

**FIG. 2:**
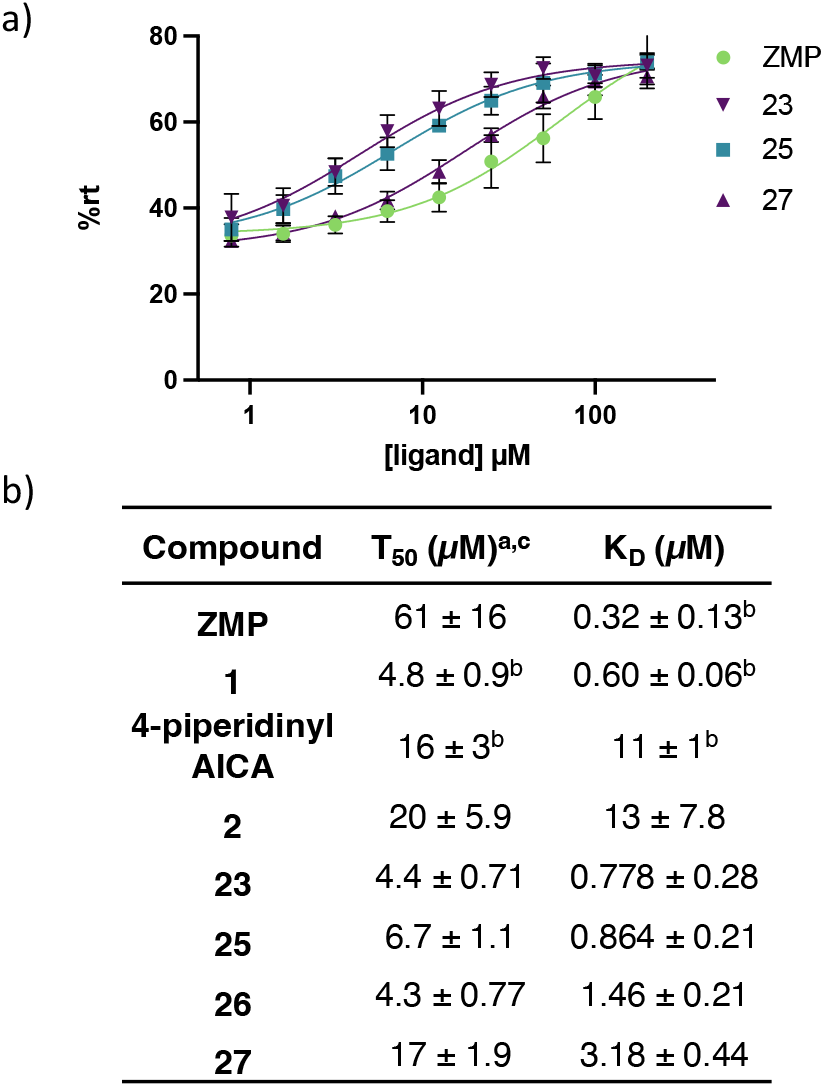
Single-Round Transcription Termination Efficiencies. a) Single-round transcription termination for **ZMP, 23, 25**, and **27** (n= 3, error bars denote standard deviations). b) Transcription termination efficiency (T50) and binding affinity (KD) for **ZMP, 1, 4-piperidinyl AICA, 2, 23**, and **25-27**. Here, the superscripts correspond to: ^*a*^values reported as mean ± standard deviation, n=3; ^*b*^ values obtained from Tran et al^48^; and ^*c*^individual single-round transcription termination titration curves for each analog are included in the SI.

### Investigation of binding kinetics by surface plasmon resonance

To investigate the role ligand binding kinetics of our synthetic analogs have on riboswitch activation, experimental determination of k_on_ for our synthetic ligands was attempted using surface plasmon resonance (SPR) experiments with the aptamer domain of the *F. ulcerans* ZTP riboswitch. Using both the traditional streptavidin reference channel subtraction method^52^ and the recently reported non-binding mutant RNA reference channel subtraction method^25^, we observed ligand binding of compound **26** with the aptamer domain of *F. ulcerans* ZTP riboswitch and obtained K_D_ values of ∼ 900 nM, which is in good agreement with our ITC value (K_D_ ∼ 1.46 ± 0.21 µM, **Supporting Figure S2**). However, the plotted response curve did not reach saturation and exceeded the theoretical max response for a 1:1 binding event by more than 2-fold, presumably due to non-specific binding or aggregation (**Supporting Figure S2**). In addition, k_on_ and K_off_ for **26** could not be extracted from the SPR response curves due to the observed steep slope for association and dissociation (**Supporting Figure S2**). Since kinetic information could not be obtained experimentally, we set out to investigate ligands via co-crystallography and use those models to facilitate the study of ligand binding kinetics by MD methods.

### Co-crystal structures of compounds 1 and 23 bound to the S. odontolytica ZTP riboswitch

Our previously reported structure of compound **1** was determined at 3.2 Å resolution bound to the *F. ulcerans* ZTP riboswitch^48^; thus, we sought to validate those findings in the *S. odontolytica* ZTP riboswitch, which has been reported to crystallize at higher resolution. We cocrystallized **1** with the *S. ondontolytica* ZTP riboswitch aptamer, solved the structure via molecular replacement, and refined the structure to 2.4 Å resolution (Table S1, Methods). In the model, the pyridine moiety is poised to hydrogen bond to the 2’-OH of G51 (3.0 Å, Fig. 3a), consistent with our previous findings. Nearby C52 is about 4.0 Å away from the pyridinyl group and hydrogen bonds with non-bridging phosphate oxygen (NBPO) O2 of C50 (3.0 Å). In the presence of ZMP, a hydrated magnesium ion makes inner sphere contacts with C52 (**Figure 3a**).

**FIG. 3:**
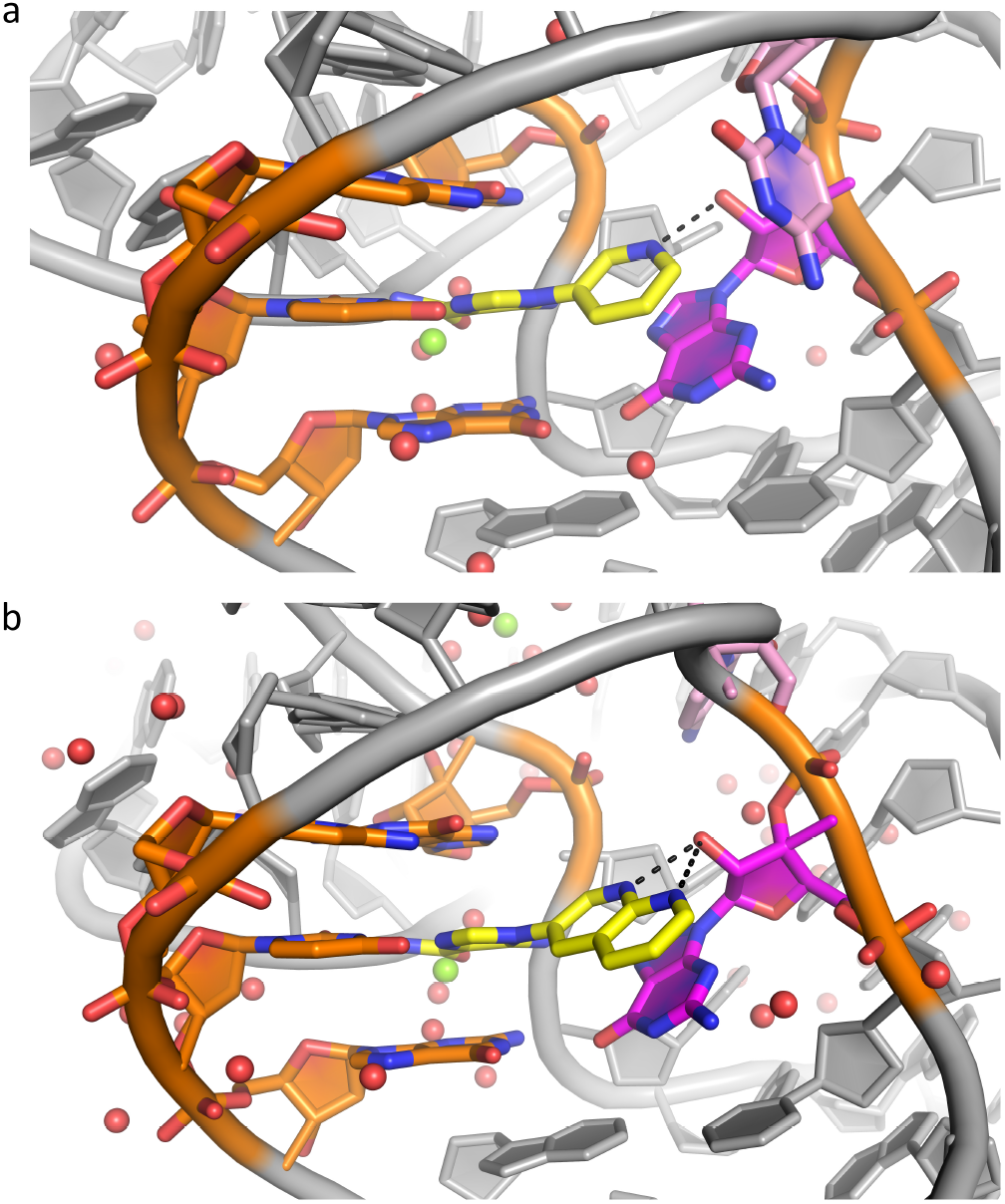
Co-crystal Structures of ZTP riboswitch in complex with synthetic analogs highlight conserved binding mode. a) Co-crystal Structure of Compound **1** yellow with *S*.*ondontolytica* ZTP riboswitch. The black dotted line indicates hydrogen bonding interaction with G51(magenta). b) Co-crystal Structure of Compound **23** (yellow) with *S. ondontolytica* ZTP riboswitch. The black dotted line indicates potential hydrogen bonding interactions between G51 (magenta) and the N1 and N8 of the napthyridine of **23**. In this model, C52 (pink) undergoes a conformational change, moving away from the napthyridine of **23**. Green spheres denote magnesium ions, and red spheres denote waters.

In contrast to **1**, for which co-crystals were readily obtained, larger synthetic ligands proved more challenging. We co-crystallized **23** to the *S. ondontolytica* RNA in different conditions, habits, and space groups (Methods). The structure determined at 2.2 Å resolution was partially solved and refined due to the presence of weakly resolved copies in the asymmetric unit (Table S1). As in **1, 23** is poised to hydrogen bond with the 2’-OH of G51 (2.9 Å) via the N1 of **23**, and the AICA moiety is bound as for **1** (**Figure 3b**). An additional hydrogen bond is also possible between N8 of **23** and the 2’-OH of G51 (3.2 Å). However, likely due to steric clashing with the larger ligand, C52 no longer hydrogen bonds to NBPO O2 and instead pairs with G34, which is unpaired in the presence of ZMP and **1**. In addition, in the presence of **23**, loop residues A16-19 undergo a conformational change that is involved in making crystal contacts. The conformation of the residue after G51 varies among ZTP riboswitch sequences. Consequently, its conformation participating in an H-bond with C50 (*i*.*e*., as in **Fig. 3** and^53^) has only been observed in S. odontolytica.

### Insights into RNA-ligand interaction mechanisms using all-atom Molecular Dynamics (MD)

As the experimental apo-structure remains unresolved and conducting an accurate MD simulation of the association event is prohibitively expensive due to the substantial associated barrier,^54^ we simulated and investigated dissociation events instead. We achieved this by learning a meaningful description of system behavior by implementing the state predictive information bottleneck^55^ (SPIB), a machine-learning^56^ (ML) framework for identifying a low-dimensional reaction coordinate and used it to implement well-tempered metadynamics^55,57–61^ (WT-MetaD) for accelerating MD simulations. We performed this ML-assisted WT-MetaD to observe 16 independent dissociation events for each of the 8 ligands chosen for computational studies (**ZMP, 1, 4-piperidinyl AICA, 2, 23, 25, 26**, and **27**). Specifics regarding the MD system configuration, feature selection, training of ML model for constructing reaction coordinates, and WT-MetaD implementation can be found in the supplementary information (SI).

Analysis of the final dissociation trajectories unraveled key mechanistic details about the individual RNA-ligand interactions as depicted in **Fig**. (4, 6, 5). In the bound conformation for the ligands, the RNA structural *Mg*^2+^ ion was observed to coordinate with 3 water molecules (**Fig**. 4(a,b)), the amide of the ligands, and with the phosphate groups of U16 and C35, respectively. Eventually, a fourth water molecule coordinates with the *Mg*^2+^ ion, leading to a loss of interaction with the ligands, allowing for dissociation (**Fig**. 4(c)).

**FIG. 4:**
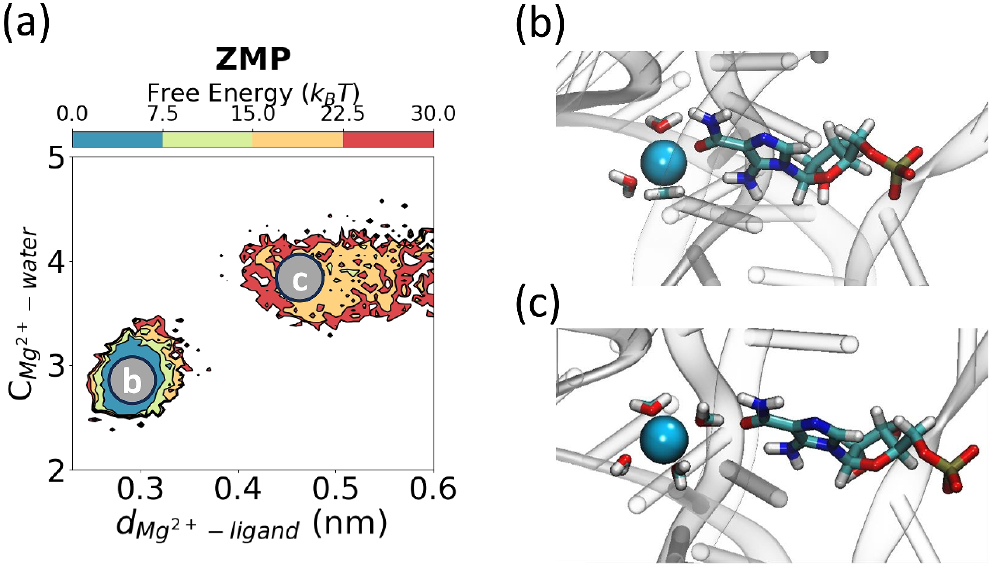
Change in structural *Mg*^2+^ coordination during ligand dissociation. (a) Free energy projections of water-*Mg*^2+^ distance as **ZMP** exits. (b,c) *Mg*^2+^ ion hydration as the ligand exits. The structures correspond to the labels in panel (a).

**FIG. 5:**
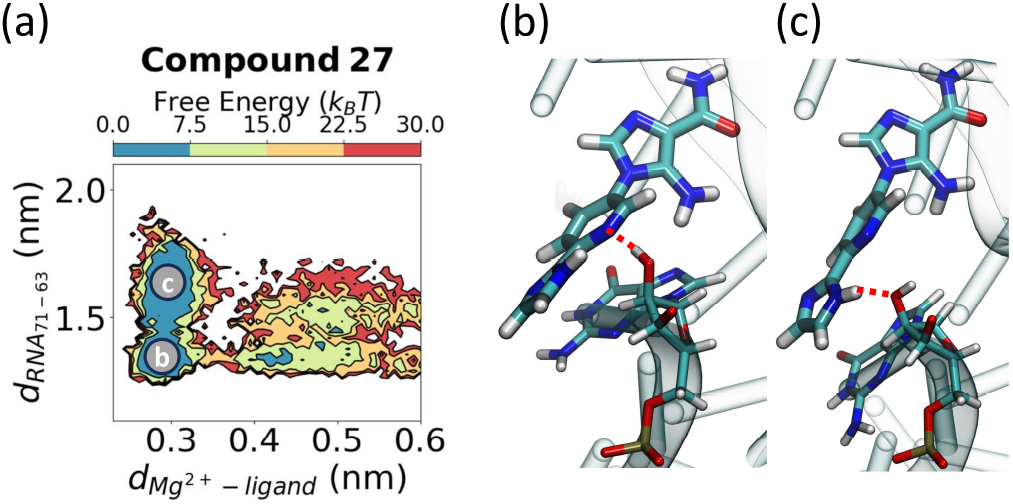
Synthetic derivatives exhibit different behavior in the bound conformation. (a) Free energy projected along the distance between the backbone of G71 and nucleobase of G63 (y-axis) as **compound 27** exits the binding site, as measured by the *Mg*^2+^-ligand distance. In the bound conformation, there are two metastable states, highlighted in (b,c) showing alternating *π* − *π* stacking and hydrogen bonding interactions with G63 between the outer ligand moieties. The red dotted lines in panels (b,c) indicate hydrogen bonds.

After decoordination between the *Mg*^2+^ and the oxygen atom of the amide moiety, the dissociation mech-anism differs between **ZMP** and synthetic ligands (**1, 4-piperidinyl AICA, 2, 23, 25, 26, 27**). **Fig**. 6(a) shows the pyridine ring of **compound 1** forms a *π* − *π* stacking interaction with G63, allowing G63 and G71 to remain in close proximity via a noncanonical interaction (**Fig**. 6(e)). However, for **ZMP**, the sugar moiety sits between G63 and G71 (**Fig**. 6(b,f)). When the synthetic ligands exit the cavity (**Fig**. 6(c)), the distance between the backbone of G71 and nucleobase of G63 increases (**Fig**. 6(c,e)). Here, we noted that this increase in distance between G63 and G71 for the synthetic ligands occurs after the hydration of the *Mg*^2+^ ion, as discussed previously. However, for **ZMP**, as the ligand exits, the average distance between G63 and G71 decreases (**Fig**. 6(f)). In this case, after the hydration of *Mg*^2+^, the volume available to **ZMP** in the cavity decreases, and **ZMP** undergoes an end-to-end contraction before exiting the cavity (**Fig**. 6(g,h)).

**FIG. 6:**
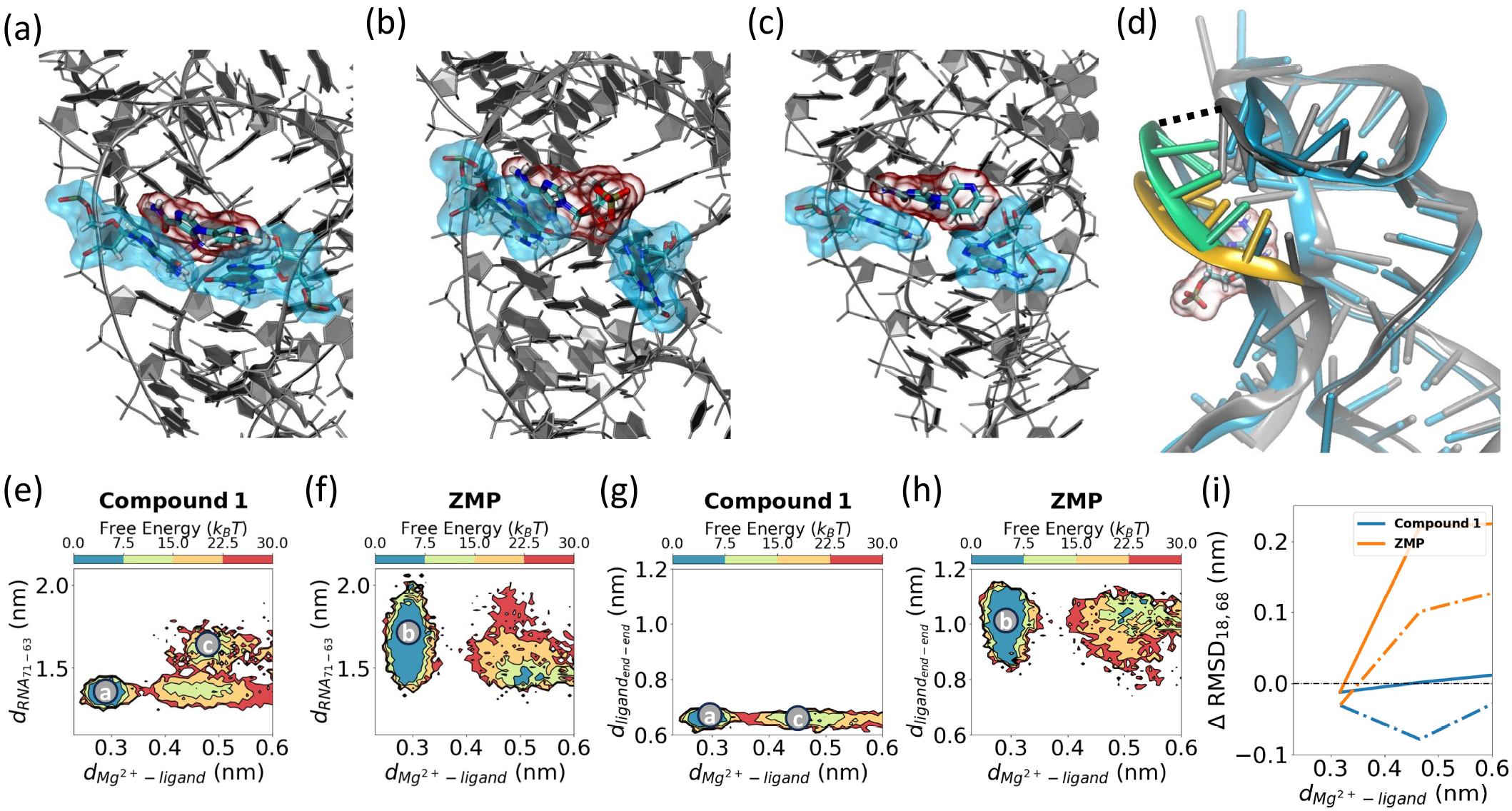
MD simulations of RNA-ligand interactions. (a) **Compound 1** bound to ZTP riboswitch with pyridine moiety exhibiting *π* − *π* stacking interactions with G63. In this conformation, RNA residues G63 and G71 (highlighted using cyan clouds) are in close proximity. (b) **ZMP** bound to ZTP riboswitch with RNA residues G63 and G71 (highlighted in cyan) separated, (c) dissociation of **1** involves opening of the G63 nucleobase to allow ligand exit, (d) increased flexibility of the P4 domain (green, and yellow) of ZTP riboswitch during ligand dissociation. Two specific residues in the P4 domain (A18 and U68) are highlighted with a dotted line. Free energies are projected on the distance between the backbone of G71 and nucleobase of G63 as the ligands exit for (e) **1** and (f) **ZMP**, respectively. In the bound conformation (panels a,c), the spread in this distance is smaller for **1** compared to **ZMP**. As the ligand exits (panel c), this distance increases for **1**. End-to-end distance for (g) **1**, and (h) **ZMP**, respectively. The rotatable bonds present in **ZMP** allow higher flexibility. (i) Δ RMSD for A18 (solid line), and U68 (dotted line) show that **ZMP** induces significant flexibility to P4 domain compared to **1** during exit.

**FIG. 7:**
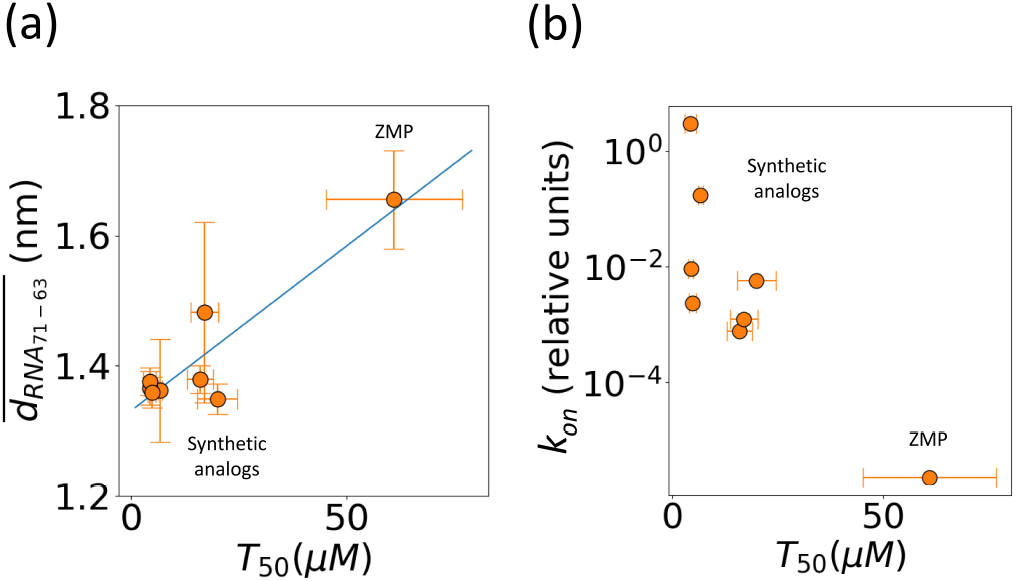
Correlation of MD-derived k_on_ and nucleobase distances with experimental measurements of transcriptional readthrough. (a) Correlation of T_50_ (x-axis) with between G71 and G63 (y-axis) in ligand-bound conformation. (b) Plot of T_50_ values from transcription termination assays (x-axis) versus relative k_on_ from frequent WT-MetaD (y-axis).

The ligands exit the cavity through a pathway close to the P4 domain (**Fig**. 6(d,i)). We observed **ZMP** inducing higher flexibility in this domain compared to the synthetic derivatives. This is quantified by Δ RMSD (Root-Mean-Square Deviation), which is defined as the change in RNA-ligand distance as the ligand exits when compared to a long (180 ns) unbiased simulation of a ligand-removed structure. Larger derivatives (**23, 25-27**) also induced higher flexibility to a few of the residues in the P4 domain, but to a lesser extent than observed for **ZMP**. A detailed summary of Δ RMSD calculations for all residues and for each ligand is provided in the SI (**Supporting Figure 6,7**).

In addition to the differences between **ZMP** and the synthetic derivatives computationally studied in this work, synthetic derivatives also displayed distinct behavior. **Compound 27** (**23**,**25** to a lesser extent) exists as two metastable states in the bound state (**Fig**. 5). We designed **Compound 27** with both a hydrogen bond donor and acceptor with the intention of introducing a bivalent hydrogen bond interaction. However, we observed two metastable states where the *π* − *π* stacking interaction alternates between the inner pyridine and outer imidazole groups due to the formation of distinct hydrogen bonds in the bound state highlighted by dotted lines in (**Fig**. 5(b,c)).

### Computational results correlate with activation potency

In addition to investigating RNA-ligand interaction mechanisms from MD simulations, we computed relative residence times (t_rel_) from the WT-MetaD time-dependent bias deposition for each of the ligands. This was done using the acceleration factor approach^57^ and is discussed in detail in the SI. Using experimentally determined K_D_, and computational rate of dissociation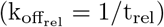, we computed the relative rate of association of the ligands 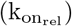 using the relation,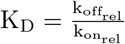. Here, we observed a Spearman’s rank correlation coefficient of − 0.71 between 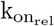 and T_50_ values from transcription termination assays. This affirms a monotonic relationship between 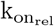 and riboswitch activation potency. That is, a higher 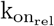 is correlated with smaller T_50_ (more potent activation). Additionally, when comparing the logarithm of the 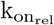 with T_50_, we observed a Pearson correlation coefficient of − 0.83, a strong linear relationship (**Fig**. 7(b)).

Furthermore, we also observed that G63 became dynamic, particularly for the larger ligands, and was correlated with the movement of the ligand from the cavity to the P4 domain prior to exit. We quantified this behavior of G63 in the ligand-bound state as a weighted average (WT-MetaD Boltzmann weights) of the distance between the G71 backbone and G63 nucleobase for all ligands. We observed a Pearson correlation coefficient of 0.91 with T_50_ values (**Fig**. 7(a)). While this supports a significant role for G63 in ligand interaction, the correlation is heavily weighted by ZMP, owing to the lack of additional compounds with T_50_ values worse than those investigated here.

## DISCUSSION

In this work, we demonstrated how structure-informed design can be used to identify new ligands that bind to and activate the *F. ulcerans* ZTP riboswitch *in vitro*. We identified analogs **23, 25**, and **26**, which have equivalent affinities (K_D_ ∼800 nM, **Table I**) and *in vitro* riboswitch activation (T_50_ ∼5 µM, **Figure 2**) as the previously reported analog **1** (K_D_ ∼610 nM and T_50_ of 4.8 µM)^48^. We did not identify analogs with improved affinity or activation than **1** in part due to the limited volume available for modification around the AICA core and the solvent accessibility of the binding cavity around the pyridine moiety of **1** (**Figure 1a**).)). While the C1 pocket offers a direction for such improvement, this region also varies among the solved structures^53,62,63^. The difficulty in identifying novel analogs with improved affinity has also been encountered with other riboswitch systems^13^. However, from our synthetic endeavor, we gained insights into the type of modifications that promote riboswitch-ligand interaction. We observed that aromatic and heteroaromatic rings can act as suitable isosteric replacements for the ribose and phosphate moiety of **ZMP**, which is consistent with previously reported studies for the ZTP riboswitch^48^ and FMN riboswitch^11,64^. In addition, from our molecular dynamics simulations, we observed two metastable states for the bound conformation of compound **27** (and **23** and **25** to a lesser extent). We intended with the design of compound **27** to shift the hydrogen bonding interaction from a hydrogen bond acceptor from pyridine in **1** to a hydrogen bond donor from the NH of the imidazole in **27**, and from our simulations, we were able to observe this binding mode shift without obtaining a co-crystal structure.

Furthermore, a disconnect between ligand affinity and riboswitch activation was observed for our synthetic analogs (**2, 23**, and **25-27**) and **ZMP** (**Figure 2**.) Our synthetic ligands (**2, 23**, and **25-27**) all displayed greater *in vitro* activation than cognate ligand **ZMP** even though their K_D_s are ∼2-20-fold weaker. The disconnect is hypothesized to result from differences in the k_on_ of ligand, the rate of transcription, and the rate of RNA unfolding^65,66^. The *F. ulcerans* ZTP riboswitch undergoes co-transcriptional folding and senses the intracellular concentration of cognate ligand **ZMP** over an ∼5-10 nucleotide window; it is expected that riboswitch activation would be influenced more by k_on_ than K_D_ since activation occurs too quickly for binding to reach equilibrium^29,67^. Our molecular dynamics (MD) simulations allowed for the calculation of ligand binding kinetics, and we found a correlation between k_on_ and riboswitch activation (T_50_) (**Fig**. 7), but in addition to kinetics, we gained insights into the differences in RNA structural dynamics during ligand dissociation.

Unlike previous studies which used MD simulations to analyze only the RNA by itself^68^or RNA-cognate ligand^29^, we performed a comparative study of our synthetic ligands and **ZMP**, and found key differences in their dissociation trajectories (**Fig**. 6). The main difference observed is the extension of the P4 domain upon ZMP exiting the cavity and the correlation of the distances between G63 and G71 with riboswitch activation. Residues in the P4 domain are part of the interdomain pseudoknot that forms the binding pocket but also overlap with the terminator hairpin sequence. The ZTP riboswitch undergoes a ligand-gated competitive strand displacement to form the terminator hairpin during transcription, and the binding of ZMP stabilizes the pseudoknot, disfavoring terminator hairpin formation^67^. We hypothesize that the differences in the G63-G71 distances and the observed extension of the P4 domain during ZMP dissociation may promote or aid internal strand displacement and termination hairpin formation. As G63 and G71 compose part of the ligand binding pocket and are essential for ZMP binding^62^, the G63-G71 distance likely reports on both ligand binding and P4 stability. A caveat here is that the native RNA chain grows during transcription, so the RNA that initially binds to the ligand may be shorter than the RNA that promotes dissociation, for example, by destabilizing P4.

From a computational perspective, we trained our ML model on the dissociation trajectory of one ligand (**compound 26**) and implemented the learned reaction coordinate to accelerate the dissociation of all the other ligands. Here, the successful dissociations beyond **compound 26** demonstrate that our approach was able to construct a transferable model. As far as we know, this is the first application of ML-augmented MD simulation for enhancing the sampling of RNA-ligand interactions where the model can be generalized to study a variety of ligands using state-of-the-art RNA forcefields^69^ with atomistic detail. Such a framework would be broadly useful in both rationalizing complex and nuanced structure-activity relationship trends often seen with RNA-binding ligands as well as designing novel, improved ligands for other therapeutically relevant RNA targets, efforts that are currently ongoing in our labs.

In conclusion, we demonstrated the utility of using computational and experimental tools to study and understand ZTP riboswitch activation by small molecules and how these tools can be used to investigate the mechanism of riboswitch activation. The insights learned from our MD simulations about RNA-ligand interactions and the observed ligand-induced structural differences could be used in future research for the design of ligands that leverage both binding kinetics, G63-G71 distance, and P4 destabilization for the design of inhibitors of *F. ulcerans* ZTP riboswitch, potentially serving as antimicrobial agents. Taken together, this methodology could be applied to prospective studies aimed at harnessing ligand association/dissociation and conformational dynamics in the design of more potent, bioactive ligands for disease-relevant RNAs.

## ACKNOWLEDGMENTS

This research was supported by the Intramural Research Programs of the National Institutes of Health, National Cancer Institute (NCI), Center for Cancer Research, Project BC011585 (to J.S.S.), National Heart, Lung and Blood Institute (NHLBI; to A.R.F), the NIH Intramural AIDS Targeted Antiviral Program (to A.R.F.), and by the National Institute of General Medical Sciences of the National Institutes of Health under Award Number R35GM142719. The content is solely the responsibility of the authors and does not represent the official views of the National Institutes of Health. Isothermal titration calorimetry was performed in the Biophysics Core of the NHLBI. X-ray diffraction data were collected at Beamline 24-ID-C of the Northeastern Collaborative Access Team beamlines, which are funded by the National Institute of General Medical Sciences from the National Institutes of Health (P30 GM124165). This research used resources of the Advanced Photon Source, a U.S. Department of Energy (DOE) Office of Science User Facility operated for the DOE Office of Science by Argonne National Laboratory under Contract No. DE-AC02-06CH11357. P.P. was part of the post-baccalaureate program of NHLBI, and C.P.J. is the recipient of the NIH Transition Award K22HL139920. S.M. thanks the NCI-UMD Partnership for Integrative Cancer Research for the fellowship. We thank UMD HPC’s Zaratan, NSF ACCESS (project CHE180027P), and NIH HPC Biowulf cluster (http://hpc.nih.gov for computational resources. P.T. is an investigator at the University of Maryland-Institute for Health Computing, which is supported by funding from Montgomery County, Maryland and The University of Maryland Strategic Partnership: MPowering the State, a formal collaboration between the University of Maryland, College Park, and the University of Maryland, Baltimore.

## References

1 X. Wu and D. P. Bartel, “Widespread influence of 3’-end structures on mammalian mrna processing and stability,” Cell 169, 905–917 (2017).

2 J. W. Fischer, V. F. Busa, Y. Shao, and A. K. Leung, “Structure-mediated rna decay by upf1 and g3bp1,” Molecular cell 78, 70–84 (2020).

3 J. L. Chen, P. Zhang, M. Abe, H. Aikawa, L. Zhang, A. J. Frank, T. Zembryski, C. Hubbs, H. Park, J. Withka, et al., “Design, optimization, and study of small molecules that target tau premrna and affect splicing,” Journal of the American Chemical Society 142, 8706–8727 (2020).

4 N. A. Naryshkin, M. Weetall, A. Dakka, J. Narasimhan, X. Zhao, Z. Feng, K. K. Ling, G. M. Karp, H. Qi, M. G. Woll, et al., “Smn2 splicing modifiers improve motor function and longevity in mice with spinal muscular atrophy,” science 345, 688–693 (2014).

5 J. Palacino, S. E. Swalley, C. Song, A. K. Cheung, L. Shu, X. Zhang, M. Van Hoosear, Y. Shin, D. N. Chin, C. G. Keller, et al., “Smn2 splice modulators enhance u1–pre-mrna association and rescue sma mice,” Nature chemical biology 11, 511–517 (2015).

6 J. R. Babendure, J. L. Babendure, J.-H. Ding, and R. Y. Tsien, “Control of mammalian translation by mrna structure near caps,” Rna 12, 851–861 (2006).

7 S. Balaratnam, Z. R. Torrey, D. R. Calabrese, M. T. Banco, K. Yazdani, X. Liang, C. R. Fullenkamp, S. Seshadri, R. J. Holewinski, T. Andresson, et al., “Investigating the nras 5’ utr as a target for small molecules,” Cell Chemical Biology (2023).

8 P. B. Kim, J. W. Nelson, and R. R. Breaker, “An ancient riboswitch class in bacteria regulates purine biosynthesis and onecarbon metabolism,” Molecular cell 57, 317–328 (2015).

9 L. Bastet, P. Turcotte, J. T. Wade, and D. A. Lafontaine, “Maestro of regulation: riboswitches orchestrate gene expression at the levels of translation, transcription and mrna decay,” RNA biology 15, 679–682 (2018).

10 W. M. Hewitt, D. R. Calabrese, and J. S. Schneekloth, “Evidence for ligandable sites in structured rna throughout the protein data bank,” Bioorganic & Medicinal Chemistry 27, 2253–2260 (2019).

11 J. A. Howe, H. Wang, T. O. Fischmann, C. J. Balibar, L. Xiao, A. M. Galgoci, J. C. Malinverni, T. Mayhood, A. Villafania, A. Nahvi, N. Murgolo, C. M. Barbieri, P. A. Mann, D. Carr, E. Xia, P. Zuck, D. Riley, R. E. Painter, S. S. Walker, B. Sherborne, R. de Jesus, W. Pan, M. A. Plotkin, J. Wu, D. Rindgen, J. Cummings, C. G. Garlisi, R. Zhang, P. R. Sheth, C. J. Gill, H. Tang, and T. Roemer, “Selective small-molecule inhibition of an rna structural element,” Nature 526, 672–677 (2015).

12 B. P. Charrette, M. A. Boerneke, and T. Hermann, “Ligand optimization by improving shape complementarity at a hepatitis c virus rna target,” ACS Chemical Biology 11, 3263–3267 (2016).

13 C. M. Connelly, T. Numata, R. E. Boer, M. H. Moon, R. S. Sinniah, J. J. Barchi, A.R. Ferré-D’Amaré, and J.S. Schneekloth Jr, “Synthetic ligands for preq1 riboswitches provide struc-tural and mechanistic insights into targeting rna tertiary structure,” Nature communications 10, 1501 (2019).

14 C. Sheridan, “Publisher correction: First small-molecule drug targeting rna gains momentum,” Nature biotechnology 39, 387– 387 (2021).

15 J. L. Childs-Disney, X. Yang, Q. M. Gibaut, Y. Tong, R. T. Batey, and M. D. Disney, “Targeting rna structures with small molecules,” Nature Reviews Drug Discovery 21, 736–762 (2022).

16 C. Connelly, M. Moon, and J. Schneekloth, “The emerging role of rna as a therapeutic target for small molecules,” Cell Chemical Biology 23, 1077–1090 (2016).

17 J. Sponer, G. Bussi, M. Krepl, P. Ban, S. Bottaro, R. A. Cunha, A. Gil-Ley, G. Pinamonti, S. Poblete, P. Jurecka, et al., “Rna structural dynamics as captured by molecular simulations: a comprehensive overview,” Chemical reviews 118, 4177–4338 (2018).

18 M. Bernetti and G. Bussi, “Integrating experimental data with molecular simulations to investigate rna structural dynamics,” Current Opinion in Structural Biology 78, 102503 (2023).

19 A. Cesari, S. Bottaro, K. Lindorff-Larsen, P. Banaš, J. Sponer, and G. Bussi, “Fitting corrections to an rna force field using experimental data,” Journal of chemical theory and computation 15, 3425–3431 (2019).

20 G. Bussi, M. Bonomi, P. Gkeka, M. Sattler, H. M. Al-Hashimi, P. Auffinger, M. Duca, Y. Foricher, D. Incarnato, A. N. Jones, et al., “Rna dynamics from experimental and computational approaches,” arXiv preprint 2404.02789 (2024).

21 F. P. Panei, P. Gkeka, and M. Bonomi, “Identifying small-molecules binding sites in rna conformational ensembles with shaman,” Nature Communications 15, 5725 (2024).

22 E. A. Dethoff, J. Chugh, A. M. Mustoe, and H. M. Al-Hashimi, “Functional complexity and regulation through rna dynamics,” Nature 482, 322–330 (2012).

23 M. H. Moon, T. A. Hilimire, A. M. Sanders, and J. Schneekloth, John S., “Measuring rna–ligand interactions with microscale thermophoresis,” Biochemistry 57, 4638–4643 (2018), doi: 10.1021/acs.biochem.7b01141.

24 S. D. Gilbert and R. T. Batey, “Monitoring rna–ligand interactions using isothermal titration calorimetry,” Riboswitches: Methods and Protocols, 97–114 (2009).

25 J. W. Arney and K. M. Weeks, “Rna–ligand interactions quantified by surface plasmon resonance with reference subtraction,” Biochemistry 61, 1625–1632 (2022).

26 A. Chauvier, P. St-Pierre, J.-F. Nadon, E. D. Hien, C. Pérez-González, S. H. Eschbach, A.-M. Lamontagne, J. C. Penedo, and D. A. Lafontaine, “Monitoring rna dynamics in native transcriptional complexes,” Proceedings of the National Academy of Sciences 118, e2106564118 (2021).

27 C. van der Feltz and A. A. Hoskins, “Methodologies for studying the spliceosome’s rna dynamics with single-molecule fret,” Methods 125, 45–54 (2017).

28 S. Warhaut, K. R. Mertinkus, P. Höllthaler, B. Fürtig, M. Heilemann, M. Hengesbach, and H. Schwalbe, “Ligand-modulated folding of the full-length adenine riboswitch probed by nmr and single-molecule fret spectroscopy,” Nucleic acids research 45, 5512–5522 (2017).

29 B. Hua, C. P. Jones, J. Mitra, P. J. Murray, R. Rosenthal, A.R. Ferré-D’Amaré, and T. Ha, “Real-time monitoring of single ztp riboswitches reveals a complex and kinetically controlled decision landscape,” Nature communications 11, 4531 (2020).

30 K. L. Frieda and S. M. Block, “Direct observation of cotranscriptional folding in an adenine riboswitch,” Science 338, 397–400 (2012).

31 J. Šponer, G. Bussi, M. Krepl, P. Banáš, S. Bottaro, R. A. Cunha, A. Gil-Ley, G. Pinamonti, S. Poblete, P. Jurečka, N. G. Walter, and M. Otyepka, “Rna structural dynamics as captured by molecular simulations: A comprehensive overview,” Chemical Reviews 118, 4177–4338 (2018), doi: 10.1021/acs.chemrev.7b00427.

32 A. C. Stelzer, A. T. Frank, J. D. Kratz, M. D. Swanson, M. J. Gonzalez-Hernandez, J. Lee, I. Andricioaei, D. M. Markovitz, and H. M. Al-Hashimi, “Discovery of selective bioactive small molecules by targeting an rna dynamic ensemble,” Nature chemical biology 7, 553–559 (2011).

33 S. A. Hollingsworth and R. O. Dror, “Molecular dynamics simulation for all,” Neuron 99, 1129–1143 (2018).

34 D. E. Shaw, J. Grossman, J. A. Bank, B. Batson, J. A. Butts, J. C. Chao, M. M. Deneroff, R. O. Dror, A. Even, C. H. Fenton, et al., “Anton 2: raising the bar for performance and programmability in a special-purpose molecular dynamics super-computer,” in SC’14: Proceedings of the International Conference for High Performance Computing, Networking, Storage and Analysis (IEEE, 2014) pp. 41–53.

35 J. D. Owens, M. Houston, D. Luebke, S. Green, J. E. Stone, and J. C. Phillips, “Gpu computing,” Proceedings of the IEEE 96, 879–899 (2008).

36 K. Lindorff-Larsen, S. Piana, R. O. Dror, and D. E. Shaw, “How fast-folding proteins fold,” Science 334, 517–520 (2011).

37 M. Shekhar, Z. Smith, M. A. Seeliger, and P. Tiwary, “Protein flexibility and dissociation pathway differentiation can explain onset of resistance mutations in kinases,” Angewandte Chemie International Edition 61, e202200983 (2022).

38 S. Mehdi, Z. Smith, L. Herron, Z. Zou, and P. Tiwary, “Enhanced sampling with machine learning,” Annual Review of Physical Chemistry 75 (2024).

39 M. Batool, B. Ahmad, and S. Choi, “A structure-based drug discovery paradigm,” International journal of molecular sciences 20, 2783 (2019).

40 L. W. Hardy and A. Malikayil, “The impact of structure-guided drug design on clinical agents,” Curr. Drug Discov 3, 15–20 (2003).

41 F. P. Panei, R. Torchet, H. Menager, P. Gkeka, and M. Bonomi, “Hariboss: a curated database of rna-small molecules structures to aid rational drug design,” Bioinformatics 38, 4185–4193 (2022).

42 J. Manigrasso, M. Marcia, and M. De Vivo, “Computer-aided design of rna-targeted small molecules: a growing need in drug discovery,” Chem 7, 2965–2988 (2021).

43 Q. Vicens, E. Mondragón, F. E. Reyes, P. Coish, P. Aristoff, J. Berman, H. Kaur, K. W. Kells, P. Wickens, J. Wilson, et al., “Structure–activity relationship of flavin analogues that target the flavin mononucleotide riboswitch,” ACS chemical biology 13, 2908–2919 (2018).

44 S. E. Motika, R. J. Ulrich, E. J. Geddes, H. Y. Lee, G. W. Lau, and P. J. Hergenrother, “Gram-negative antibiotic active through inhibition of an essential riboswitch,” Journal of the American Chemical Society 142, 10856–10862 (2020).

45 M. Mandal and R. R. Breaker, “Gene regulation by riboswitches,” Nature Reviews Molecular Cell Biology 5, 451–463 (2004).

46 K. F. Blount and R. R. Breaker, “Riboswitches as antibacterial drug targets,” Nature biotechnology 24, 1558–1564 (2006).

47 S. Mehdi, C. R. Fullenkamp, C. P. Jones, P. Tiwary, and J. S. Schneekloth, “Insights into the discrepancy between affinity and activation in f. ulcerans ztp riboswitch activators through structure-informed design and machine learning-augmented molecular dynamics simulations,” Biophysical Journal 123, 454a (2024).

48 B. Tran, P. Pichling, L. Tenney, C. M. Connelly, M. H. Moon, A.R. Ferré-D’Amaré, J. S. Schneekloth, and C. P. Jones, “Parallel discovery strategies provide a basis for riboswitch ligand design,” Cell chemical biology 27, 1241–1249 (2020).

49 O. Livi, P. Ferrarini, I. Tonetti, F. Smaldone, and G. Zefola, “Synthesis and pharmacological activity of 1, 2, 3-triazole derivatives of naphthalene, quinoline and pyridine,” Il Farmaco; Edizione Scientifica 34, 217–228 (1979).

50 A. Das, “Studies on complex π-π and t-stacking features of imidazole and phenyl/p-halophenyl units in series of 5-amino-1-(phenyl/p-halophenyl) imidazole-4-carboxamides and their carbonitrile derivatives: Role of halogens in tuning of conformation,” Journal of Molecular Structure 1147, 520–540 (2017).

51 I. Artsimovitch and T. M. Henkin, “In vitro approaches to analysis of transcription termination,” Methods 47, 37–43 (2009).

52 T. Vo, A. Paul, A. Kumar, D. W. Boykin, and W. D. Wilson, “Biosensor-surface plasmon resonance: A strategy to help establish a new generation rna-specific small molecules,” Methods 167, 15–27 (2019).

53 J. J. Trausch, J.G. Marcano-Velázquez, M. M. Matyjasik, and R. T. Batey, “Metal ion-mediated nucleobase recognition by the ztp riboswitch,” Chemistry & biology 22, 829–837 (2015).

54 E. R. Beyerle, S. Mehdi, and P. Tiwary, “Quantifying energetic and entropic pathways in molecular systems,” The Journal of Physical Chemistry B 126, 3950–3960 (2022).

55 D. Wang and P. Tiwary, “State predictive information bottleneck,” The Journal of Chemical Physics 154 (2021).

56 D. P. Kingma and M. Welling, “Auto-encoding variational bayes,” arXiv preprint 1312.6114 (2013).

57 P. Tiwary and M. Parrinello, “From metadynamics to dynamics,” Physical review letters 111, 230602 (2013).

58 P. Tiwary and M. Parrinello, “A time-independent free energy estimator for metadynamics,” The Journal of Physical Chemistry B 119, 736–742 (2015).

59 J. M. L. Ribeiro, P. Bravo, Y. Wang, and P. Tiwary, “Reweighted autoencoded variational bayes for enhanced sampling (rave),” The Journal of chemical physics 149 (2018).

60 S. Mehdi, D. Wang, S. Pant, and P. Tiwary, “Accelerating all-atom simulations and gaining mechanistic understanding of biophysical systems through state predictive information bottleneck,” Journal of chemical theory and computation 18, 3231– 3238 (2022).

61 Y. Wang, S. Parmar, J. S. Schneekloth, and P. Tiwary, “Interrogating rna–small molecule interactions with structure probing and artificial intelligence-augmented molecular simulations,” ACS Central Science 8, 741–748 (2022).

62 C. P. Jones and A.R. Ferré-D’Amaré, “Recognition of the bacterial alarmone zmp through long-distance association of two rna subdomains,” Nature structural & molecular biology 22, 679–685 (2015).

63 A. Ren, K. R. Rajashankar, and D. J. Patel, “Global rna fold and molecular recognition for a pfl riboswitch bound to zmp, a master regulator of one-carbon metabolism,” Structure 23, 1375– 1381 (2015).

64 K. F. Blount, C. Megyola, M. Plummer, D. Osterman, T. O’Connell, P. Aristoff, C. Quinn, R. A. Chrusciel, T. J. Poel, H. J. Schostarez, et al., “Novel riboswitch-binding flavin analog that protects mice against clostridium difficile infection without inhibiting cecal flora,” Antimicrobial agents and chemotherapy 59, 5736–5746 (2015).

65 J. K. Wickiser, M. T. Cheah, R. R. Breaker, and D. M. Crothers, “The kinetics of ligand binding by an adenine-sensing riboswitch,” Biochemistry 44, 13404–13414 (2005).

66 C. Helmling, D.-P. Klötzner, F. Sochor, R. A. Mooney, A. Wacker, R. Landick, B. Fürtig, A. Heckel, and H. Schwalbe, “Life times of metastable states guide regulatory signaling in transcriptional riboswitches,” Nature Communications 9, 944 (2018).

67 E. J. Strobel, L. Cheng, K. E. Berman, P. D. Carlson, and J. B. Lucks, “A ligand-gated strand displacement mechanism for ztp riboswitch transcription control,” Nature chemical biology 15, 1067–1076 (2019).

68 H. Yu-Nan, W. Kang, S. Yu, X. Xiao-Jun, W. Yan, L. Xing-Ao, and S. Ting-Ting, “Molecular dynamics simulation on the thermosinus carboxydivorans pfl ztp riboswitch by ligand binding,” Biochemical and Biophysical Research Communications 627, 184–190 (2022).

69 M. R. Tucker, S. Piana, D. Tan, M. V. LeVine, and D. E. Shaw, “Development of force field parameters for the simulation of single-and double-stranded dna molecules and dna–protein complexes,” The Journal of Physical Chemistry B 126, 4442–4457 (2022).

