## Supplementary Information for "Machine learning-augmented molecular dynamics simulations (MD) reveal insights into the disconnect between affinity and activation of ZTP riboswitch ligands"

### I. BIOPHYSICAL AND *IN VITRO* METHODS AND MATERIALS

#### A. RNA and DNA Sequences:

#### B. Crystallization, model building, and refinement

The 64-nt *S. ondontolytica* RNA was transcribed from a template consisting of a 5'-hammerhead ribozyme [1] and the ZTP riboswitch cDNA (**Table S1**, S.ondontolytica ZTP Riboswitch used for crystallography). A 2'-O-methylated reverse primer was used during PCR-amplification of the transcription template to ensure homogeneity, as previously described [2]. Thus, the 64-nt RNA used herein lacks the 5'-phosphate present in an earlier study[3]. The RNA was folded in a buffer containing 25 mM HEPES-KOH, pH 7.4, 150 mM KCl, and 3-fold excess ligand (either **1** or **23**) by heating for 95 ℃ for 2 min and immediately placed on ice. $MgCl_2$ was added to 10 mM final, and RNA was incubated for 15 min at 37 ℃ and stored at room temperature until setting up hanging drop crystal trays using the vapor diffusion method.

For **1**, 6 g/l RNA was mixed with a reservoir solution of 27-31% (v/v) polyethylene glycol (PEG) 4000, 0.1 M sodium acetate, pH 4.5, and 0.2 M ammonium acetate. Drops consisted of 0.7 uL folded RNA and 1.4 uL reservoir solution, 1 uL of each, or 1.4 uL folded RNA and 0.7 uL reservoir solution. Hanging drops were equilibrated against 0.5 mL reservoir solution in 15-well EasyXtal trays (NeXtal) at 21 ℃. Crystals were observed after 1 month. Crystals were cryoprotected in a solution containing the RNA folding buffer, crystal reservoir, 24.8% (v/v) PEG 4000, and 10% (v/v) ethylene glycol and were vitrified via flash plunging into liquid nitrogen.

For **23**, 4 g/l RNA was mixed with a reservoir solution of 125-300 mM LiSO4, 23-27% PEG 3350. Hanging drops consisted of 0.2 uL reservoir solutioin and 0.2 uL folded RNA and were equilibrated at 21 ℃ in 96-well plates. Crystals appeared after 4-5 days. Crystals were cryoprotected in a solution containing the RNA folding buffer and reservoir solution except that PEG 3350 was increased to 30% (v/v) and that 5% (v/v) ethylene glycol was added. Crystals were flash plunged into liquid nitrogen.

Both datasets were phased via molecular replacement using PHASER [4] with a search model of the *S. odontolytica* RNA (PDB: 4XWF) with ligands and ions removed. For **1**, molecular replacement resulted in a translation-factor Z-score (TFZ) of 28.1 and log-

**Table S1. Oligonucleotide sequences**

| Oligonucleotides: | Sequence: |
|---|---|
| *F. ulcerans* ZTP riboswitch ITC construct RNA | UAUCAGUUAUAUGACUGACGGAACGUGGAAUUAACCACAUGAAGUAUAACG<br>AUGACAAUGCCGACCGUCUGGGCG |
| *F. ulcerans* ZTP riboswitch ITC construct forward primer | TAATACGACTCACTATAGG |
| *F. ulcerans* ZTP riboswitch ITC construct reverse primer | mCmGCCCAGACGGTCGGCAT |
| *F. ulcerans* ZTP riboswitch SPR construct RNA | UAUCAGUUAUAUGACUGACGGAACGUGGAAUUAACCACAUGAAGUAUAACG<br>AUGACAAUGCCGACCGUCUGGGCGUUCAUUAGCGGUAUUCCGGAAC |
| *F. ulcerans* ZTP riboswitch SPR construct PCR RP | mGmUTCCGGAATACCGCTAATGAACGCCCAGACGGTCGGCATTG |
| *F. ulcerans* ZTP riboswitch SPR Non-binding mutant (U70C) construct RNA | UAUCAGUUAUAUGACUGACGGAACGUGGAAUUAACCACAUGAAGUAUAACG<br>AUGACAAUGCCGACCGUC**C**GGGCGUUCAUUAGCGGUAUUCCGGAAC |
| *F. ulcerans* ZTP riboswitch SPR Non-binding mutant (U70C) construct PCR RP | mGmUTCCGGAATACCGCTAATGAACGCCCGGACGGTCGGCATTG |
| *F. ulcerans* ZTP riboswitch SPR biotinylated-DNA hybridization anchor | 5′-/Biotin/-GTTCCGGAATACCGCTAATG |
| *F. ulcerans* ZTP riboswitch single-round transcription termination template forward primer | CTTGATTCTAGCTGATCGTGGACC |
| *F. ulcerans* ZTP riboswitch single-round transcription termination template reverse primer | TCAACAATTATATTTGATATTCCC |
| *F. ulcerans* ZTP riboswitch single-round transcription termination template in PCR2.1 plasmid | CTTGATTCTAGCTGATCGTGGACCGGAAGGTGAGCCAGTGAGTTGATTGCA<br>GTCCAGTTACGCTGGAGTCTGAGGCTCGTCCTGAATGATATGCGGCCTCTT<br>GACTATTTTACCTCTGGCGGTGATAATGGTTGCAATGTAGTAAGGAGGTTG<br>TATGGAAGATTATCAGTTATATGACTGA<br>CGGAACGTGGAATTAACCACATGAAGTATAACGATGACAATGCCGACCGTC<br>TGGGCGAACAAGCCTAGATTGTCGGTTTTTTTTATACATTTTTTTAGGAGG<br>AGATAATAAGGGAATATCAAATATAATTGTTGA |
| *S. ondontolytica* ZTP Riboswitch used for crystallography | GGGUCGUGACUGGCGAACAGGUGGGAAACCACCGGGGAGC<br>GACCCUUGCCGCCCGCCUGGGCAA |

likelihood gain (LLG) of 1251. For **23**, molecular replacement resulted in TFZ and LLG and

42.7 and 4333, respectively. Refinement was performed in Phenix[5]. Initial refinements used individual B-factor refinement, real space, and simulated annealing, and final refinements implemented TLS refinement. For **1**, which crystallized in the same space group as the previously described ZMP-bound RNA[3], model building was not necessary beyond ligand placement and addition of waters and ions. Data were initially refined using the entire resolution range (to 2.25 Å) and later limited to 2.43 Å. For **23**, a conformational change was evident upon inspection, and residues 16-19 and C52 were rebuilt. Weak density present at the former position of C52 suggested the possible presence of an ion (e.g., hydrated magnesium), which was not built. Two weaker copies of the RNA were partially visible but could not be built. Reindexing and solving in lower symmetry space groups (P *6*, P *3 1 2*, and P *3 2 1*), assuming twinning, did not improve the quality of weak density. Data were refined initially to 2.7 Å later to 2.15 Å.

**Isothermal Titration Calorimetry (ITC)**

ITC measurements were performed using a Malvern MicroCal iTC200, and data were fit using SEDPHAT and NITIPIC as previously described [6, 7]. All experiments were performed in 50 mM Hepes-KOH, pH 7.4, 150 mM KCl, and 10 mM $MgCl_2$ and up to 3% DMSO at 37 ℃ in triplicate with ZTP riboswitch sequence indicated in (**Table S1**, *F. ulcerans* ZTP riboswitch ITC construct RNA).

**Single-Round Transcription Termination Assays**

Single-round Transcription Termination assays were performed as previously described [7]. Samples were separated using 8% denaturing PAGE and imaged by exposing the gel to a storage phosphor screen overnight. Band intensities were measured using ImageQuant software. The percentage of readthrough was calculated after background correcting all bands and dividing the intensity of the full-length readthrough band by the total intensity of terminated plus readthrough bands. These values were fitted to a simple binding isotherm assuming a 1:1 binding stoichiometry in OriginLab 8.0 to determine the $T_{50}$ value.

**Table S2**. Summary of crystallographic statistics

| | **B28** | **A012** |
|---|---|---|
| **Data collection** | | |
| Space group | $P4_32_12$ | P622 |
| Cell dimensions | | |
| a, b, c (Å) | 41.035, 41.035, 237.398 | 118.247, 118.247, 202.054 |
| a, b, g (°) | 90, 90, 90 | 90, 90, 120 |
| Resolution (Å) | 40.44 - 2.25 | 102.40 - 1.98 |
| $R_{merge}$ | 0.09664 (0.8578)* | 0.2546 (2.508) |
| CC1/2 | 0.998 (0.854) | 0.997 (0.831) |
| <I> / <s(I)> | 16.27 (1.89) | 12.41 (2.34) |
| Completeness (%) | 99.63 (98.99) | 99.89 (100.00) |
| Redundancy | 12.0 (13.4) | 39.1 (38.8) |
| **Refinement** | | |
| Resolution (Å) | 40.44 - 2.43 (2.517 - 2.43) | 101 - 2.15 (2.227 - 2.15) |
| No. unique reflections | 8378 (788) | 46116 (4513) |
| $R_{work}$ / $R_{free}$ (%) | 0.2072 (0.4939) / 0.2563 (0.5649) | 0.3097 (0.3533) / 0.3338 (0.3695) |
| No. atoms | | |
| RNA | 1376 | 1376 |
| Ligand | 23 | 28 |
| Ions/water | 17 | 70 |
| Mean B-factors (Å$^2$) | | |
| RNA | 72.53 | 47.08 |
| Ligand | 61.74 | 27.62 |
| Ions/water | 59.66 | 37.55 |
| r.m.s. deviations | | |
| Bond lengths (Å) | 0.005 | 0.005 |
| Bond angles (°) | 1.10 | 0.92 |
| Mean coordinate precision (Å) | 0.49 | 0.42 |
| PDB code | 8VQV | 8VVJ |

*Values in parentheses correspond to highest resolution shells

##### Surface Plasmon resonance (SPR)

##### Streptavidin Reference Channel SPR Surface generation

SPR binding assays were performed on a Biacore 3000 instrument from GE Healthcare in a similar manner as previously described[8]. A CM5 sensor chip was docked and primed with running buffer (50 mM Hepes-KOH, pH 7.4, 150 mM KCl, 10 mM $MgCl_2$, 0.005% tween-20, 5% DMSO) before surface generation. For surface generation, flow cells 1 (FC1) and 2 (FC2) were activated by injecting an aqueous solution of EDS/NHS (0.4M/0.1M) for

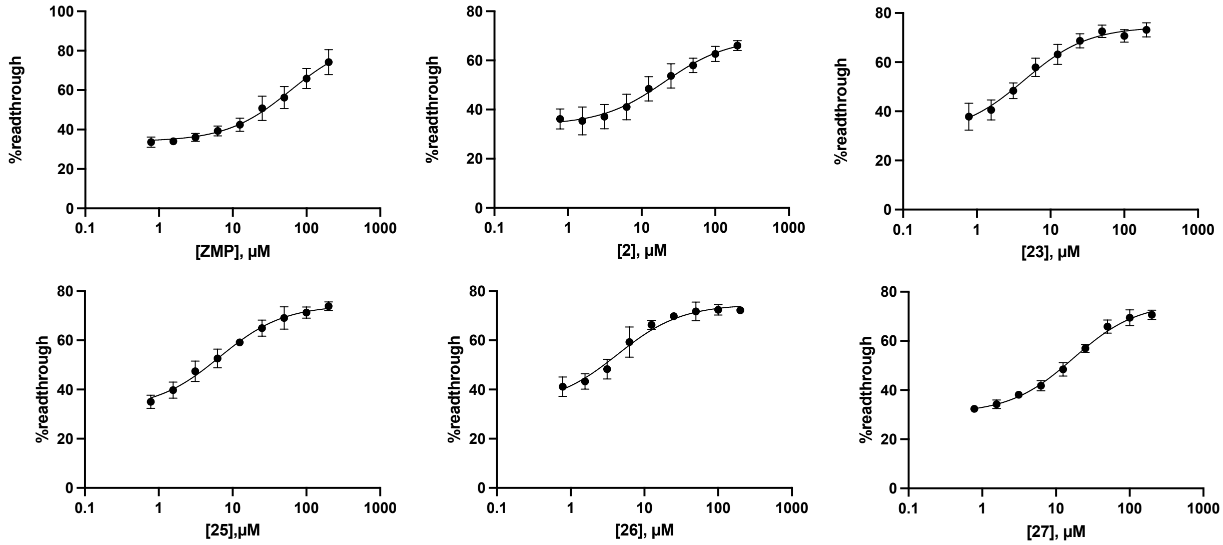

**Supporting Figure 1: Single-round transcription termination assay.** Transcription termination assay for **ZMP**, **2, 23, 25, 26,** and **27.** (n=3, error bars denote standard deviations).

15 min with a flow rate of 5 $\mu$L/min. Following surface activation, a solution of streptavidin (0.2 mg/mL in 10 mM sodium acetate buffer, pH 4.5) was injected for 30 min with a flow rate of 5 $\mu$L/min, to reach an immobilization of approximately 6,000-8,000 RU. Following streptavidin immobilization, the surface was deactivated by injecting a 1 M aqueous solution of ethanolamine (pH 8.5) for 10 min, followed by injection of a 10 mM aqueous solution of NaOH for 2 min to remove any unbound streptavidin.

After surface generation, F. ulcerans ZTP RNA (**Table S1**, *F. ulcerans* ZTP riboswitch SPR construct RNA) and biotin-labeled hybridization DNA anchor (**Table S1**, F. ulcerans ZTP riboswitch SPR biotinylated-DNA hybridization anchor) were diluted and combined in 50 mM Hepes-KOH pH 7.4, 150 mM KCl, 10 mM $MgCl_2$, 0.005% tween-20 and 5% DMSO to a final concentration of 5 $\mu$M RNA and 2.5 $\mu$M DNA. The diluted RNA/DNA solution was hybridized by incubation in a heat block at 95 ℃ for 5 min and then removed and allowed to slowly cool down to room temperature for one hour. After cooling, the RNA-DNA duplex was injected onto FC2 at a flow rate of 5$\mu$L/min for 30 min.

**Non-binding RNA mutant Reference channel SPR surface generation**

The same surface activation was completed for the non-binding RNA mutant reference channel experiments, as described in the previous section. The method differs from the above protocol after the surface generation step. As before, F. ulcerans ZTP RNA (**Table S1**, *F. ulcerans* ZTP riboswitch SPR construct RNA) and biotin-labeled hybridization DNA anchor (**Table S1**, F. ulcerans ZTP riboswitch SPR biotinylated-DNA hybridization anchor) were diluted and combined in 50 mM Hepes-KOH pH 7.4, 150 mM KCl, 10 mM $MgCl_2$, 0.005% tween-20 and 5% DMSO to a final concentration of 5 $\mu$M RNA and 2.5 $\mu$M DNA. The diluted RNA/DNA solution was hybridized by incubation in a heat block at 95 ℃ for 5 min and then removed and allowed to cool down to room temperature for one hour. At the same time, a mutated non-binding ZTP RNA aptamer (U70C mutant, **Table S1**, *F. ulcerans* ZTP riboswitch SPR Non-binding mutant (U70C) construct RNA) and biotin-labeled hybridization DNA anchor (**Table S1**, F. ulcerans ZTP riboswitch SPR biotinylated-DNA hybridization anchor), were diluted and combined in 50 mM Hepes-KOH pH 7.4, 150 mM KCl, 10 mM $MgCl_2$, 0.005% tween-20 and 5% DMSO to a final concentration of 5 $\mu$M RNA and 2.5 $\mu$M DNA. The diluted RNA/DNA solution was hybridized by incubation in a heat block at 95 ℃ for 5 min and then removed and allowed to cool down to room temperature slowly. After cooling, the RNA-DNA duplex (binding variant) was injected onto FC2 at a flow rate of 5 $\mu$L/min for 30 min, followed by injection of the non-binding RNA-DNA duplex onto FC1 (reference channel) for 30 min at a flow rate of 5 $\mu$L/min.

For binding analysis, a 10 mM DMSO stock of each ligand was diluted into running buffer (50 mM Hepes-KOH pH 7.4, 150 mM KCl, 10 mM $MgCl_2$, 0.005% tween-20), resulting in a final DMSO concentration of 5%. Ligands were serially diluted (1:1) from 10 $\mu$M, with each titration series consisting of 8-10 points. Ligands were injected onto FC1 and Fc2 for 2 min at a flow rate of 30 $\mu$L/min, followed by 300s dissociation. If necessary, an injection of 1M KCl (50 $\mu$L) could be performed between samples. The final binding signal curve was obtained by reference channel subtraction, and the $K_D$ was calculated by plotting SPR response versus concentration in GraphPad Prism version 10.1 and fit to a single site binding model.

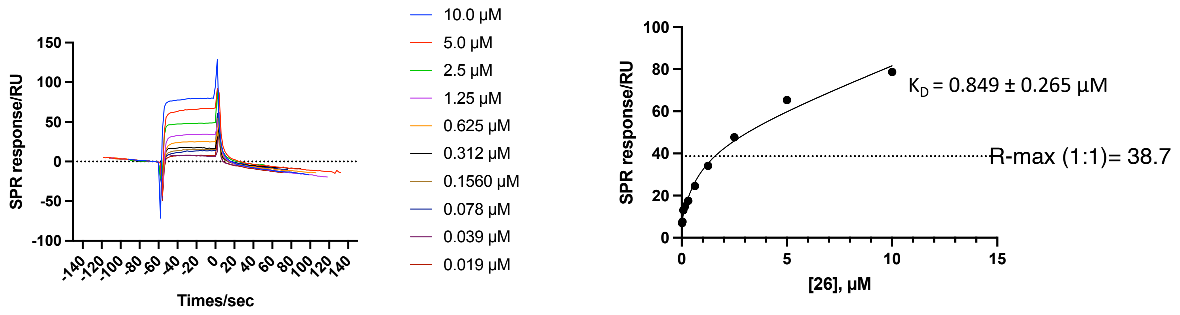

**b)** Non-binding Mutant Subtracted SPR Titration for **26**

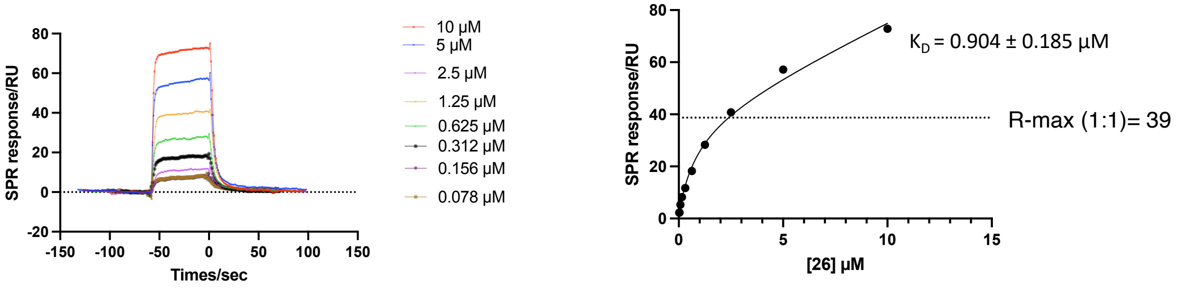

**Supporting Figure 2: Surface Plasmon Resonance titration of 26. a)** SPR titration of **26** with streptavidin reference channel. Titration did not saturate and surpassed 1:1 binding (R-max = 38.7). In addition, kinetic analysis can not be performed due to the steep slope of association and dissociation. **b)** SPR titration of **26** with a mutant ZTP riboswitch (does not bind to ZMP) reference channel. As with, streptavidin, titration did not saturate and surpassed 1:1 binding (R-max = 39). In addition, kinetic analysis could not be performed due to the steep slope of the SPR response curve.

#### II. MOLECULAR DYNAMICS

##### A. Machine-learning informed Molecular Dynamics Simulations

In this study, we adopt well-tempered metadynamics[9, 10] (WT-MetaD) for accelerating molecular dynamics (MD) simulations. An effective implementation of WT-MetaD requires the knowledge of reaction coordinates (RCs) that can distinguish between different metastable states constituting the rare event of interest. Mathematically, if $\mathbf{R}$ denotes the position coordinates of a system and $s(\mathbf{R})$ represents the RCs of the system at time $t$, then WT-MetaD allows the calculation of the average of an equilibrium $\mathbf{R}$ dependent operator $\langle O(\mathbf{R})\rangle_{eq}$ as shown in **Eq. 1**. Here, $O(\mathbf{R},t)$, $\beta$, $V(s(\mathbf{R},t))$, and $c(t)$ denote the position-dependent operator from biased WT-MetaD simulation at time $t$, inverse temperature, history-dependent bias deposited at time $t$, and position independent component of the bias respectively. Additionally, when adopting infrequent WT-MetaD, the transition time ($t$) to go from one metastable state (A) to (B) can be estimated from the acceleration factor, $\alpha(t) = \langle e^{\beta(V(s(\mathbf{R},t)))}\rangle_{s\in A}$.

$$\langle O(\mathbf{R})\rangle_{eq} = \langle O(\mathbf{R},t)e^{\beta(V(s(\mathbf{R},t))-c(t))}\rangle \tag{1}$$

The key challenge here is the knowledge of RCs ($s(\mathbf{R},t)$), which are typically unknown *a priori* for complex events involving biophysical systems. In our work, we address this issue by employing state predictive information bottleneck (SPIB), a machine learning (ML) method for learning RCs, which we discuss in detail below.

**State predictive information bottleneck (SPIB)**

SPIB[11] constructs an information bottleneck (IB) following Shannon's rate-distortion theory[12] to learn a physically meaningful low-dimensional representation ($\mathbf{s}$) of the high-dimensional MD time-series data ($\mathbf{X}$). In contrast to the previously developed RAVE,[13] SPIB directly predicts the metastable state ($\mathbf{y}$) of a system after a time delay ($\Delta t$) by maximizing the simplified objective function, $\mathcal{L}$ shown in **Eq.** 2. Here, the key idea is that the low-dimensional embedding ($\mathbf{s}$) should be as predictive as possible about the time-delayed metastable state ($\mathbf{y}_{\Delta t}$) determined by the mutual information, $I(\mathbf{s},\mathbf{y}_{\Delta t})$. At the same time, $\mathcal{L}$ uses minimal information from the input ($\mathbf{X}$) for the construction of $\mathbf{s}$ to ensure the exclusion of irrelevant information from the low dimensional embedding, determined by the mutual information, $I(\mathbf{X},\mathbf{s})$. The trade-off between these two competing terms is tuned by the hyperparameter $\beta$.

$$\mathcal{L} = I(\mathbf{s},\mathbf{y}_{\Delta t}) - \beta I(\mathbf{X},\mathbf{s}) \tag{2}$$

In practice, SPIB employs a variational autoencoder[14] (VAE) consisting of an encoder-decoder pair to implement the IB. Once the ML model is trained, only the encoder is used to learn on-the-fly low-dimensional representation as a function of the inputs during simulation, which can then be used as the reaction coordinate of WT-MetaD for biasing.

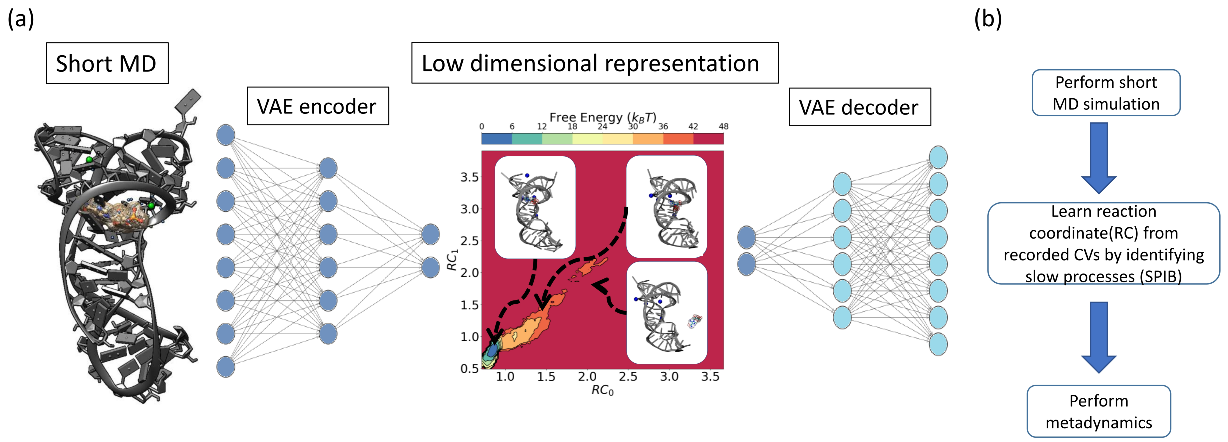

**Supporting Figure 3: Overview of SPIB augmented MD.** A short MD with aggressive WT-MetaD using simple RCs is performed to observe dissociation. Afterward, the recorded CVs are passed through the VAE architecture of SPIB to learn a low-dimensional representation, which serves as the RC of subsequent rounds of WT-MetaD. The learned representation is meaningful as different points on this space correspond to different stages of the ligand dissociation process, with the bound state having the lowest free energy calculated using **Eq.** 1.

SPIB iteratively improves and learns the metastable state definitions from an initial state assignment from the user. The initial state assignment can be made from simple *a priori* information about the system. Since this work involves the dissociation of small molecules from the riboswitch, the initial states for all models were assigned based on the distance between the $Mg^{2+}$ ion inside the cavity and the center of mass (COM) of the polar group of the respective small molecules.

##### System setup and CV details

We employed the experimentally solved crystal structure (6WZS) of the ZTP riboswitch bound to **1**[7] as the initial structure for all simulations. This structure has 15 missing residues (1-5, 48-54, 57-59) which we modeled using ChimeraX.[15] Afterwards, we obtained bound structures for all the other ligands by swapping out **1** and using template-based molecular docking within Molsoft ICM-Pro.

The very recent DES-AMBER forcefield[16] was used to parametrize the RNA, and the small molecules were parametrized using GAFF2.[17] Here, we note a key aspect of the DES-

AMBER forcefield, which rescales partial charges of non-neutralized atom groups by a factor of 0.9. The work[16] by Piana *et al.* highlights the important role of divalent cations (e.g., $Mg^{2+}$) in providing structural stability to RNAs and reports that this rescaling scheme leads to much better agreement with experimental results. The six synthetic derivatives (**compounds 2, 23, 25, 26, 27**) studied in this work are charge neutral, and the results from GAFF2 were kept unchanged. Since the phosphate group of ZMP loses two protons in is a dianion at physiological conditions, we modeled ZMP accordingly, and after generating the GAFF2 forcefield, we rescaled the partial charge of the phosphate group by a factor of 0.9 to be compatible with the DES-AMBER forcefield. We took a similar approach when parametrizing the only positively charged derivative (**4-piperidinyl AICA**). The bound structure was then solvated using $30,234$ TIP4P[18] water molecules and neutralized using 64 $Na^+$ ions. Additionally, $Na^+$ and $Cl^-$ ions were added to maintain a 0.15 M concentration to mimic the physiological conditions of the experiments. We performed all the simulations at 303.15 K temperature and 1 atom pressure using the Nose-Hoover thermostat[19] and Parinello-Rahman barostat[20], respectively.

Since we are studying a dissociative process, a natural choice of collective variables (CVs) for this system is the pairwise distances between the ligand and the RNA residues. Here, we define three regions of interest for each of the ligands. The polar group region ($l_1$) that electrostatically interacts with the $Mg^{2+}$ inside the cavity, the core region ($l_2$) stacking on $RNA_{71}$ residue, the phosphate group for the cognate or the outermost aromatic ring region for the synthetic ligands ($l_3$). By calculating the distances between the center of masses (COMs) of $l_1, l_2, l_3$, and all the 75 residues of the RNA and the $Mg^{2+}$ inside the cavity, we get 228 pairwise distances as CVs. We also monitor $\alpha, \beta, \gamma, \delta, \zeta, \epsilon, \chi$ torsion angles for all the residues. To investigate the role of water and ions in the dissociation mechanism, we monitor water and ion coordination number of the $Mg^{2+}$ inside the cavity and $l_1, l_2, l_3$ regions of the ligands.

###### SPIB augmented MD

An accurate study of small molecule binding events is very challenging through MD as the ligand needs to explore a very high dimensional configuration space and enter the binding cavity. Hence, in this research, we aim to understand the interaction mechanisms between

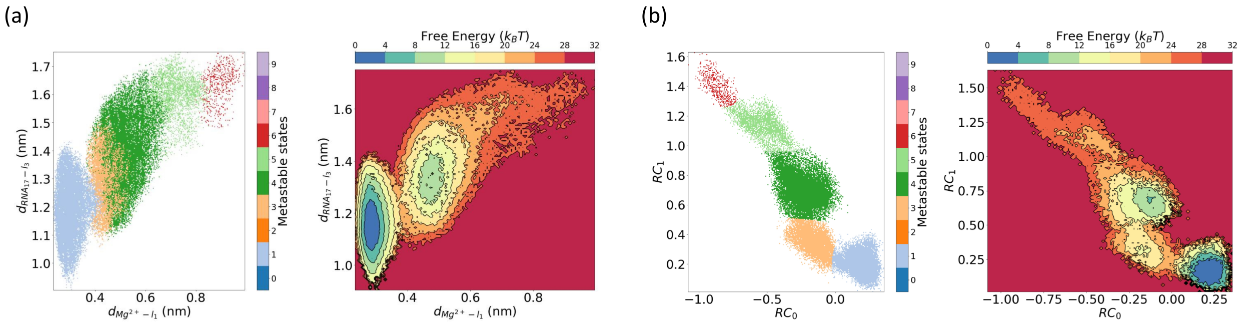

**Supporting Figure 4: Converged SPIB results after training.** (a) The SPIB learned metastable states are projected along the expertise-based RCs using different colors in the left plot. As the ligand exits the cavity, both $d_{Mg^{2+}-l_1}$ and $d_{RNA_{17}-l_3}$ increase, and SPIB detects different metastable states across the dissociation path. The right plot shows the free energy projected along these distances calculated from the logarithm of their Boltzmann weighted probabilities according to **Eq.** 1. (b) Metastable state projections

ZTP riboswitch and the ligands by simulating the dissociation process.

We summarize our protocol in **Fig.** 3. After performing energy minimization and equilibration, we perform 50 ns unbiased simulation and record the standard deviation of all the CVs for only one of the ligand-RNA complexes (**26**). Afterward, WT-MetaD is implemented for this ligand to observe dissociation by biasing $d_{MG^{2+}-l_1}$, and $d_{RNA_{17}-l3}$. We took 75% of the standard deviation of the unbiased run as WT-Metad $\sigma$, height = 1.5 KJ/mol, bias factor = 40, pace = 2.0 ps. Without using such a high $\sigma$ we were unable to observe dissociation in this round.

We learn an improved RC from this simulation data through SPIB in three steps. Here, we note that there are two classes of pairwise distances that can help to distinguish bound-unbound states. Pairwise distances where (1) RNA residues need to undergo a conformational change for the ligand to dissociate, (2) RNA residues that don't undergo significant structural change, but the pairwise distances change because of their relative position along the dissociation path. For an effective implementation of WT-MetaD, we use SPIB to perform feature selection in the first two steps by identifying RNA residues with the highest changes in torsion angles, and regions of the ligand that are most informative when computing pairwise distances, respectively. In the third step, we used the top 16 most informative pairwise distances to construct the low-dimensional representation using a linear encoder,

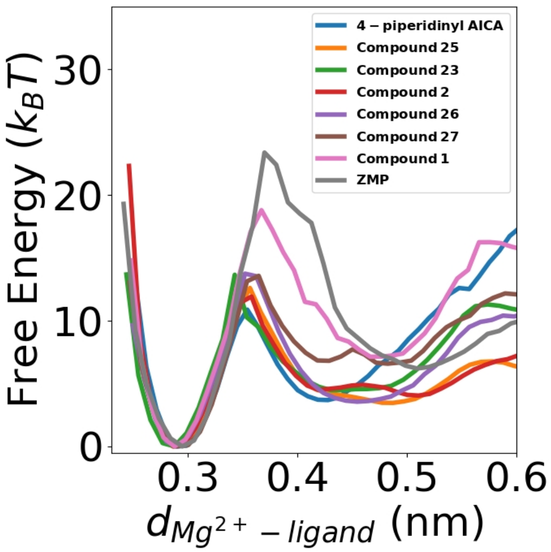

**Supporting Figure 5: One-dimensional free energy profile during dissociation.** Free energy projected along the distance between $MG^{2+}$ and computationally studied ligands showing a high free energy barrier for **ZMP, and compound 1**.

which we biased using WT-MetaD. During ML model training, all the distances were normalized using min-max scaling, and sines and cosines of all the torsion angles were used. The SPIB learned RC is then biased using 25% of their standard deviation as $\sigma$ and keeping all the other WT-MetaD hyperparameters the same as above. Finally, the learned model for synthetic compound **26** was transferred to other small molecules, and WT-MetaD was implemented using the same hyperparameters. Simulations of each ligand were performed in 16 individual replicas to observe 16 separate dissociation events for calculation of all the CVs.

#### III. CHEMISTRY

##### A. General Procedure:

All non-aqueous reactions were performed in flame-dried or oven-dried round-bottomed flasks under an argon atmosphere. Stainless steel syringes or cannula were used to transfer air- and moisture-sensitive liquids. Reaction temperatures were controlled using a thermocouple thermometer and analog hotplate stirrer and monitored using liquid-in-glass thermometers. Unless otherwise noted, reactions were conducted at room temperature (approximately 21-23 °C). Flash column chromatography was performed using Teledyne CombiFlash

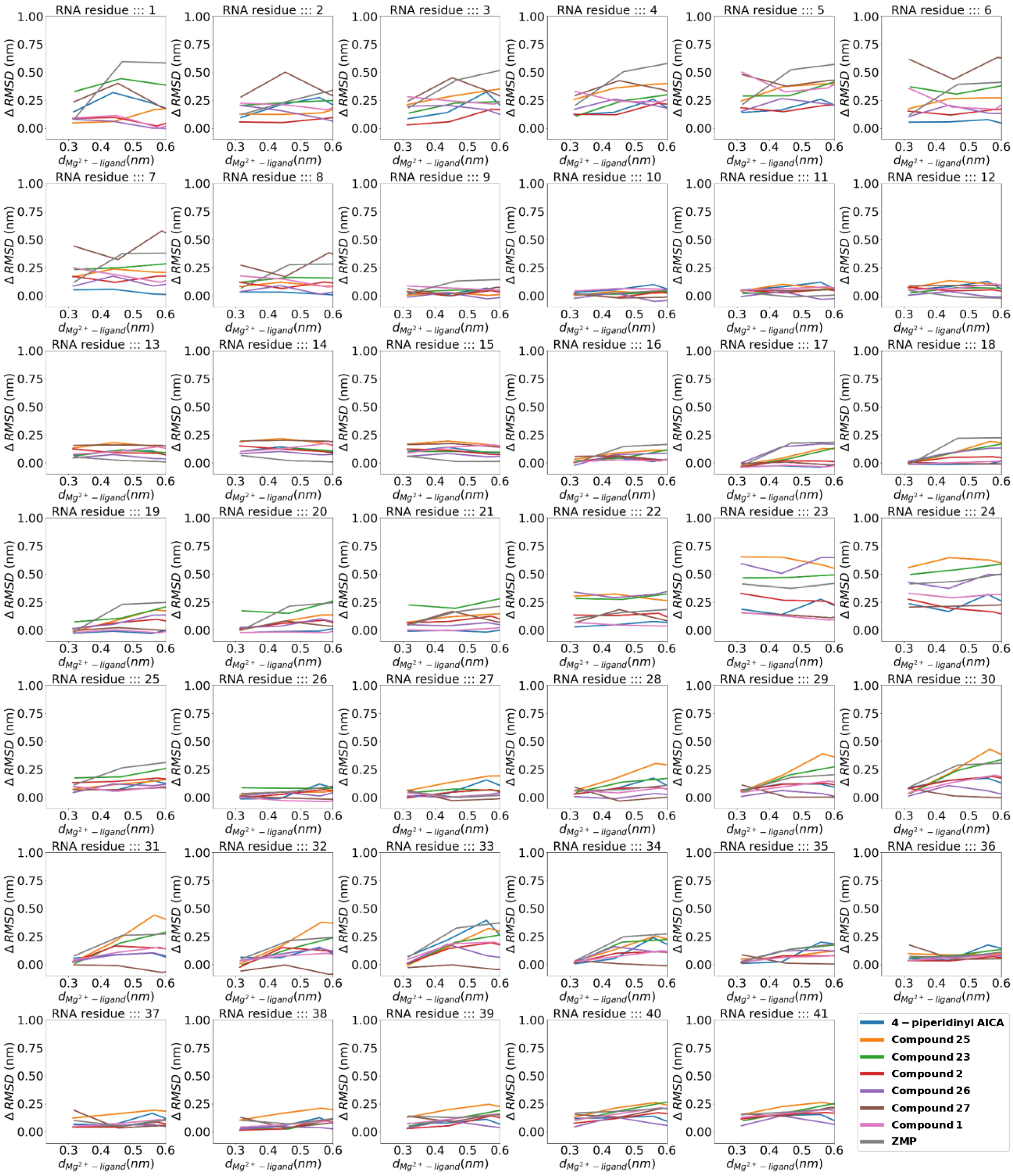

**Supporting Figure 6:** $\Delta$ RMSD during dissociation for the computationally studied ligands (residues 1-41). **ZMP** shows the highest change in $\Delta RMSD$ for residues in the P4 domain $(17-21)$ facing exit of the cavity. Here, we also note high changes for the missing residues that were modeled using ChimeraX.

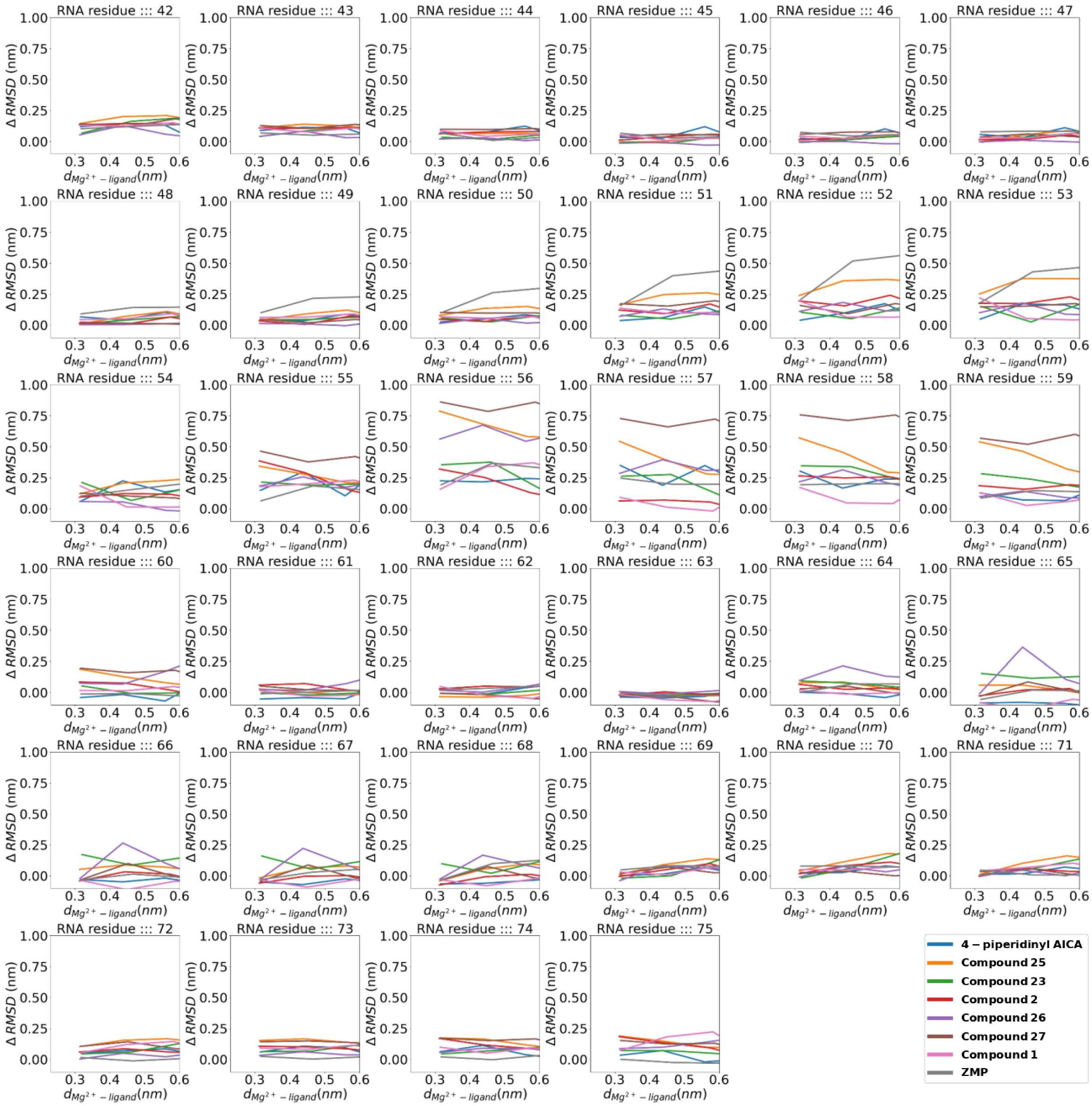

**Supporting Figure 7:** Δ RMSD during dissociation for the computationally studied ligands (residues 42-75).

column chromatography system using RediSep silica gel columns. Reactions were monitored by analytical thin-layer chromatography using EMD Silica Gel 60 F254 glass-backed pre-coated silica gel plates. The plates were visualized with UV light (254 nm) and stained with potassium by charring. Yields were reported as isolated, spectroscopically pure compounds.

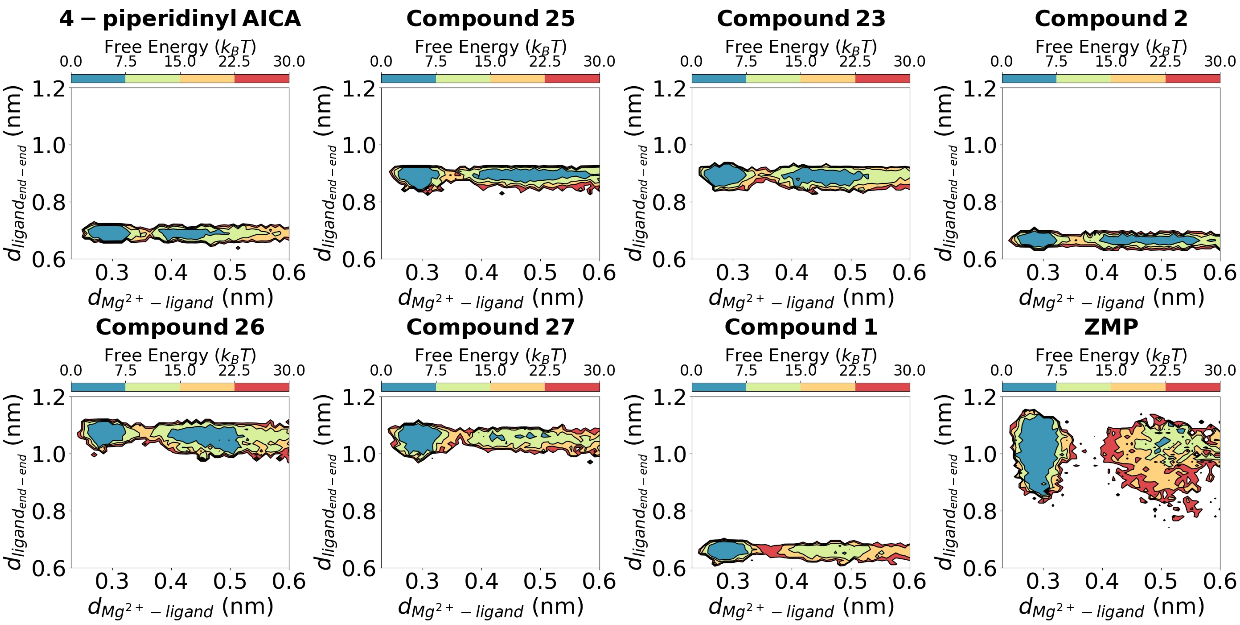

**Supporting Figure 8:** End-to-end $(l_3 - l_1)$ distance for the computationally studied ligands during dissociation showing the flexibility of **ZMP** particularly in the bound state $(d_M g^{2+} - ligand < 0.35nm)$ due to its higher rotatable bonds.

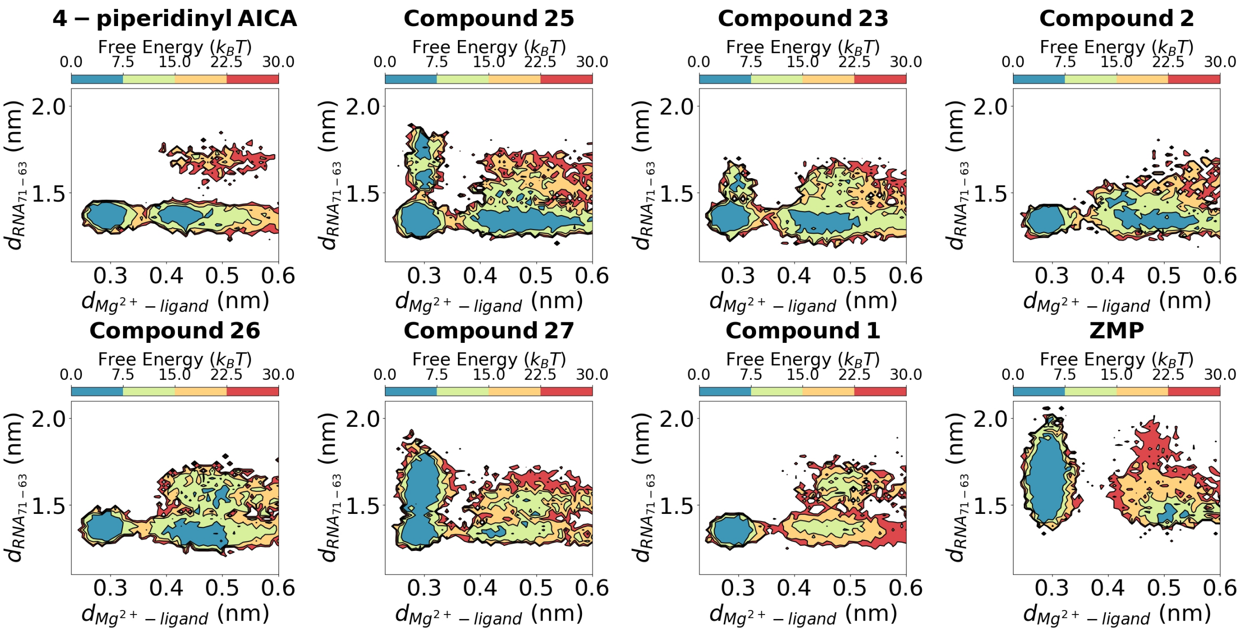

**Supporting Figure 9:** Distance between G71 backbone and G63 nucleobase for the computationally studied ligands during dissociation showing extension during bound state $(d_M g^{2+} - ligand < 0.35nm)$ for compounds **25,23,27,ZMP**.

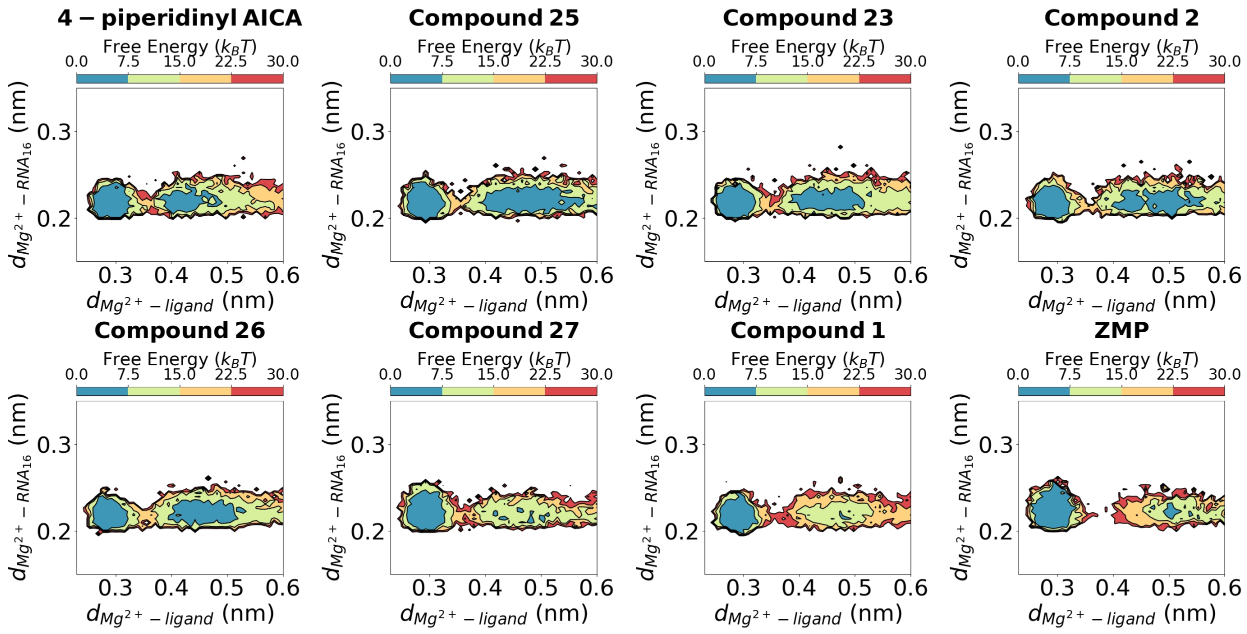

**Supporting Figure 10:** Distance between the coordinating oxygen atoms in the phosphate groups of U16, and $MG^{2+}$ for the computationally studied ligands during dissociation showing robustness of the simulations.

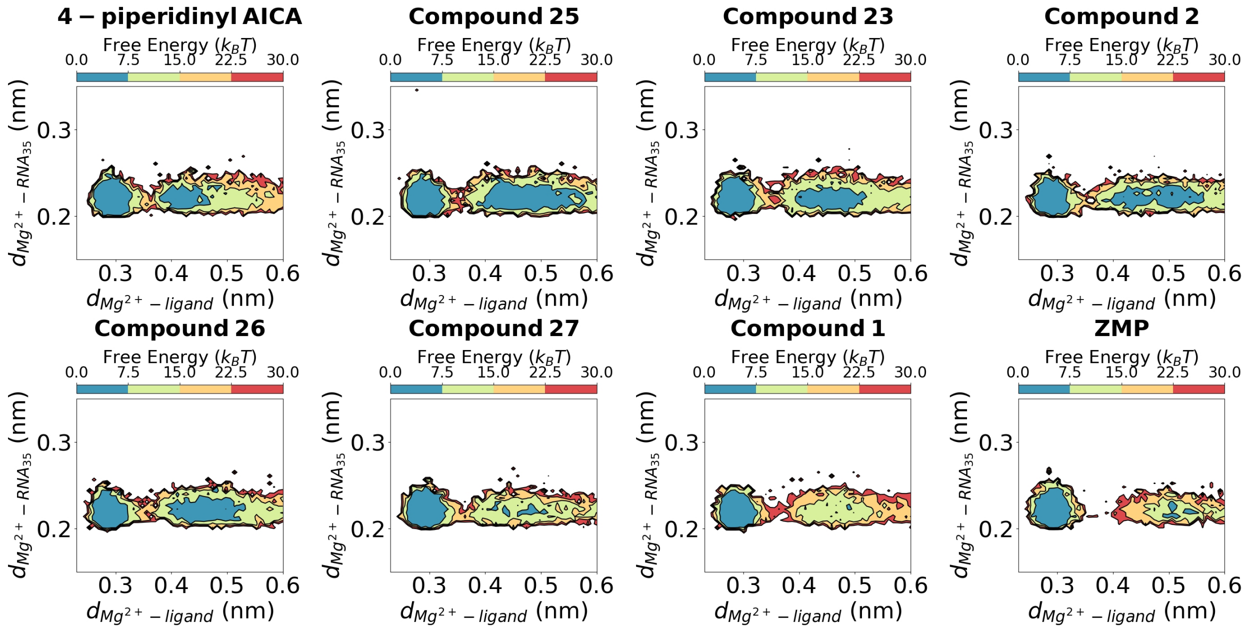

**Supporting Figure 11:** Distance between the coordinating oxygen atoms in the phosphate groups of C35, and $MG^{2+}$ for the computationally studied ligands during dissociation showing robustness of the simulations.

##### B. Materials:

Solvents and chemicals were purchased from commercial sources and used without further purification. Deuterated solvents were purchased from Cambridge Isotope Laboratories or Sigma Aldrich.

##### C. Instrumentation:

Reverse phase column chromatography (Teledyne CombiFlash) was performed using a RediSep C18 Gold 15.5 g or 30 g columns with UV/Vis detection. 1H NMR spectra were recorded on Bruker 500 MHz spectrometers and are reported relative to internal chloroform (1H, $\delta$ 7.26), methanol (1H,$\delta$ 3.31), and DMSO (1H,$\delta$ 2.50). Data for 1H NMR spectra are reported as follows: chemical shift ($\delta$ ppm), multiplicity (s = singlet, d = doublet, t = triplet, dd = doublet of doublet, ddd = doublet of doublet of doublet, m = multiplet, br=broad), coupling constants (Hz), and integration. 13C NMR was recorded on Bruker 500 MHz spectrometers at 125 MHz and is reported relative to internal chloroform (13C, $\delta$ 77.1), methanol (13C, $\delta$) 49.2), and DMSO (13C, $\delta$ 40.3). Low-resolution mass spectra were acquired on a Shimadzu LCMS 2030 system using electrospray ionization (ESI) in positive mode. Separation was performed on Phenomenex Kinetex C18 Column (50 X 2.1 mm; particle size: 2.6 $\mu$M). Analytes were eluted using a water/acetonitrile gradient with 0.1 (%) formic acid as a modifier with a 5-step LCMS program. LCMS program: Flow rate: 0.3 mL/min; Step 1: pre-equilibration for 2.50 min with 90% acetonitrile in water (0.1% formic acid), Step 2: 0.3% acetonitrile in water to 90% acetonitrile in water (0.1% formic acid) gradient over 1 min, Step 3: 90% acetonitrile in water (0.1% formic acid) for 0.25 min, Step 4: 90% acetonitrile in water to 0.6% acetonitrile in water (0.1% formic acid) over 0.5 min. High-resolution mass spectrometry data were acquired on an Agilent 6520 Accurate-Mass Q-TOF LC/MS System (Agilent Technologies, Inc.) equipped with a dual electrospray source operated in the positive ion mode. Separation was performed on Zorbax 300SB-C18 Poroshell column (2.1 mm x 150 mm; particle size 5 $\mu$m). The analytes were eluted using a water/acetonitrile gradient with 0.1% formic acid. Data were acquired at high resolution (1,700 m/z), 4 GHz. An internal mass calibration sample was infused continuously during the LC/MS runs to maintain mass accuracy during the run time. Data acquisition and analysis were performed using MassHunter Workstation Data Software, LCMS Data Acquisition (version B.06.01), and Qualitative Analysis (version B.07.00).

#### D. Synthetic Procedures:

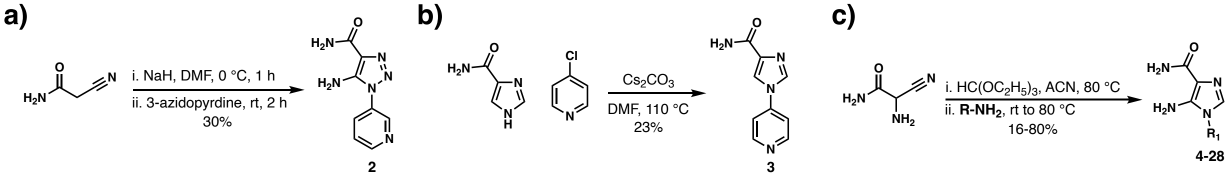

**Supporting Figure 12: Synthesis of *F. ulcerans* ZTP riboswitch interacting compounds 2-28.** **a)** Synthetic Scheme for compound **1**. **b)** Synthetic Scheme for compound **2**. **c)**Synthetic Scheme for compounds **4-28**.

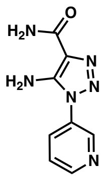

**5-amino-1-(pyridin-3-yl)-1H-1,2,3-triazole-4-carboxamide (2)** To a 0 ℃ suspension of sodium hydride (35.0 mg, 1.48 mmol, 1.77 equiv) in DMF (1.40 mL) was added a solution of 2-cyanoacetamide (77.0 mg, 0.923 mmol, 1.10 equiv) in DMF (1.40 mL) dropwise. The reaction mixture was maintained at 0 ℃ for 1h, then a solution of 3-azidopyridine (100 mg, 0.839 mmol, 1.00 equiv) in DMF (1.40 mL) was added slowly over 3 min. The reaction was allowed to warm to room temperature. After 2 h, the reaction was judged complete by LCMS and concentrated *in vacuo*. The crude residue was purified by flash column chromatography (silica gel; gradient elution, 0-10% DCM/MeOH) to afford 52.0 mg (30%) of **2** as tan solid: $^{1}$H NMR (500 MHz, DMSO-d6) $\delta$ 8.82 (s, 1H), 8.73 (s, 1H), 8.05 (d, J = 8.3 Hz, 2H), 7.65 (s, 1H), 7.24 (s, 2H), 6.56 (s, 3H); $^{13}$C NMR (126 MHz, DMS) $\delta$ 164.1, 149.9, 145.3, 145.1, 132.2, 131.7, 124.5, 121.6; LRMS calculated for C8H9N6O$^{+}$ [M+H]$^{+}$, m/z 205.20, measured LC/MS (ESI) Rt 2.03 min, m/z 205.15.

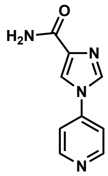

**1-(pyridin-4-yl)-1H-imidazole-4-carboxamide (3)** A solution of 1H-imidazole-4-carboxamide (250 mg, 2.25 mmol, 1 equiv), 4-chloropyridine (255 mg, 2.25 mmol, 1 equiv) and cesium carbonate (1.47 g, 4.50 mmol, 2 equiv) in DMF (4.50 mL) was heated to 110 ℃ for 18 h. After the consumption of the starting material, as judged by TLC, the reaction was allowed to cool to room temperature and concentrated *in vacuo*. The crude residue was purified by reverse phase flash column chromatography (RediSep Gold C18, gradient elution, 0-100% acetonitrile/water with 0.1% TFA) to afford 98 mg (23%) of **3** as a white solid: $^{1}$H NMR (500 MHz, DMSO-d6) $\delta$ 8.74 (dd, J = 6.4, 1.5 Hz, 2H), 8.65 (d, J = 1.5 Hz, 1H), 8.51 (d, J = 1.5 Hz, 1H), 7.95 (dd, J = 6.4, 1.6 Hz, 2H), 7.53 (s, 1H), 7.32 (s, 1H); $^{13}$C NMR (126 MHz, DMSO) $\delta$ 163.3, 150.5, 143.5 138.8, 135.8, 119.7, 114.4; LRMS calculated for C9H9N4O$^{+}$ [M+H]$^{+}$ m/z 189.20, measured LC/MS (ESI) Rt 1.66 min, m/z 189.06.

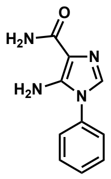

**5-amino-1-phenyl-1H-imidazole-4-carboxamide (4)** Triethylorthoformate (0.550 mL, 3.33 mmol, 1.1 equiv) was added to a solution of 2-amino-2-cyanoacetamide (300 mg, 3.03 mmol, 1 equiv) in acetonitrile (1 mL). The reaction mixture was heated to and maintained at 80 ℃ for 1 h. The reaction mixture was allowed to cool to room temperature, before the addition of aniline (0.300 mL, 3.33 mmol, 1.1 equiv). The reaction mixture was heated to and maintained at 80 ℃ for 18 h. Then, the reaction mixture was allowed to cool to room temperature, and the solvent was removed *in vacuo*. The crude residue was purified by flash column chromatography (silica gel, gradient elution 0-40% DCM/MeOH) to afford 212.3 mg (34%) of **4** as a tan solid: $^{1}$H NMR (500 MHz, DMSO-d6) $\delta$ 7.71 – 7.54 (m, 2H), 7.53 – 7.40 (m, 3H), 7.37 (s, 1H), 6.93 (s, 1H), 6.79 (s, 1H), 5.75 (s, 2H); $^{13}$C NMR (126 MHz, DMSO-d6) $\delta$ 166.7, 142.6, 134.66, 129.86, 129.7, 128.1, 124.5, 113.0; LRMS calculated for C10H11N4O$^{+}$ [M+H]$^{+}$ m/z 203.22, measured LC/MS (ESI) Rt 2.80 min, m/z 203.10.

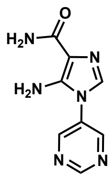

**5-amino-1-(pyrimidin-5-yl)-1H-imidazole-4-carboxamide (5)** To a solution of 2-amino-2-cyanoacetamide (100 mg, 1.01 mmol, 1 equiv) in acetonitrile (1 mL) was added triethyl orthoformate (168 $\mu$L, 1.01 mmol, 1 equiv). The reaction mixture was heated to 80 ℃ for 1 h. The reaction mixture was allowed to cool to room temperature, then pyrimidin-5-amine (192 mg, 2.02 mmol, 2 equiv) was added. The reaction mixture was heated to and maintained at 80 ℃ for 5 h. The reaction was allowed to cool to room temperature, then concentrated in vacuo. The crude residue was purified by flash column chromatography (silica gel, gradient elution, 0-30% DCM/MeOH) to afford 17.0 mg (8%) of **5** as a grey solid: $^{1}$H NMR (500 MHz, DMSO-d6) $\delta$ 9.28 (s, 1H), 9.02 (s, 2H), 7.49 (s, 1H), 6.98 (s, 1H), 6.83 (s, 1H), 5.97 (s, 2H); $^{13}$C NMR (126 MHz, DMSO-d6) $\delta$ 166.5, 157.4, 153.2, 142.8, 130.4, 129.5, 113.2; LRMS calculated for C8H9N6O$^{+}$ [M+H]$^{+}$ m/z 205.20, measured LC/MS (ESI) Rt 1.70 min, m/z 205.15.

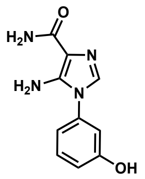

**5-amino-1-(3-hydroxyphenyl)-1H-imidazole-4-carboxamide (6)** To a solution of 2-amino-2-cyanoacetamide (100 mg, 1.01 mmol, 1 equiv) in acetonitrile (1 mL) was added triethyl orthoformate (168 $\mu$L, 1.01 mmol, 1 equiv). The reaction mixture was heated to 80 ℃ for 1 h. The reaction mixture was allowed to cool to room temperature, and then 3-aminophenol (220 mg, 2.02 mmol, 2 equiv) was added. The reaction mixture was heated to and maintained at 80 ℃ for 18 h. The reaction was allowed to cool to room temperature, then concentrated in vacuo. The crude residue was purified by flash column chromatography (silica gel, gradient elution, 0-30% DCM/MeOH) to afford 186 mg (84%) of **6** as a grey solid: $^1$H NMR (500 MHz, DMSO-d6) $\delta$ 7.36 – 7.31 (m, 1H), 7.31 (s, 1H), 6.90 – 6.88 (m, 2H), 6.86-6.85 (m, 2H), 6.76 (s, 1H), 5.72 (s, 2H); $^{13}$C NMR (126 MHz, DMSO-d6) $\delta$ 166.8, 158.4, 142.5, 135.5, 130.7, 129.6, 115.2, 114.9, 112.8, 111.3; LRMS calculated for C10H11N4O+ [M+H]+ m/z 219.22, measured LC/MS (ESI) Rt 2.27 min, m/z 219.15.

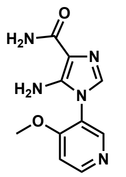

5-amino-1-(4-methoxypyridin-3-yl)-1H-imidazole-4-carboxamide (**7**) To a solution of 2-amino-2-cyanoacetamide (100 mg, 1.01 mmol, 1 equiv) in DMF (1 mL) was added triethyl orthoformate (168 $\mu$L, 1.01 mmol, 1 equiv). The reaction mixture was heated to 80 ℃ for 1 h. The reaction mixture was allowed to cool to room temperature, and then 4-methoxypyridin-3-amine (125.4 mg, 1.01 mmol, 1 equiv) was added. The reaction mixture was heated to and maintained at 80 ℃ for 2 h. The reaction was allowed to cool to room temperature, then concentrated in vacuo. The crude residue was purified by flash column chromatography (silica gel, gradient elution, 0-30% DCM/MeOH) to afford 38.3 mg (16%) of XX as a white solid: 1H NMR (500 MHz, DMSO-d6) $\delta$ 8.57 (d, J = 5.8 Hz, 1H), 8.40 (s, 1H), 7.31 (d, J = 5.8 Hz, 1H), 7.18 (s, 1H), 6.88 (s, 1H), 6.73 (s, 1H), 5.67 (s, 2H), 3.88 (s, 3H); 13C NMR (126 MHz, DMSO-d6) $\delta$ 166.7, 160.6, 152.0, 148.6, 143.9, 130.2, 120.1, 112.0, 108.5, 56.2; LRMS calculated for C10H12N5O2$^+$ [M+H]$^+$ m/z 234.24, measured LC/MS (ESI) Rt 1.65

min, m/z 234.10.

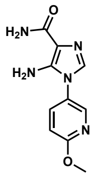

**5-amino-1-(6-methoxypyridin-3-yl)-1H-imidazole-4-carboxamide (8)** To a solution of 2-amino-2-cyanoacetamide (50 mg, 0.505 mmol, 1 equiv) in acetonitrile (500 $\mu$L) was added triethyl orthoformate (84.0 $\mu$L, 0.505 mmol, 1 equiv). The reaction mixture was heated to 80 ℃ for 1 h. The reaction mixture was allowed to cool to room temperature, and then 6-methoxypyridin-3-amine (62.7 mg, 0.505 mmol, 1 equiv) and pyridine (0.051 mmol, 0.1 equiv) were added. The reaction mixture was stirred at room temperature for 18 h. The crude residue was filtered and washed with acetonitrile (3x5 mL) and dried in vacuo to afford 59.8 mg (51%) of **8** as a tan solid: $^{1}$H NMR (500 MHz, DMSO-d6) $\delta$ 8.29 (d, J = 2.7 Hz, 1H), 7.84 (dd, J = 8.7, 2.8 Hz, 1H), 7.31 (s, 1H), 7.00 (dd, J = 8.9, 0.7 Hz, 1H), 6.93 (s, 1H), 6.81 – 6.74 (m, 1H), 5.76 (s, 2H), 3.91 (s, 3H); $^{13}$C NMR (126 MHz, DMSO-d6) $\delta$ 166.7, 163.0, 143.6, 143.2, 136.7, 129.9, 125.5, 112.7, 111.2, 53.7; LRMS calculated for C10H12N5O2$^{+}$ [M+H]$^{+}$ m/z 234.24, measured LC/MS (ESI) Rt 2.71 min, m/z 234.20.

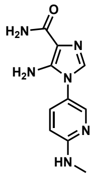

**5-amino-1-(6-(methylamino)pyridin-3-yl)-1H-imidazole-4-carboxamide (9)** To a solution of 2-amino-2-cyanoacetamide (46.9 mg, 0.475 mmol, 1 equiv) in acetonitrile (400 $\mu$L) was added triethyl orthoformate (78.7 $\mu$L, 0.475 mmol, 1 equiv). The reaction mixture was heated to 80 ℃ for 1 h. The reaction mixture was allowed to cool to room temperature, then N2-methylpyridine-2,5-diamine (58.3 mg, 0.475 mmol, 1 equiv) and pyridine (3.82 $\mu$L, 0.047 mmol, 0.1 equiv) were added. The reaction mixture was stirred at room temperature for 18 h. The crude residue was filtered and washed with acetonitrile (3 x 5 mL) and dried *in vacuo* to afford 50.8 mg (46%) of **9** as a tan solid: 1H NMR (500 MHz, DMSO-d6) $\delta$ $^{1}$H NMR (500 MHz, DMSO-d6) $\delta$ 8.03 (d, J = 2.6 Hz, 1H), 7.44 (dd, J = 8.9, 2.7 Hz, 1H), 7.19 (s, 1H), 6.88 (s, 1H), 5.62 (s, 2H), 2.81 (d, J = 4.8 Hz, 3H); $^{13}$C NMR (126 MHz, DMSO-d6) $\delta$ 166.7, 159.1, 144.5, 143.3, 134.3, 130.1, 119.9, 112.4, 107.9, 28.00; LRMS calculated for

C10H13N6O$^{+}$ [M+H]$^{+}$ m/z 233.25, measured LC/MS (ESI) Rt 1.62 min, m/z 233.10.

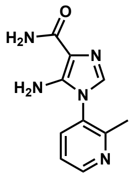

**5-amino-1-(2-methylpyridin-3-yl)-1H-imidazole-4-carboxamide (10)** To a solution of 2-amino-2-cyanoacetamide (200 mg, 2.02 mmol, 1 equiv) in acetonitrile (6 mL) was added triethyl orthoformate (336 $\mu$L, 2.02 mmol, 1 equiv). The reaction mixture was heated to 80 ℃ for 1 h. The reaction mixture was allowed to cool to room temperature, and then 2-methylpyridin-3-amine (218.4 mg, 2.02 mmol, 1 equiv) was added. The reaction mixture was heated to and maintained at 80 ℃ for 18 h. The reaction was allowed to cool to room temperature, then concentrated in vacuo. The crude residue was purified by flash column chromatography (silica gel, gradient elution, 0-30% DCM/MeOH) to afford 189.0 mg (43%) of 10 as an off-white solid: $^{1}$H NMR (500 MHz, DMSO-d6) $\delta$ 8.60 (dd, J = 4.8, 1.6 Hz, 1H), 7.76 (dd, J = 7.9, 1.6 Hz, 1H), 7.43 (dd, J = 7.9, 4.8 Hz, 1H), 7.24 (s, 1H), 6.91 (s, 1H), 6.76 (s, 1H), 5.72 (s, 2H), 2.29 (s, 3H); $^{13}$C NMR (126 MHz, DMSO-d6) $\delta$ 166.7, 155.8, 149.7, 143.5, 136.1, 129.6, 129.5, 122.3, 112.1, 20.6; LRMS calculated for C10H12N5O$^{+}$ [M+H]$^{+}$ m/z 218.24, measured LC/MS (ESI) Rt 1.75 min, m/z 218.15.

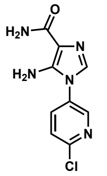

**5-amino-1-(6-chloropyridin-3-yl)-1H-imidazole-4-carboxamide (11**) To a solution of 2-amino-2-cyanoacetamide (100 mg, 1.01 mmol, 1 equiv) in acetonitrile (1 mL) was added triethyl orthoformate (168 $\mu$L, 1.01 mmol, 1 equiv). The reaction mixture was heated to 80 ℃ for 1 h. The reaction mixture was allowed to cool to room temperature, and then 6-chloropyridin-3-amine (129.8 mg, 1.01 mmol, 1 equiv) was added. The reaction mixture was heated and maintained at 80 ℃ for 18 h. The reaction was allowed to cool to room temperature, then concentrated *in vacuo*. The crude residue was purified by flash column chromatography (silica gel, gradient elution, 0-30% DCM/MeOH) to afford 98.8 mg (41%) of **11** as a white solid: $^{1}$H NMR (500 MHz, DMSO-d6) $\delta$ 8.60 (d, J = 2.9 Hz, 1H), 8.05 (dd, J = 8.4, 2.8 Hz, 1H), 7.73 (d, J = 8.5 Hz, 1H), 7.43 (s, 1H), 6.97 (s, 1H), 6.82 (s, 1H), 5.90

(s, 2H); $^{13}$C NMR (126 MHz, DMSO-d6) $\delta$ 166.5, 149.2, 146.0, 142.8, 136.3, 131.0, 129.6, 125.0, 113.1; LRMS calculated for C9H9ClN5O$^{+}$ [M+H]$^{+}$ m/z 238.65, measured LC/MS (ESI) Rt 2.55 min, m/z 238.15.

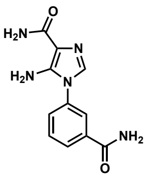

**5-amino-1-(3-carbamoylphenyl)-1H-imidazole-4-carboxamide (12)** To a solution of 2-amino-2-cyanoacetamide (100 mg, 1.01 mmol, 1 equiv) in DMF (1 mL) was added triethyl orthoformate (168 $\mu$L, 1.01 mmol, 1 equiv). The reaction mixture was heated to 80 ℃ for 1 h. The reaction mixture was allowed to cool to room temperature, and then 3-aminobenzamide (137 mg, 1.01 mmol, 1 equiv) was added. The reaction mixture was heated to and maintained at 80 ℃ for 2 h. The reaction was allowed to cool to room temperature, then concentrated in vacuo. The crude residue was purified by flash column chromatography (silica gel, gradient elution, 0-30% DCM/MeOH) to afford 42.6 mg (17%) of **12** as a tan solid: $^{1}$H NMR (500 MHz, DMSO-d6) $\delta$ 8.10 (s, 1H), 7.95 – 7.93 (m, 2H), 7.66 – 7.6 (m, 2H), 7.53 (s, 1H), 7.41 (s, 1H), 6.95 (s, 1H), 6.78 (s, 1H), 5.82 (s, 2H); $^{13}$C NMR (126 MHz, DMSO-d6) $\delta$ 166.9, 166.7, 142.6, 135.9, 134.6, 129.9, 129.7, 127.3, 127.2, 123.5, 112.9; LRMS calculated for C11H12N5O2+ [M+H]+ m/z 246.25, measured LC/MS (ESI) Rt 1.70 min, m/z 246.10. **(R)-5-amino-1-(piperidin-3-yl)-1H-imidazole-4-carboxamide (13)** To

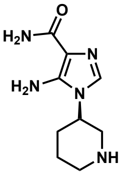

a solution of 2-amino-2-cyanoacetamide (50 mg, 0.505 mmol, 1 equiv) in acetonitrile (1.25 mL) was added triethyl orthoformate (168 $\mu$L, 1.01 mmol, 2 equiv). The reaction mixture was heated to 80 ℃ for 1 h. The reaction mixture was allowed to cool to room temperature, and then tert-butyl (R)-3-aminopiperidine-1-carboxylate (101.2 mg, 0.505 mmol, 1 equiv) and pyridine (4 $\mu$L, 0.05 mmol, 0.1 equiv) were added. The reaction mixture was stirred at 60 ℃ for 2 h. The crude residue was concentrated *in vacuo*. The crude product was then treated with trifluoroacetic acid (0.5 mL) for 1.5 h, followed by direct purification by flash column chromatography (silica gel, gradient elution, 0-30% DCM/MeOH) to afford 46.8 mg

(44% over two steps) of **13** as a white solid: $^1$H NMR (500 MHz, DMSO-d6) $\delta$ 7.30 (s, 1H), 6.82 (s, 1H), 6.69 (s, 1H), 5.79 (s, 2H), 5.75 (s, 1H, NH), 4.21 – 4.17 (m, 1H), 3.44 (dd, J = 12.0, 4.1 Hz, 1H), 3.30 (m, 1H), 3.21 (t, J = 11.9 Hz, 1H), 2.85 (td, J = 12.8, 2.9 Hz, 1H), 2.11 – 2.02 (m, 1H), 1.95 (td, J = 13.1, 3.7 Hz, 2H), 1.82 – 1.69 (m, 1H); $^{13}$C NMR (126 MHz, DMSO-d6) $\delta$ 166.6, 142.2, 127.2, 112.9, 47.3, 45.7, 42.3, 28.3, 21.4; LRMS calculated for C9H16N5O$^+$ [M+H]$^+$ m/z 210.26, measured LC/MS (ESI) Rt 1.62 min, m/z 210.15.

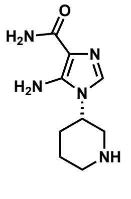

**(S)-5-amino-1-(piperidin-3-yl)-1H-imidazole-4-carboxamide (14)** To a solution of 2-amino-2-cyanoacetamide (50 mg, 0.505 mmol, 1 equiv) in acetonitrile (1.25 mL) was added triethyl orthoformate (168 $\mu$L, 1.01 mmol, 2 equiv). The reaction mixture was heated to 80 ℃ for 1 h. The reaction mixture was allowed to cool to room temperature, and then tert-butyl (S)-3-aminopiperidine-1-carboxylate (101.2 mg, 0.505 mmol, 1 equiv) and pyridine (4 $\mu$L, 0.05 mmol, 0.1 equiv) were added. The reaction mixture was stirred at 60 ℃ for 2 h. The crude residue was concentrated *in vacuo*. The crude product was then treated with trifluoroacetic acid (0.5 mL) for 1.5 h, followed by direct purification by flash column chromatography (silica gel, gradient elution, 0-30% DCM/MeOH) to afford 43.8 mg (41% over two steps) of **14** as a white solid: $^1$H NMR (500 MHz, DMSO-d6) $\delta$ 7.30 (s, 1H), 6.82 (s, 1H), 6.69 (s, 1H), 5.79 (s, 2H), 5.75 (s, 1H, NH), 4.21 – 4.16 (m, 1H), 3.44 (dd, J = 11.9, 4.1 Hz, 1H), 3.31 (d, J = 18.7 Hz, 2H), 3.21 (t, J = 11.9 Hz, 1H), 2.85 (td, J = 12.7, 2.8 Hz, 1H), 2.07 – 2.04 (m, 1H), 1.98-19.1 (m, 2H), 1.79 – 1.73 (m, 1H); $^{13}$C NMR (126 MHz, DMSO-d6) $\delta$ 166.8, 158.3, 142.5, 135.6, 130.7, 129.6, 115.2, 114.9, 112.8, 111.4; LRMS calculated for C9H16N5O$^+$ [M+H]$^+$ m/z 210.26, measured LC/MS (ESI) Rt 1.62 min, m/z 210.20.

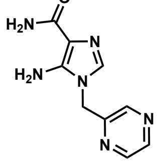

**5-amino-1-(pyrazin-2-ylmethyl)-1H-imidazole-4-carboxamide (15)** To a solution of 2-amino-2-cyanoacetamide (50 mg, 0.505 mmol, 1 equiv) in acetonitrile (1.25 mL) was added triethyl orthoformate (168 $\mu$L, 1.01 mmol, 2 equiv) and pyridine (4 $\mu$L, 0.05 mmol, 0.1 equiv). The reaction mixture was heated to 80 ℃ for 1 h. The reaction mixture was allowed to cool to room temperature, and then pyrazin-2-ylmethanamine (54.6 mg, 0.505 mmol, 1 equiv) was added. The reaction mixture was stirred at 60 ℃ for 1 h. Then, the reaction was allowed to cool to room temperature, and hexane (3 mL) was added. The residue was allowed to stir for 1h, filtered and washed with acetonitrile and hexanes (1:1, 3 x 3 mL) to afford 57.9 mg (53%) of **15** as a white solid: $^1$H NMR (500 MHz, DMSO-d6) $\delta$ 8.62 (m, 1H), 8.59 (d, J = 2.5 Hz, 1H), 8.53 (d, J = 1.5 Hz, 1H), 7.22 (s, 1H), 6.79 (s, 1H), 6.65 (s, 1H), 5.81 (s, 2H), 5.27 (s, 2H); $^{13}$C NMR (126 MHz, DMSO-d6) $\delta$ 166.7, 151.6, 144.4, 143.9, 143.4, 143.1, 130.3, 112.6, 45.3; LRMS calculated for C9H11N6O$^+$ [M+H]$^+$ m/z 219.23, measured LC/MS (ESI) Rt 1.62 min, m/z 219.15.

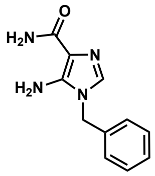

**5-amino-1-benzyl-1H-imidazole-4-carboxamide (16)** To a solution of 2-amino-2-cyanoacetamide (50 mg, 0.505 mmol, 1 equiv) in acetonitrile (1.25 mL) was added triethyl orthoformate (168 $\mu$L, 1.01 mmol, 2 equiv) and pyridine (4 $\mu$L, 0.05 mmol, 0.1 equiv). The reaction mixture was heated to 80 ℃ for 1 h. The reaction mixture was allowed to cool to room temperature, and then benzylamine (54.1 mg, 0.505 mmol, 1 equiv) was added. The reaction mixture was stirred at 60 ℃ for 1 h. Then, the reaction was allowed to cool to room temperature, and hexane (3 mL) was added. The residue was allowed to stir for 1h, filtered and washed with acetonitrile and hexanes (1:1, 3 x 3 mL) to afford 57.9 mg (53%) of **16** as a white solid: $^1$H NMR (500 MHz, DMSO-d6) $\delta$ 7.35 (t, J = 7.4 Hz, 2H), 7.28 (t, J = 7.3 Hz, 1H), 7.20 (d, J = 6.9 Hz, 2H), 7.18 (s, 1H), 6.77 (s, 1H), 6.63 (s, 1H), 5.83 (s, 2H), 5.07 (s, 2H); $^{13}$C NMR (126 MHz, DMSO-d6) $\delta$ 166.7, 143.0, 136.9, 129.9, 128.6, 127.5, 127.2, 11257, 45.7; LRMS calculated for C11H13N4O$^+$ [M+H]+ m/z 217.25, measured LC/MS (ESI) Rt 2.53 min, m/z 217.20.

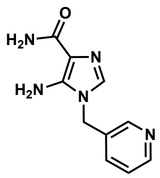

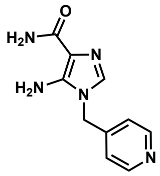

**5-amino-1-(pyridin-4-ylmethyl)-1H-imidazole-4-carboxamide (18)** To a solution of 2-amino-2-cyanoacetamide (50 mg, 0.505 mmol, 1 equiv) in acetonitrile (1.25 mL) was added triethyl orthoformate (168 $\mu$L, 1.01 mmol, 2 equiv) and pyridine (4 $\mu$L, 0.05 mmol, 0.1 equiv). The reaction mixture was heated to 80 ℃ for 1 h. The reaction mixture was allowed to cool to room temperature, and then 4-aminomethylpyridine (54.1 mg, 0.505 mmol, 1 equiv) was added. The reaction mixture was stirred at 60 ℃ for 1 h. Then, the reaction was allowed to cool to room temperature, and hexane (3 mL) was added. The residue was allowed to stir for 1h, filtered and washed with acetonitrile and hexanes (1:1, 3 x 3 mL) to afford 79.2 mg (73%) of **18** as a white solid: $^1$H NMR (500 MHz, DMSO-d6) $\delta$ 8.53 (d, J = 6.1 Hz, 2H), 7.23 (s, 1H), 7.09 (d, J = 6.1 Hz, 2H), 6.81 (s, 1H), 6.66 (s, 1H), 5.84 (s, 2H), 5.15 (s, 2H); $^{13}$C NMR (126 MHz, DMSO-d6) $\delta$ 166.7, 149.9, 145.9, 143.1, 130.1, 121.8, 112.5, 44.7; LRMS calculated for C10H12N5O$^+$ [M+H]+ m/z 218.24, measured LC/MS (ESI) Rt 1.55 min, m/z 218.10.

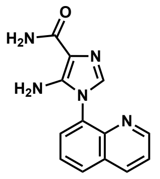

**5-amino-1-(quinolin-8-yl)-1H-imidazole-4-carboxamide (19)** To a solution of 2-amino-2-cyanoacetamide (100 mg, 1.01 mmol, 1.0 equiv) in acetonitrile (3 mL) was added triethyl orthoformate (168 $\mu$L, 1.01 mmol, 1.0 equiv). The reaction mixture was heated to 80 ℃ for 1 h. The reaction mixture was allowed to cool to room temperature, and then quinoline-8-amine (146 mg, 1.01 mmol, 1 equiv) was added. The reaction mixture was heated to and maintained at 85 ℃ for 18 h. The reaction was allowed to cool to room temperature, then concentrated *in vacuo*. The crude residue was purified by flash column chromatography (silica gel, gradient elution, 0-40% DCM/MeOH) to afford 149 mg (58%) of **19** as a yellow solid: $^{1}$H NMR (500 MHz, DMSO-d6) $\delta$ 8.96 (dd, J = 4.2, 1.7 Hz, 1H), 8.55 (dd, J = 8.4, 1.7 Hz, 1H), 8.17 (dd, J = 8.4, 1.4 Hz, 1H), 7.87 (dd, J = 7.4, 1.4 Hz, 1H), 7.77 (t, J = 7.8 Hz, 1H), 7.68 (dd, J = 8.3, 4.2 Hz, 1H), 7.32 (s, 1H), 6.92 (s, 1H), 6.76 (s, 1H), 5.52 (s, 2H); $^{13}$C NMR (126 MHz, DMSO-d6) $\delta$ 166.8, 151.4, 144.2, 142.8, 136.7, 131.6, 131.3, 129.4, 128.9, 128.1, 126.5, 122.4, 112.4; LRMS calculated for $C13H12N5O^{+}$ [M+H]+ m/z 254.27, measured LC/MS (ESI) Rt 2.82 min, m/z 254.10.

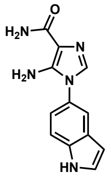

**5-amino-1-(1H-indol-5-yl)-1H-imidazole-4-carboxamide (20)** To a solution of 2-amino-2-cyanoacetamide (200 mg, 2.02 mmol, 1.0 equiv) in acetonitrile (6 mL) was added triethyl orthoformate (336 $\mu$L, 2.02 mmol, 1.0 equiv). The reaction mixture was heated to 80 ℃ for 1 h. The reaction mixture was allowed to cool to room temperature, and then 5-aminoindole (531 mg, 4.04 mmol, 2 equiv) was added. The reaction mixture was heated to and maintained at 85 ℃ for 18 h. The reaction was allowed to cool to room temperature, then concentrated *in vacuo*. The crude residue was purified by flash column chromatography (silica gel, gradient elution, 0-30% DCM/MeOH) to afford 134 mg (26%) of **20** as a white solid: $^{1}$H NMR (500 MHz, DMSO-d6) $\delta$ 11.37 (s, 1H), 7.62 (d, J = 2.0 Hz, 1H), 7.54 (d, J = 8.5 Hz, 1H), 7.49 (t, J = 2.8 Hz, 1H), 7.28 (s, 1H), 7.12 dd, J = 8.5, 2.1 Hz, 1 H), 6.88 (s, 1H), 6.73 (s, 1H), 6.54 (s, 1H), 5.61 (s, 2H); $^{13}$C NMR (126 MHz, DMSO-d6) $\delta$ 166.8, 143.2, 135.3, 130.2, 127.9, 127.3, 126.1, 118.3, 116.7, 112.5, 112.3, 101.7; LRMS calculated for $C12H12N5O^{+}$ [M+H]+ m/z 242.26, measured LC/MS (ESI) Rt 3.09 min, m/z 242.15.

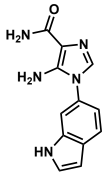

**5-amino-1-(1H-indol-6-yl)-1H-imidazole-4-carboxamide (21)** To a solution of 2-amino-2-cyanoacetamide (100 mg, 1.01 mmol, 1.0 equiv) in acetonitrile (3 mL) was added triethyl orthoformate (168 $\mu$L, 1.01 mmol, 1.0 equiv). The reaction mixture was heated to 80 ℃ for 1 h. The reaction mixture was allowed to cool to room temperature, and then

6-aminoindole (265 mg, 2.02 mmol, 2 equiv) was added. The reaction mixture was heated to and maintained at 85 ℃ for 18 h. The reaction was allowed to cool to room temperature, then concentrated *in vacuo*. The crude residue was purified by flash column chromatography (silica gel, gradient elution, 0-30% DCM/MeOH) to afford 52 mg (21%) of **21** as a white solid: $^{1}$H NMR (500 MHz, DMSO-d6) $\delta$ 11.37 (s, 1H), 7.69 (d, J = 8.3 Hz, 1H), 7.50 – 7.45 (m, 2H), 7.32 (s, 1H), 7.05 (dd, J = 8.3, 1.9 Hz, 1H), 6.88 (s, 1H), 6.73 (s, 1H), 6.53 (s, 1H), 5.67 (s, 2H); $^{13}$C NMR (126 MHz, DMSO-d6) $\delta$ 166.8, 143.1, 135.6, 130.1, 127.9, 127.4, 127.2, 120.9, 116.1, 112.5, 108.2, 101.3; LRMS calculated for $C12H12N5O^{+}$ [M+H]+ m/z 242.26, measured LC/MS (ESI) Rt 3.45 min, m/z 242.05.

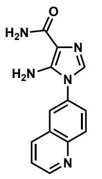

**5-amino-1-(quinolin-6-yl)-1H-imidazole-4-carboxamide (22)** To a solution of 2-amino-2-cyanoacetamide (200 mg, 2.02 mmol, 1 equiv) in acetonitrile (6 mL) was added triethyl orthoformate (336 $\mu$L, 2.02 mmol, 1 equiv). The reaction mixture was heated to 80 ℃ for 1 h. The reaction mixture was allowed to cool to room temperature, and then 6-aminoquinoline (582.4 mg, 4.04 mmol, 2 equiv) was added. The reaction mixture was heated to and maintained at 80 ℃ for 3.5 h. The reaction was allowed to cool to room temperature, then concentrated *in vacuo*. The crude residue was purified by flash column chromatography (silica gel, gradient elution, 0-30% DCM/MeOH) to afford 131 mg (26%) of **22** as an off-white solid: $^{1}$H NMR (500 MHz, DMSO-d6) $\delta$ 8.99 (dd, J = 4.2, 1.7 Hz, 1H), 8.47 (d, J = 8.0 Hz, 1H), 8.19 (d, J = 9.0 Hz, 1H), 8.16 (d, J = 2.5 Hz, 1H), 7.89 (dd, J = 9.0, 2.4 Hz, 1H), 7.64 (dd, J = 8.3, 4.2 Hz, 1H), 7.51 (s, 1H), 6.98 (s, 1H), 6.82 (s, 1H), 5.91 (s, 2H); $^{13}$C NMR (126 MHz, DMSO-d6) $\delta$ 166.7, 151.4, 146.7, 142.8, 136.3, 132.4, 130.7, 129.7, 128.1, 126.5, 123.2, 122.3, 113.0; LRMS calculated for C13H12N5O$^{+}$ [M+H]+ m/z 254.27, measured LC/MS (ESI) Rt 2.11 min, m/z 254.15.

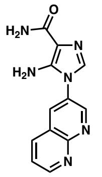

**5-amino-1-(1,8-naphthyridin-3-yl)-1H-imidazole-4-carboxamide (23)** To a solution of 2-amino-2-cyanoacetamide (54.0 mg, 0.543 mmol, 1.05 equiv) in acetonitrile (2 mL) was added triethyl orthoformate (90.0 $\mu$L, 0.543 mmol, 1 equiv). The reaction mixture was heated to 80 ℃ for 1 h. The reaction mixture was allowed to cool to room temperature, and then 1,8-naphthyridin-3-amine (75 mg, 0.517 mmol, 1 equiv) was added. The reaction mixture was heated to and maintained at 65 ℃ for 18 h. The reaction was allowed to cool to room temperature, then concentrated *in vacuo*. The crude residue was purified by flash column chromatography (silica gel, gradient elution, 0-30% DCM/MeOH) to afford 40 mg (30%) of **23** as an off-white solid: 1H NMR (500 MHz, DMSO-d6) $\delta$ 9.20 (s, 1H), 9.17 (s, 1H), 8.68 (s, 1H), 8.58 (d, J = 8.2 Hz, 1H), 7.76 (m, 1H), 7.58 (s, 1H), 7.00 (s, 1H), 6.84

(s, 1H), 6.00 (s, 2H); 13C NMR (126 MHz, DMSO-d6) δ 166.6, 154.3, 154.2, 150.3, 143.1, 138.1, 132.6, 129.7, 128.9, 123.2, 122.3, 113.0; LRMS calculated for $C12H11N6O2^+$ [M+H]+ m/z 255.26, measured LC/MS (ESI) Rt 1.94 min, m/z 255.05.

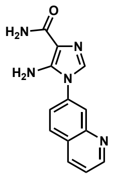

**5-amino-1-(quinolin-7-yl)-1H-imidazole-4-carboxamide (24)** To a solution of 2-amino-2-cyanoacetamide (100 mg, 1.01 mmol, 1.0 equiv) in acetonitrile (3 mL) was added triethyl orthoformate (168 μL, 1.01 mmol, 1 equiv). The reaction mixture was heated to 80 ℃ for 1 h. The reaction mixture was allowed to cool to room temperature, and then 1,8-naphthyridin-3-amine (75 mg, 0.517 mmol, 1 equiv) was added. The reaction mixture was heated to and maintained at 85 ℃ for 18 h. The reaction was allowed to cool to room temperature, then concentrated *in vacuo*. The crude residue was purified by flash column chromatography (silica gel, gradient elution, 0-30% DCM/MeOH) to afford 31 mg (12%) of **24** as an off-white solid: $^1$H NMR (500 MHz, DMSO-d6) δ 9.00 (dd, J = 4.2, 1.7 Hz, 1H), 8.52 – 8.46 (m, 1H), 8.19 (d, J = 8.7 Hz, 1H), 8.15 (d, J = 2.1 Hz, 1H), 7.77 (dd, J = 8.7, 2.2 Hz, 1H), 7.63 (dd, J = 8.3, 4.2 Hz, 1H), 7.56 (s, 1H), 6.98 (s, 1H), 6.83 (s, 1H), 5.92 (s, 2H); $^{13}$C NMR (126 MHz, DMSO-d6) δ 166.7, 151.7, 147.8, 142.7, 136.0, 135.4, 129.9, 129.8, 127.1, 123.4, 123.4, 122.1, 113.2; LRMS calculated for $C13H12N5O^+$ [M+H]+ m/z 254.27, measured LC/MS (ESI) Rt 2.29 min, m/z 254.10.

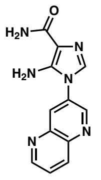

**5-amino-1-(1,5-naphthyridin-3-yl)-1H-imidazole-4-carboxamide (25)** To a solution of 2-amino-2-cyanoacetamide (56 mg, 0.574 mmol, 1.0 equiv) in acetonitrile (2 mL) was added triethyl orthoformate (114 μL, 0.688 mmol, 1.2 equiv). The reaction mixture was heated to 80 ℃ for 1 h. The reaction mixture was allowed to cool to room temperature, and then 1,5-naphthyridin-3-amine (100 mg, 0.688 mmol, 1.2 equiv) was added. The reaction mixture was heated to and maintained at 85 ℃ for 5 h. The reaction was allowed to cool to room temperature, then concentrated *in vacuo*. The crude residue was purified by flash column chromatography (silica gel, gradient elution, 0-30% DCM/MeOH) to afford 23 mg (16%) of **25** as a tan solid: $^{1}$H NMR (500 MHz, DMSO-d6) $\delta$ 9.17 (s, 1H), 9.11 (s, 1H), 8.61 (s, 1H), 8.55 (d, J = 8.6 Hz, 1H), 7.90 – 7.87 (m, 1H), 7.63 (s, 1H), 7.01 (s, 1H), 6.85 (s, 1H), 6.02 (s, 2H); $^{13}$C NMR (126 MHz, DMSO-d6) $\delta$ 166.6, 152.5, 147.9, 142.9, 142.8, 142.2, 136.8, 131.4, 131.3, 129.9, 125.3, 113.3; LRMS calculated for C12H11N6O+ [M+H]+ m/z 255.26, measured LC/MS (ESI) Rt 1.86 min, m/z 255.10.

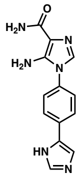

**1-(4-(1H-imidazol-5-yl)phenyl)-5-amino-1H-imidazole-4-carboxamide (26)** To a solution of 2-amino-2-cyanoacetamide (93 mg, 0.942 mmol, 1.0 equiv) in acetonitrile (1.6 mL) was added triethyl orthoformate (156 $\mu$L, 0.942 mmol, 1.0 equiv). The reaction mixture was heated to 80 ℃ for 1 h. The reaction mixture was allowed to cool to room temperature, and then 4-(1H-imidazol-5-yl)aniline (150 mg, 0.942 mmol, 1 equiv) was added. The reaction mixture was heated to and maintained at 85 ℃ for 18 h. The reaction was allowed to cool to room temperature, then concentrated *in vacuo*. The crude residue was purified by flash column chromatography (silica gel, gradient elution, 0-30% DCM/MeOH) to afford 63 mg (25%) of **26** as a grey solid: $^{1}$H NMR (500 MHz, DMSO-d6) $\delta$ 7.93 (d, J = 8.6 Hz, 2H), 7.80 (s, 1H), 7.71 (s, 1H), 7.48 (d, J = 8.5 Hz, 2H), 7.37 (s, 1H), 6.93 (s, 1H), 6.77 (s, 1H), 5.77 (s, 2H); $^{13}$C NMR (126 MHz, DMSO-d6) $\delta$ 166.8, 142.7, 136.3, 135.7, 134.2, 132.3, 129.6, 125.3, 125.0, 124.7, 112.9; LRMS calculated for C13H13N6O$^{+}$ [M+H]+ m/z 269.29, measured LC/MS (ESI) Rt 1.65 min, m/z 269.15.

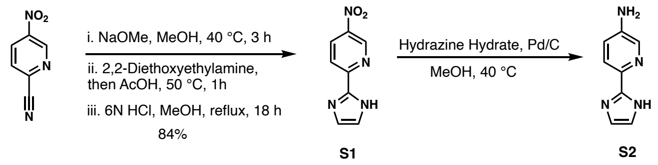

**Supporting Figure 13: Scheme S2. Synthesis of aniline S2**

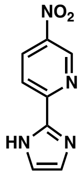

**2-(1H-imidazol-2-yl)-5-nitropyridine (S1)** To a solution of 5-nitropicolinonitrile (500 mg, 3.35 mmol, 1.0 equiv) in methanol (136 $\mu$L) was added sodium methoxide (36.2 mg, 0.671 mmol. 0.2 equiv). The reaction was heated and maintained at 40 ℃ for 3 h. Then, the reaction was allowed to cool to room temperature, and 2,2-diethoxyethan-1-amine (488 $\mu$L, 3.35 mmol, 1.0 equiv) was added. Let stir for 10 min, then acetic acid (384 $\mu$L, 6.71 mmol, 2.0 equiv) was added (drop wise). Once the addition was complete, the reaction was heated and maintained at 50 ℃ for 1 h. The reaction was removed from heat and allowed to cool to room temperature. Then methanol (6 mL) and 6N HCl (2 mL) were added, and the reaction was heated and maintained at reflux for 18 h. The reaction was allowed to cool to room temperature and concentrated *in vacuo*. The crude residue was diluted with water/Et2O (1:1, 10 mL). The organic and aqueous layers were separated, and the aqueous layer was adjusted to pH 8-9 with 1M NaOH. The aqueous layer was cooled and maintained at 0 ℃ for 1 h. The precipitate was collected by vacuum filtration to afford 468 mg (73%) of **S1** as a yellow solid: $^1$H NMR (500 MHz, DMSO-d6) $\delta$ 13.26 (s, 1H), 9.35 (d, J = 2.6 Hz, 1H), 8.64 (dd, J = 8.8, 2.6 Hz, 1H), 8.24 (d, J = 8.8 Hz, 1H), 7.40 (s, 1H), 7.23 (s, 1H); $^{13}$C NMR (126 MHz, DMSO-d6) $\delta$ 153.0, 144.9, 143.9, 142.9, 132.8, 131.2, 121.0, 119.6. Consistent with literature values[21].

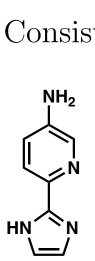

**6-(2,3-dihydro-1H-imidazol-2-yl)pyridin-3-amine (S2)** To a solution of 2-(1H-imidazol-2-yl)-5-nitropyridine (S1B) (226 mg, 1 Eq, 1.19 mmol) in MeOH (6 mL) was added Pd/C (12.6 mg, 0.1 Eq, 0.119 mmol, 10 wt%). Then, the solution was parged with $H_2$(g) for

5 min. The reaction was maintained under a Hydrogen atmosphere at room temperature for 2 h, filtered through a plug of celite, and concentrated *in vacuo* to afford 175 mg (92%) of **S2** as a yellow solid: $^1$H NMR (500 MHz, DMSO-d6) $\delta$ 12.24 (s, 1H), 7.94 (d, J = 2.7 Hz, 1H), 7.70 (d, J = 8.5 Hz, 1H), 7.04 (s, 1H), 6.99 (dd, J = 8.5, 2.7 Hz, 1H), 6.91 (s, 1H), 5.51 (s, 2H); $^{13}$C NMR (126 MHz, DMSO-d6) $\delta$ 146.6, 144.5, 137.7, 134.8, 128.4, 120.5, 120.1, 116.9. Consistent with literature values[21].

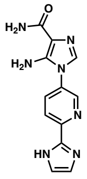

**1-(6-(1H-imidazol-2-yl)pyridin-3-yl)-5-amino-1H-imidazole-4-carboxamide (27)** To a solution of 2-amino-2-cyanoacetamide (61 mg, 0.642 mmol, 1.0 equiv) in acetonitrile (1.0 mL) was added triethyl orthoformate (103 $\mu$L, 0.642 mmol, 1.0 equiv). The reaction mixture was heated to 80 ℃ for 1 h. The reaction mixture was allowed to cool to room temperature, and then **S2** (100 mg, 0.942 mmol, 1 equiv) was added. The reaction mixture was heated to and maintained at 85 ℃ for 18 h. The reaction was allowed to cool to room temperature and concentrated *in vacuo*. The crude residue was purified by flash column chromatography (silica gel, gradient elution, 0-30% DCM/MeOH) to afford 133 mg (79%) of **27** as a tan solid (TFA salt): $^1$H NMR (500 MHz, DMSO-d6) $\delta$ 8.99 (s, 1H), 8.37-8.33 (m, 2H), 7.84 (s, 2H), 7.67 (s, 1H), 6.99 (s, 2H, exchanges with $D_2O$); $^{13}$C NMR (126 MHz, DMSO-d6) $\delta$ 166.2, 158.2 (q, $J$ = 37 Hz, TFA), 146.0, 142.8, 142.2, 141.7, 134.1, 132.8, 129.8, 122.1, 121.8, 112.8; LRMS calculated for C12H12N7O$^+$ [M+H]+ m/z 270.28, measured LC/MS (ESI) Rt 1.59 min, m/z 270.15.

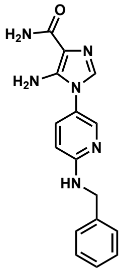

**5-amino-1-(6-(benzylamino)pyridin-3-yl)-1H-imidazole-4-carboxamide (28)** To a solution of 2-amino-2-cyanoacetamide (50.0 mg, 0.505 mmol, 1 equiv) in acetonitrile (500 $\mu$L) was added triethyl orthoformate (84.0 $\mu$L, 0.505 mmol, 1 equiv). The reaction mixture was heated to 80 ℃ for 1 h. The reaction mixture was allowed to cool to room temperature, and then N2-methylpyridine-2,5-diamine (58.3 mg, 0.475 mmol, 1 equiv) and pyridine (3.82 $\mu$L, 0.047 mmol, 0.1 equiv) were added. The reaction mixture was stirred at room temperature for 18 h. The crude residue was filtered and washed with acetonitrile (3x5 mL) and dried *in vacuo* to afford 50.8 mg (46%) of **28** as a tan solid: $^1$H NMR (500 MHz, DMSO-d6) $\delta$ 8.02 (d, J = 2.6 Hz, 1H), 7.49 (t, J = 6.0 Hz, 1H), 7.46 (dd, J = 8.8, 2.7 Hz, 1H), 7.38 – 7.29 (m, 4H), 7.26 – 7.21 (m, 1H), 7.20 (s, 1H), 6.86 (s, 1H), 6.72 (s, 1H), 6.64 (d, J = 8.9 Hz, 1H), 5.63 (s, 2H), 4.52 (d, J = 6.0 Hz, 2H); $^{13}$C NMR (126 MHz, DMSO-d6) $\delta$ 166.7, 158.3, 144.4, 143.3, 140.2, 134.4, 130.1, 128.3, 128.1, 127.2, 127.2, 126.6, 120.4, 112.5, 108.3, 44.2; LRMS calculated for $C16H17N6O^+$ [M+H]+ m/z 309.35, measured LC/MS (ESI) Rt 3.37 min, m/z 309.20.

#### E. NMR Spectra

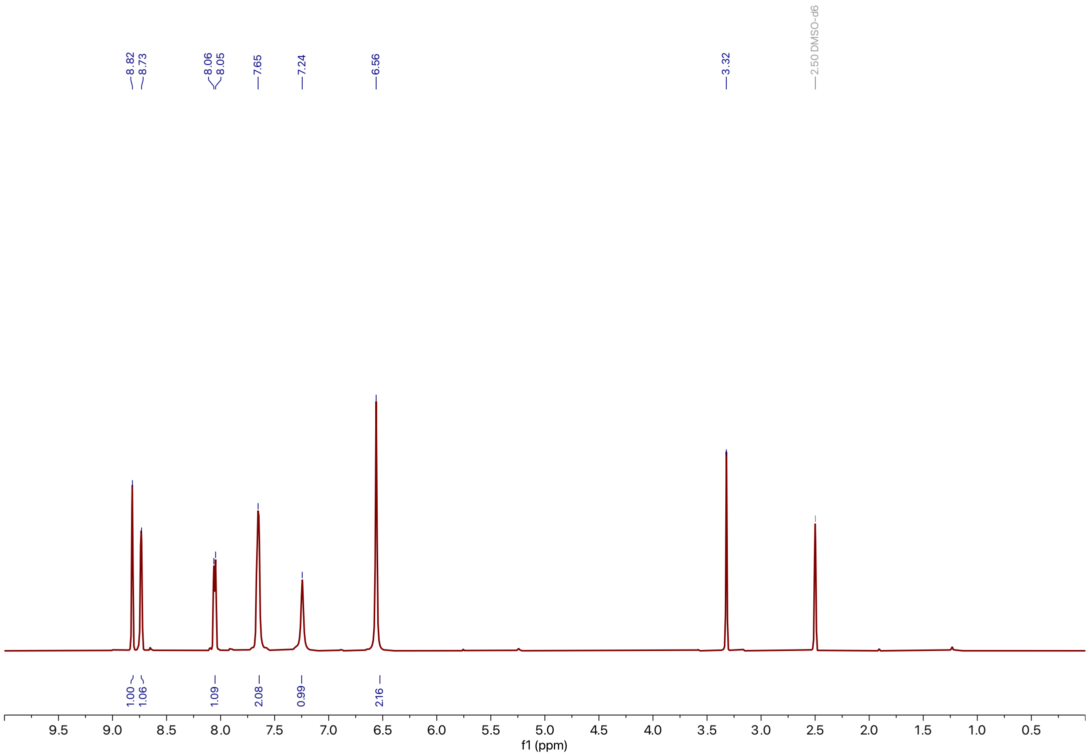

**Supporting Figure 14:** $^1$H NMR spectra of **2**

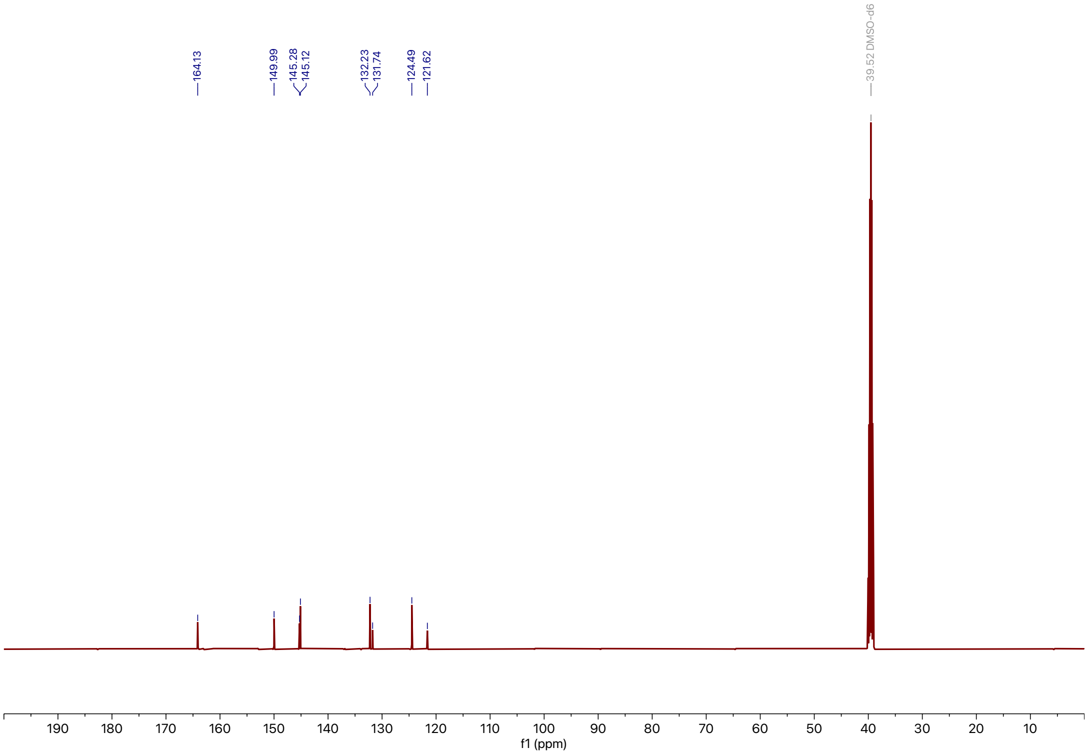

**Supporting Figure 15:** $^{13}$C NMR spectra of **2**

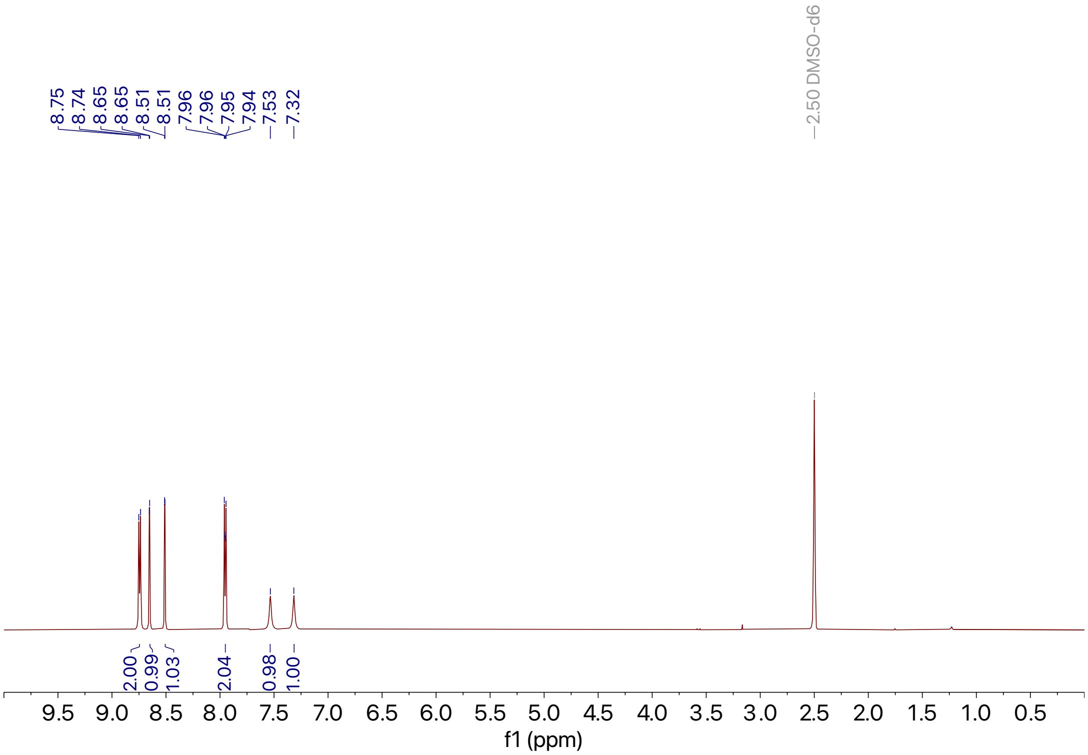

**Supporting Figure 16:** $^{1}$H NMR spectra of **3**

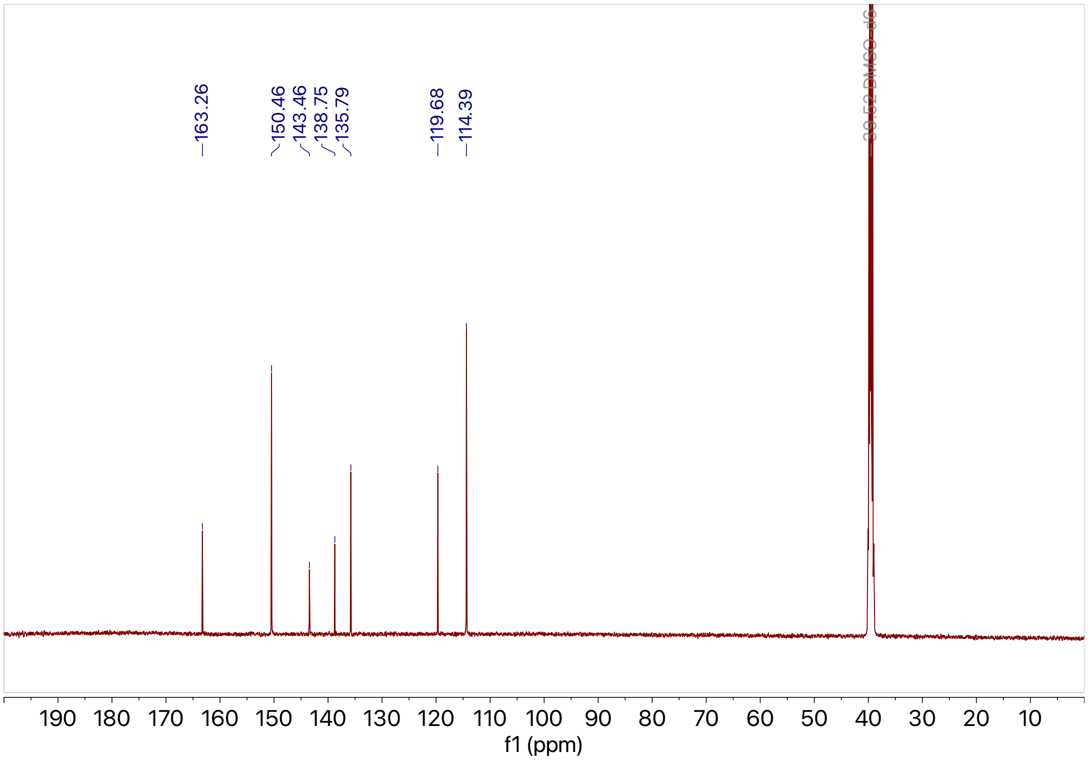

**Supporting Figure 17:** $^{13}$C NMR spectra of **3**

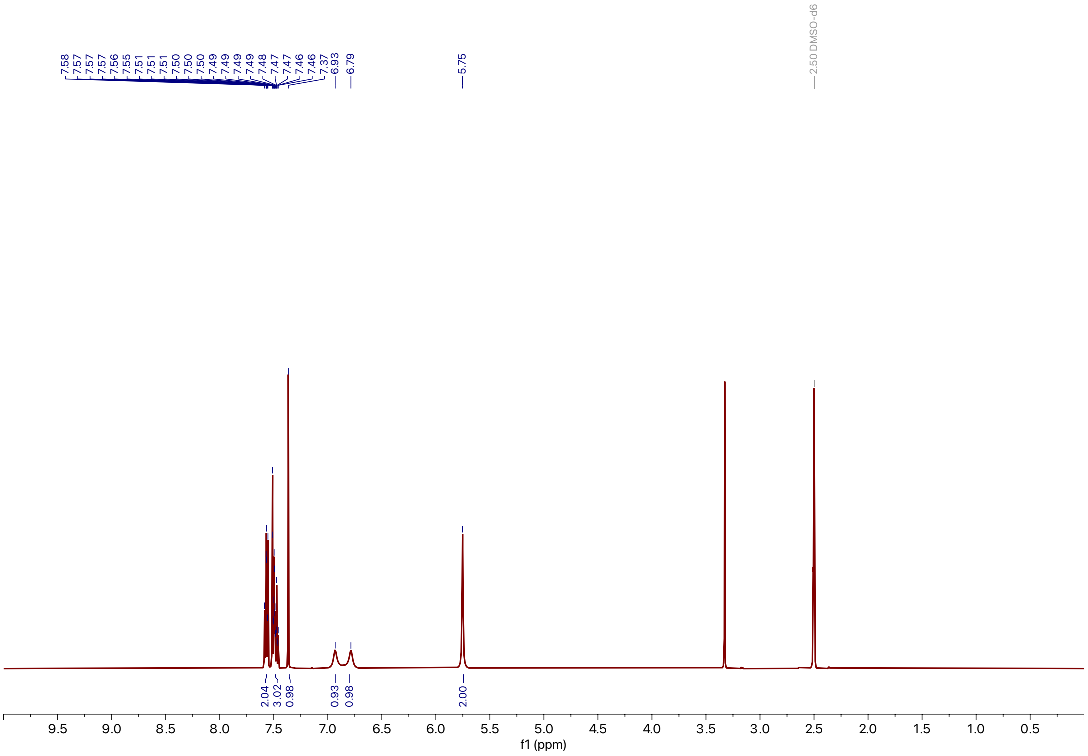

**Supporting Figure 18:** $^{1}$H NMR spectra of **4**

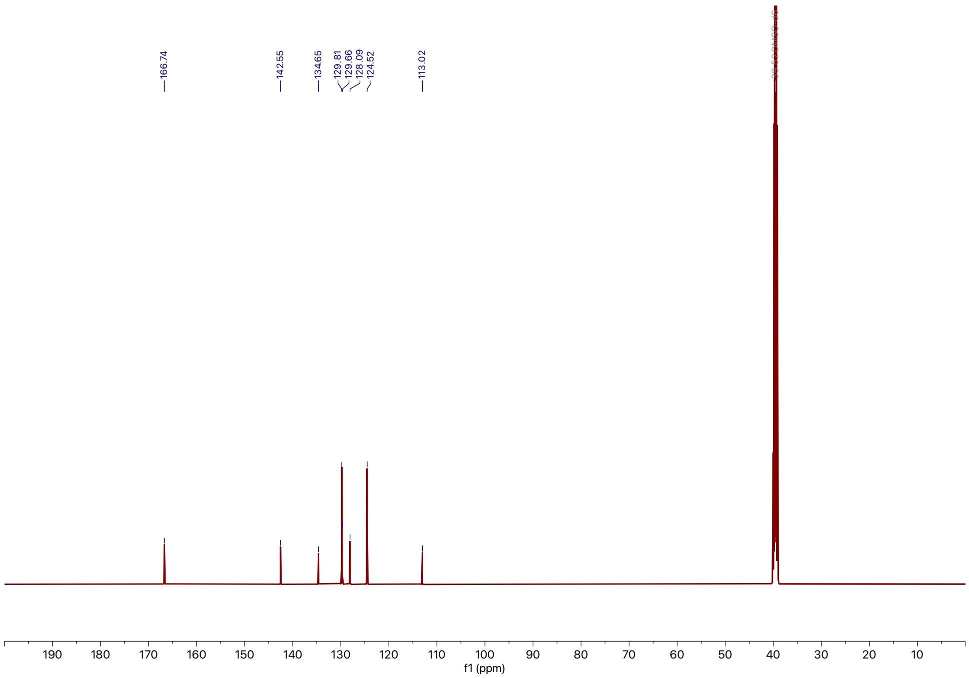

**Supporting Figure 19:** $^{13}$C NMR spectra of **4**

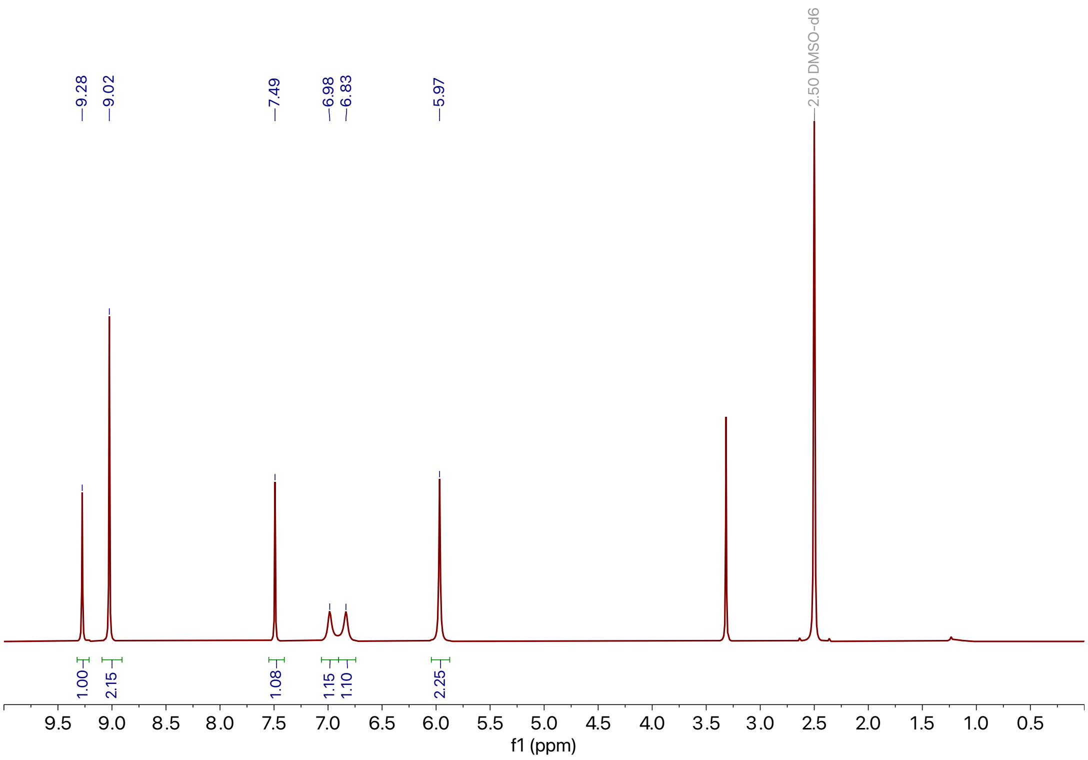

**Supporting Figure 20:** $^{1}$H NMR spectra of **5**

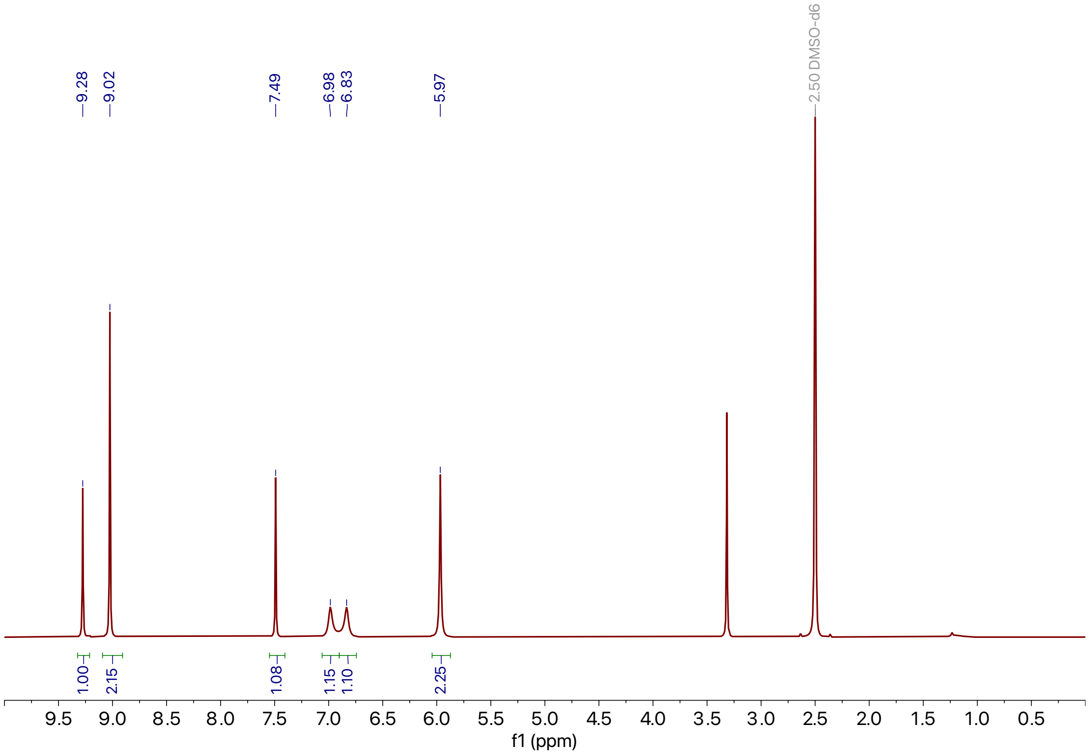

**Supporting Figure 21:** $^{13}$C NMR spectra of **5**

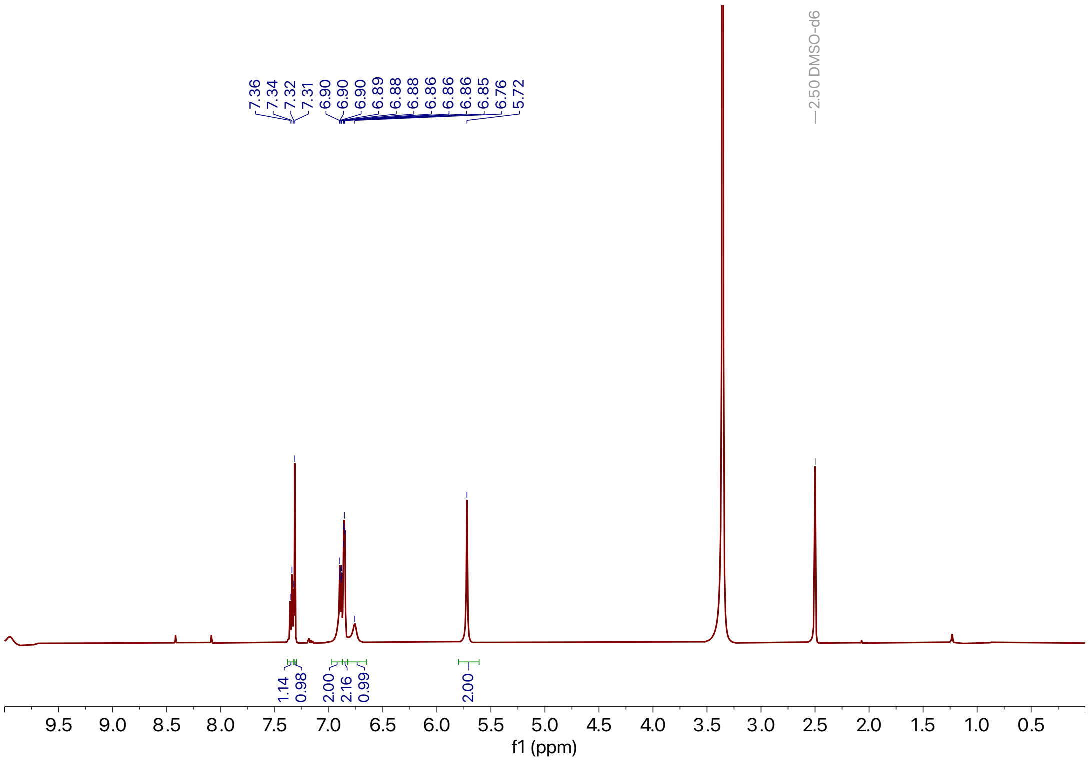

**Supporting Figure 22:** $^{1}$H NMR spectra of **6**

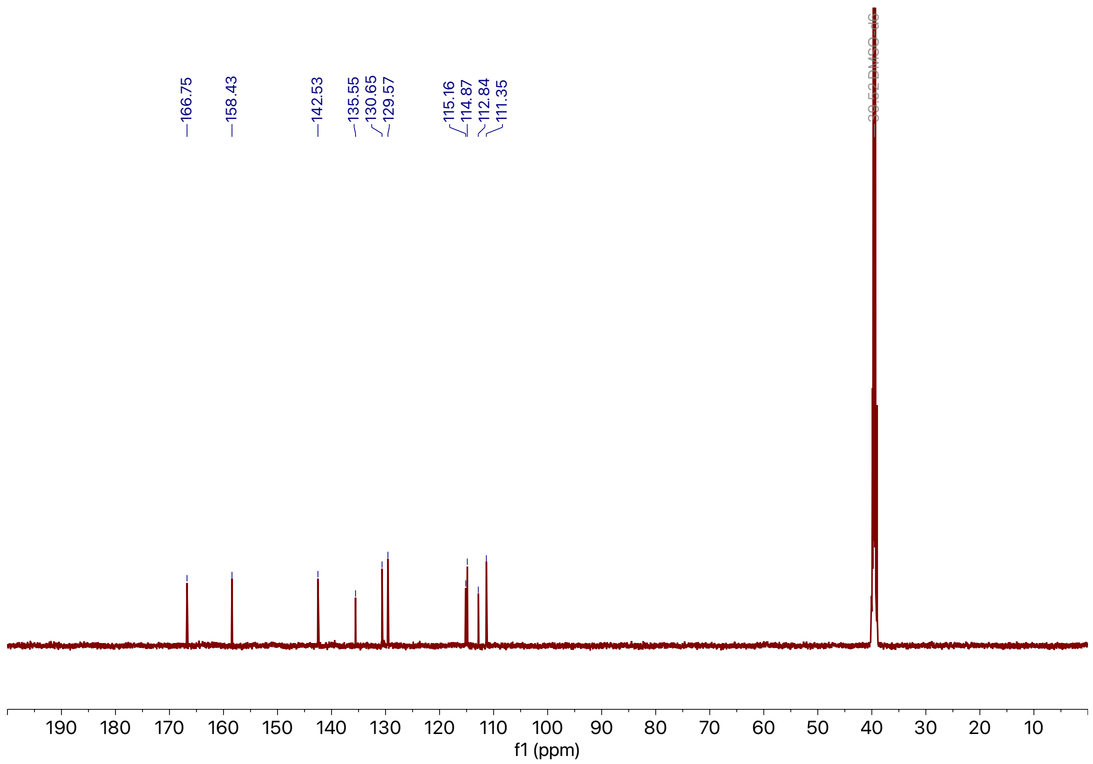

**Supporting Figure 23:** $^{13}$C NMR spectra of **6**

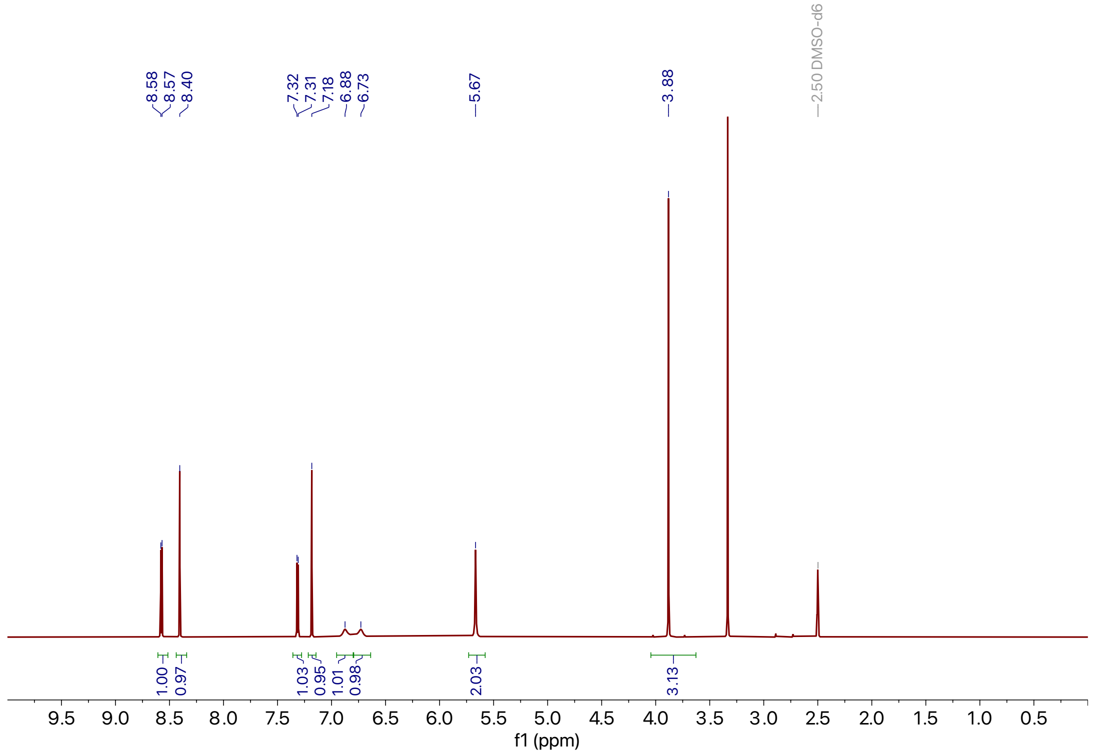

**Supporting Figure 24:** $^{1}$H NMR spectra of **7**

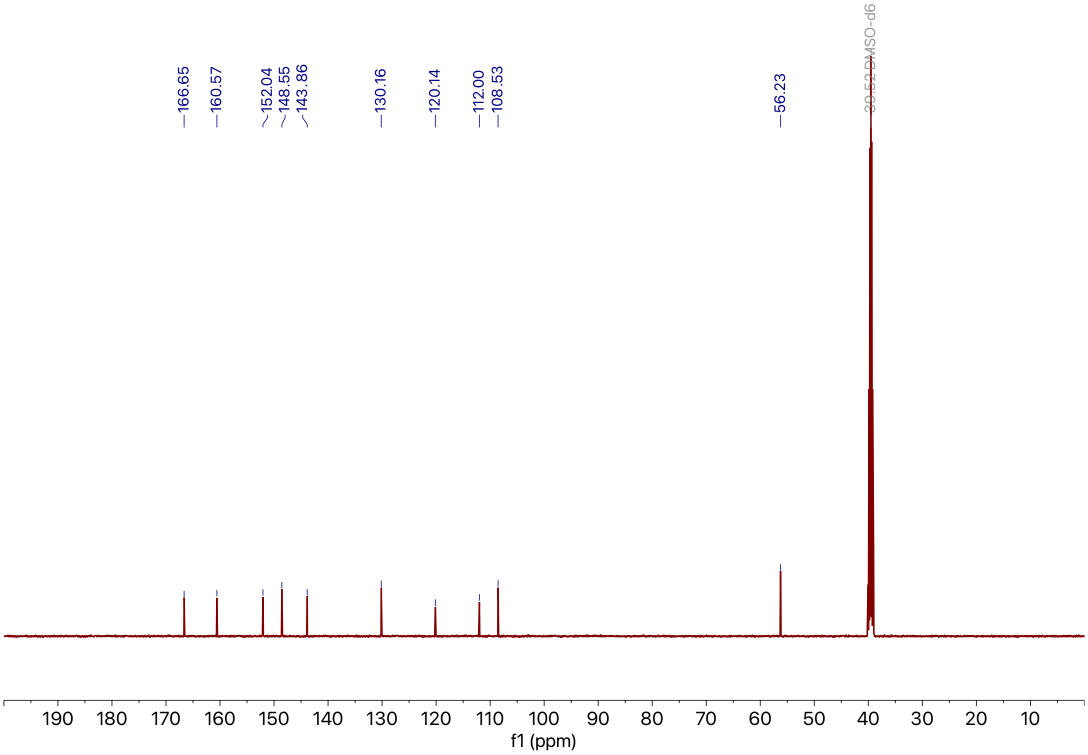

**Supporting Figure 25:** $^{13}$C NMR spectra of **7**

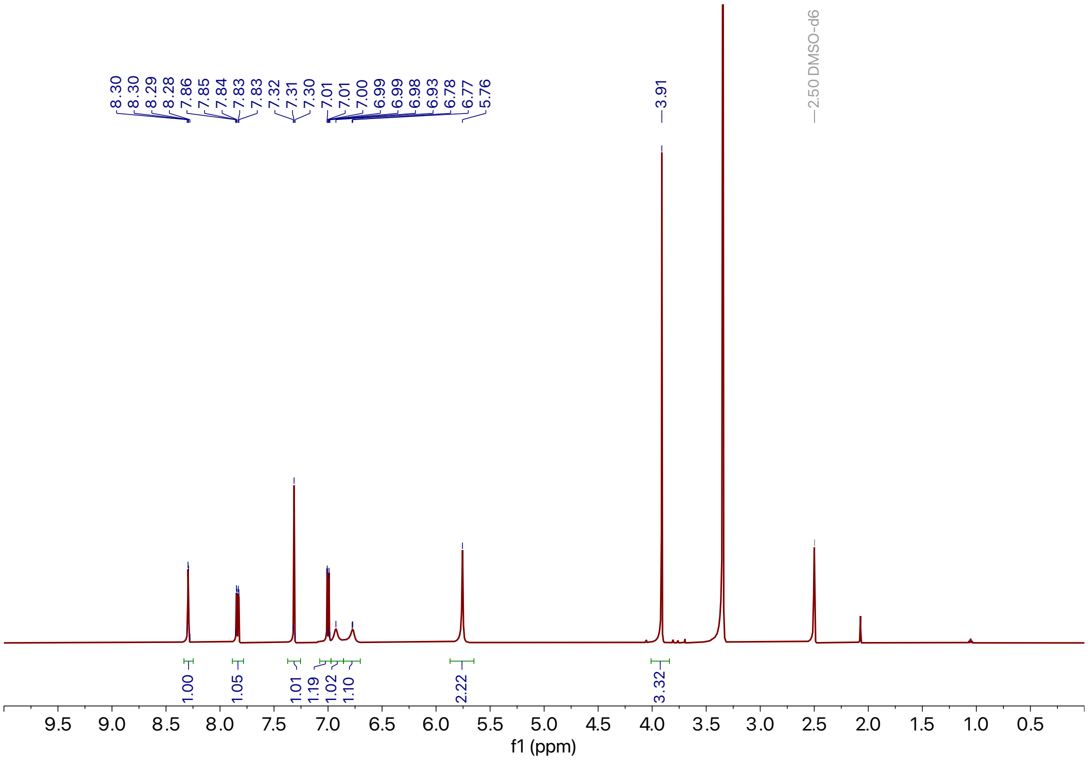

**Supporting Figure 26:** $^{1}$H NMR spectra of **8**

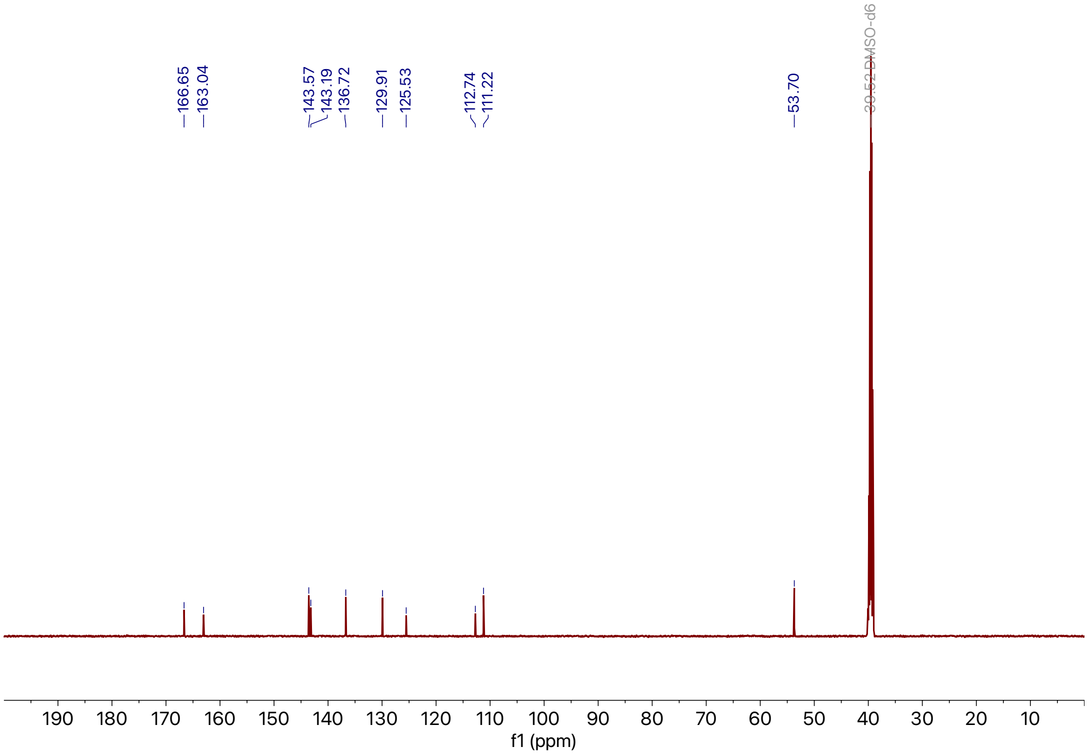

**Supporting Figure 27:** $^{13}$C NMR spectra of **8**

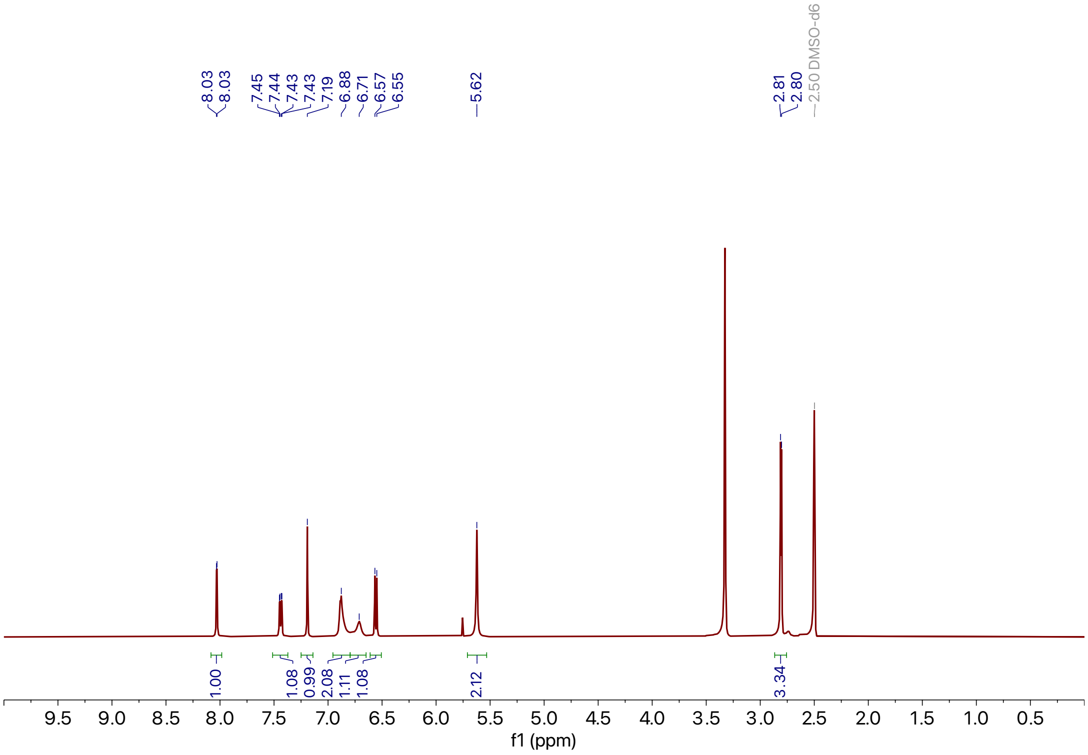

**Supporting Figure 28:** $^{1}$H NMR spectra of **9**

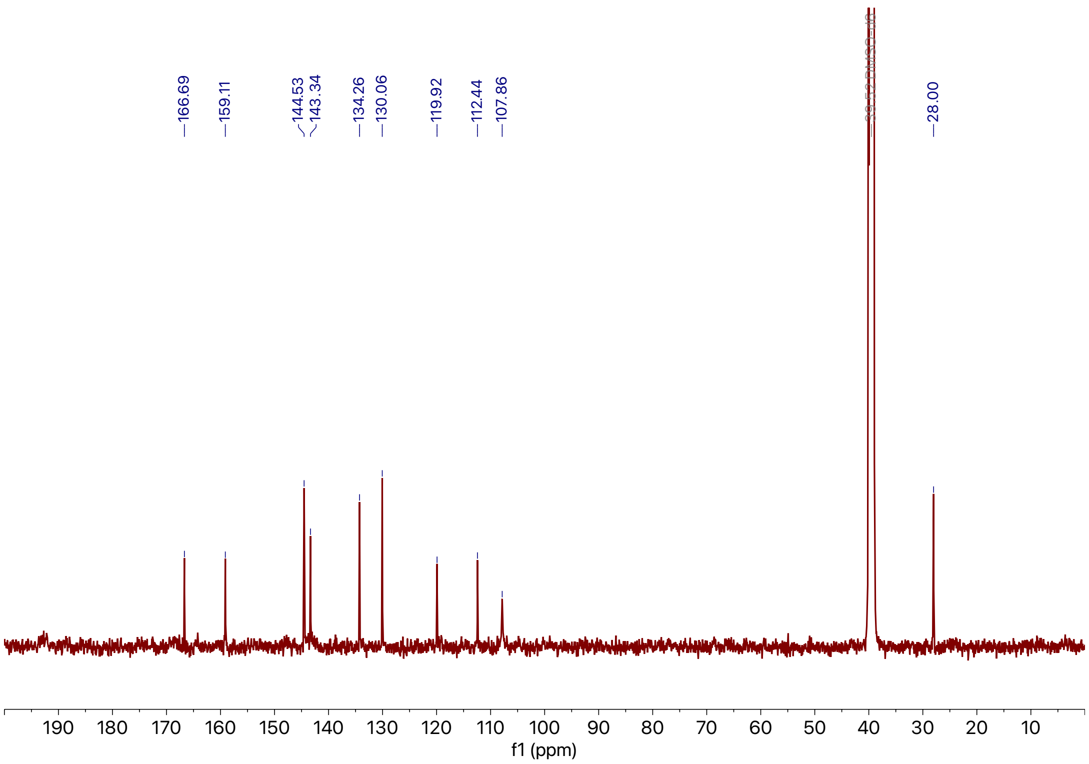

**Supporting Figure 29:** $^{13}$C NMR spectra of **9**

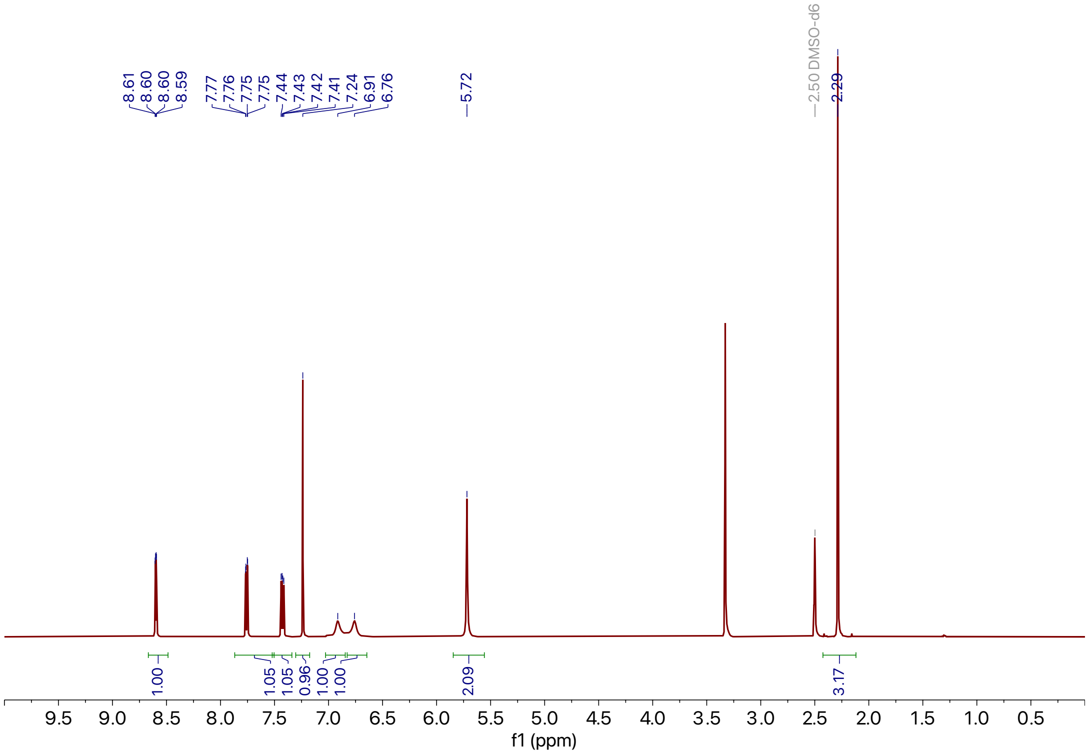

**Supporting Figure 30:** $^{1}$H NMR spectra of **10**

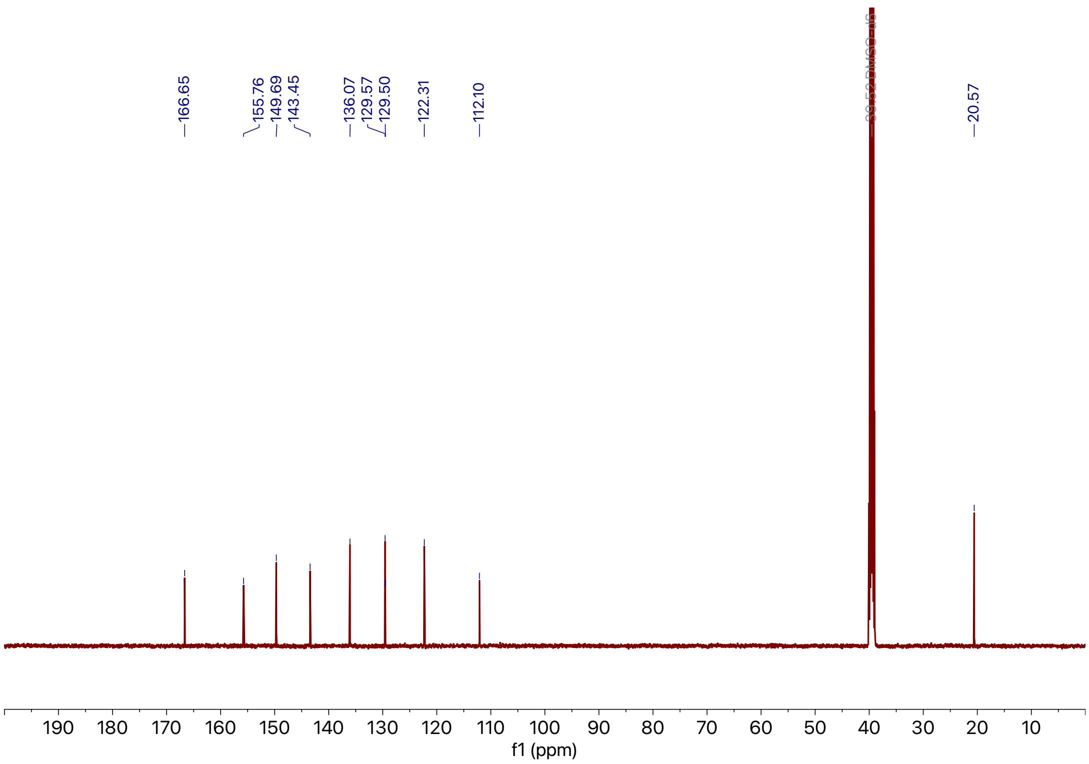

**Supporting Figure 31:** $^{13}$C NMR spectra of **10**

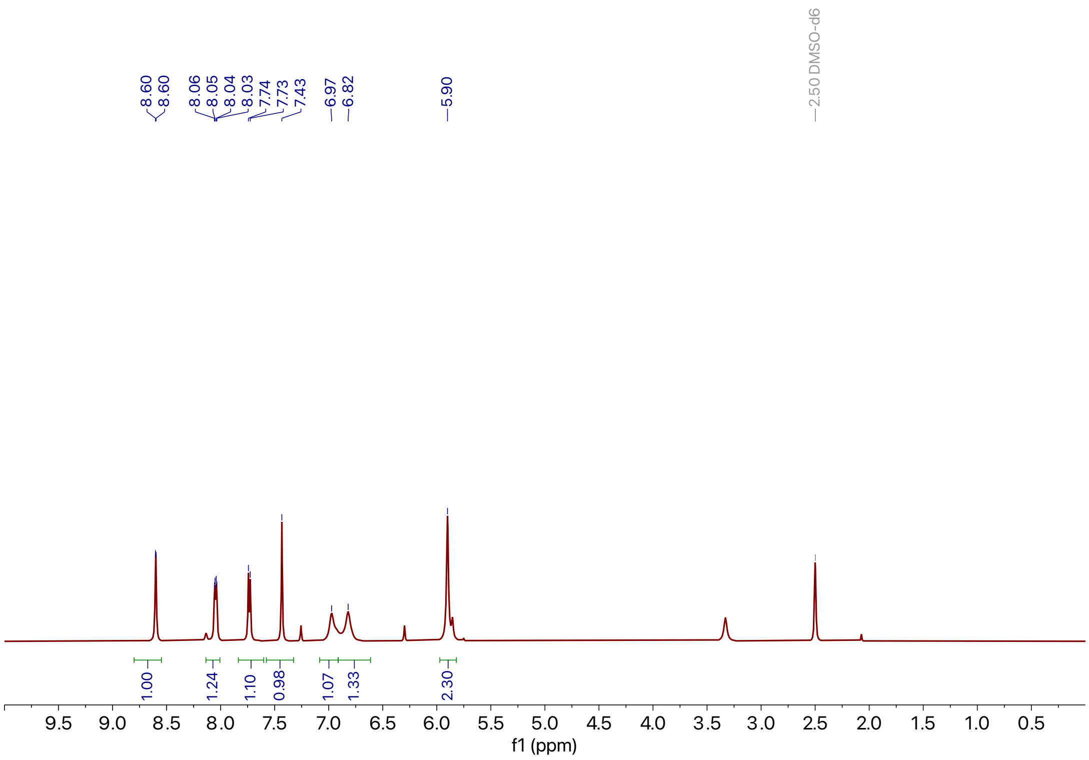

**Supporting Figure 32:** $^1$H NMR spectra of **11**

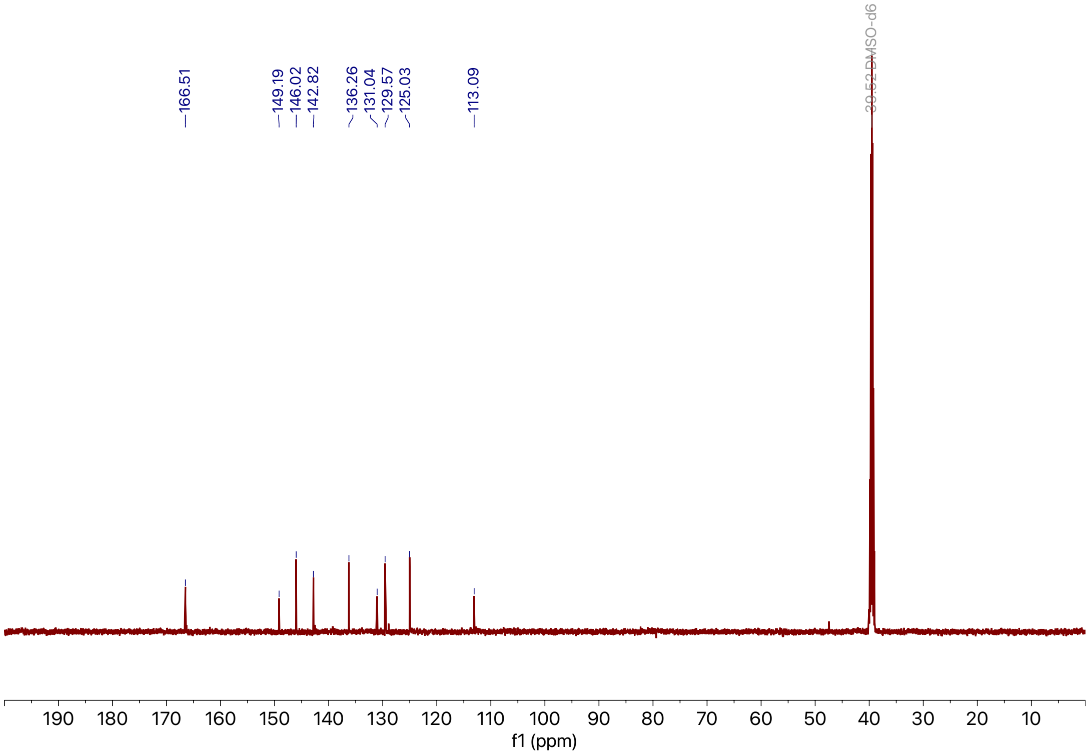

**Supporting Figure 33:** $^{13}$C NMR spectra of **11**

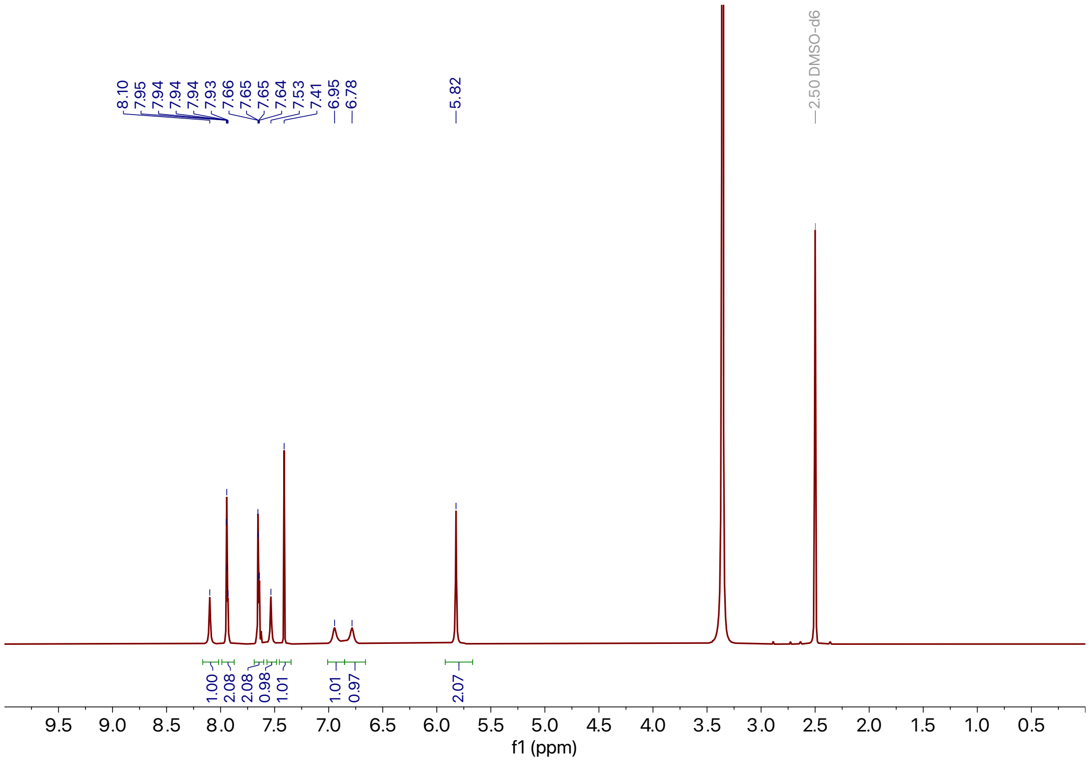

**Supporting Figure 34:** $^{1}$H NMR spectra of **12**

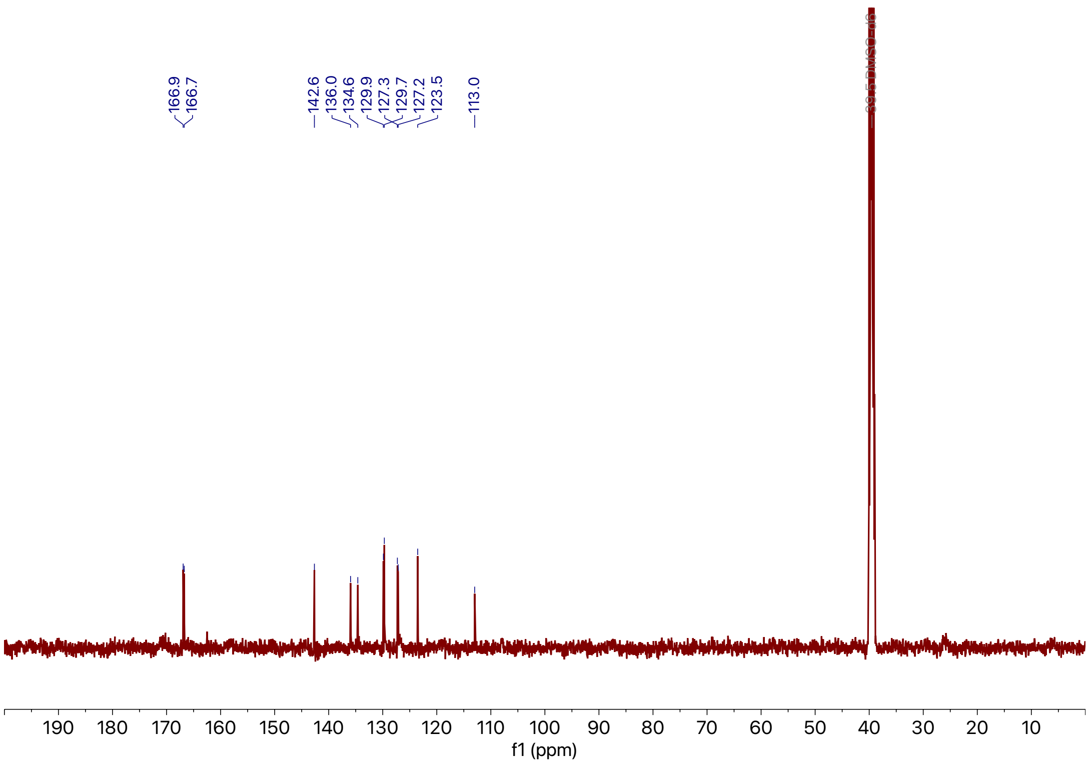

**Supporting Figure 35:** $^{13}$C NMR spectra of **12**

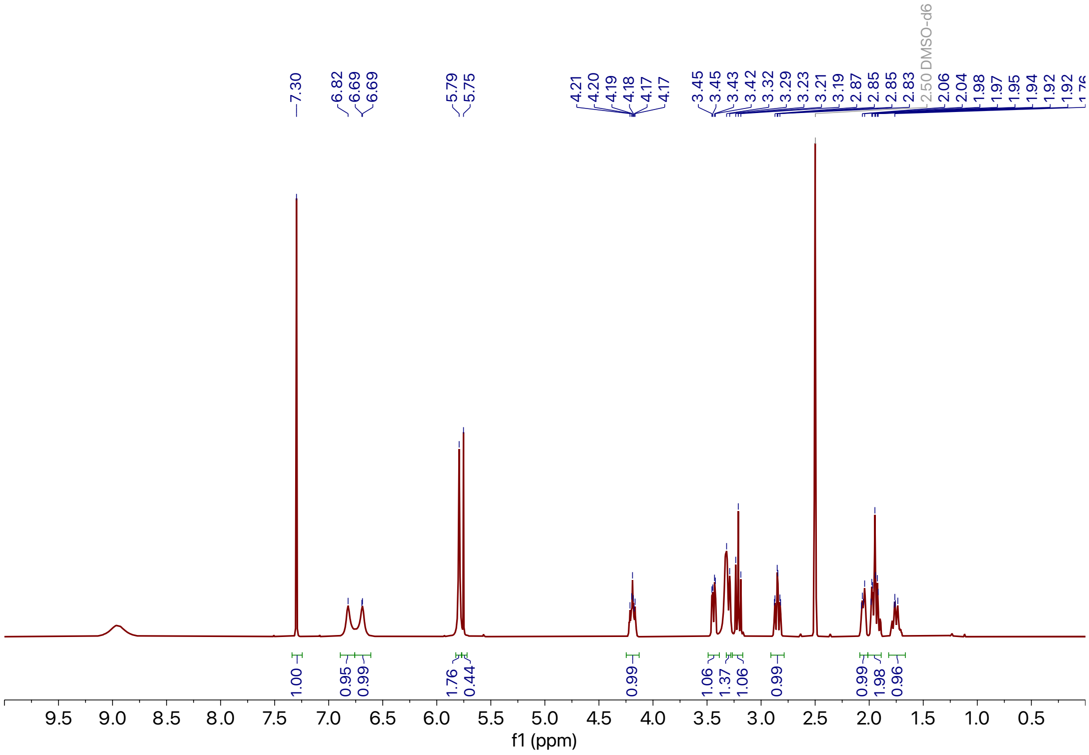

**Supporting Figure 36:** $^{1}$H NMR spectra of **13**

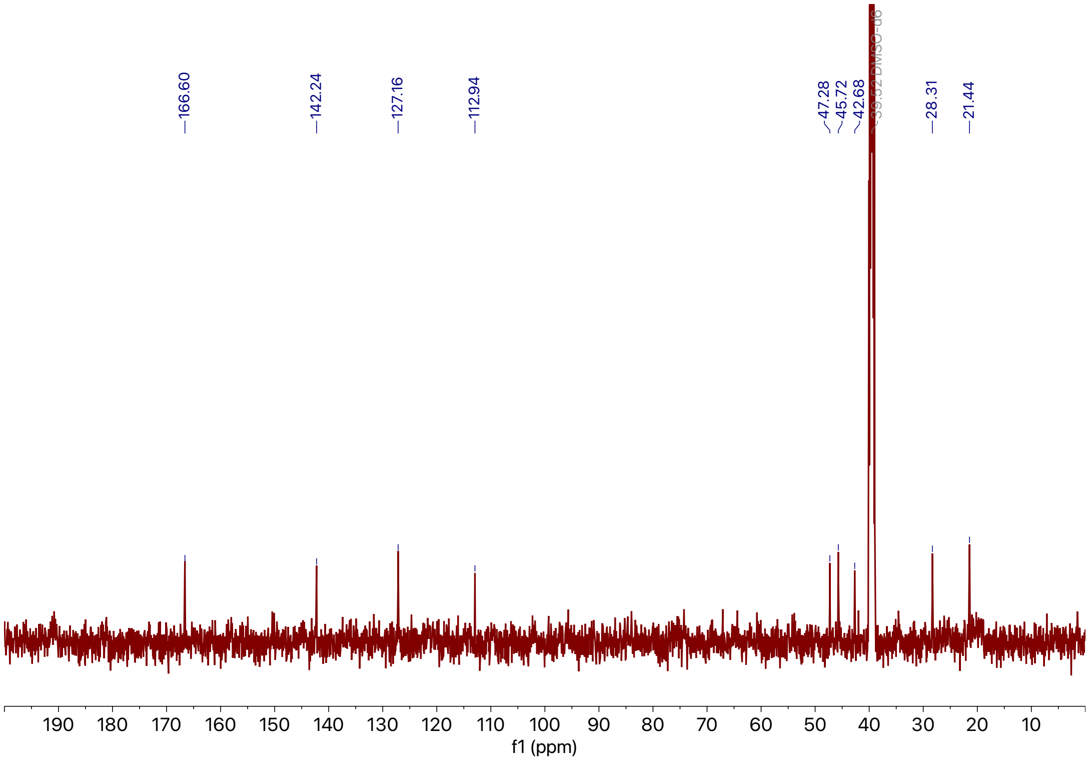

**Supporting Figure 37:** $^{13}$C NMR spectra of **13**

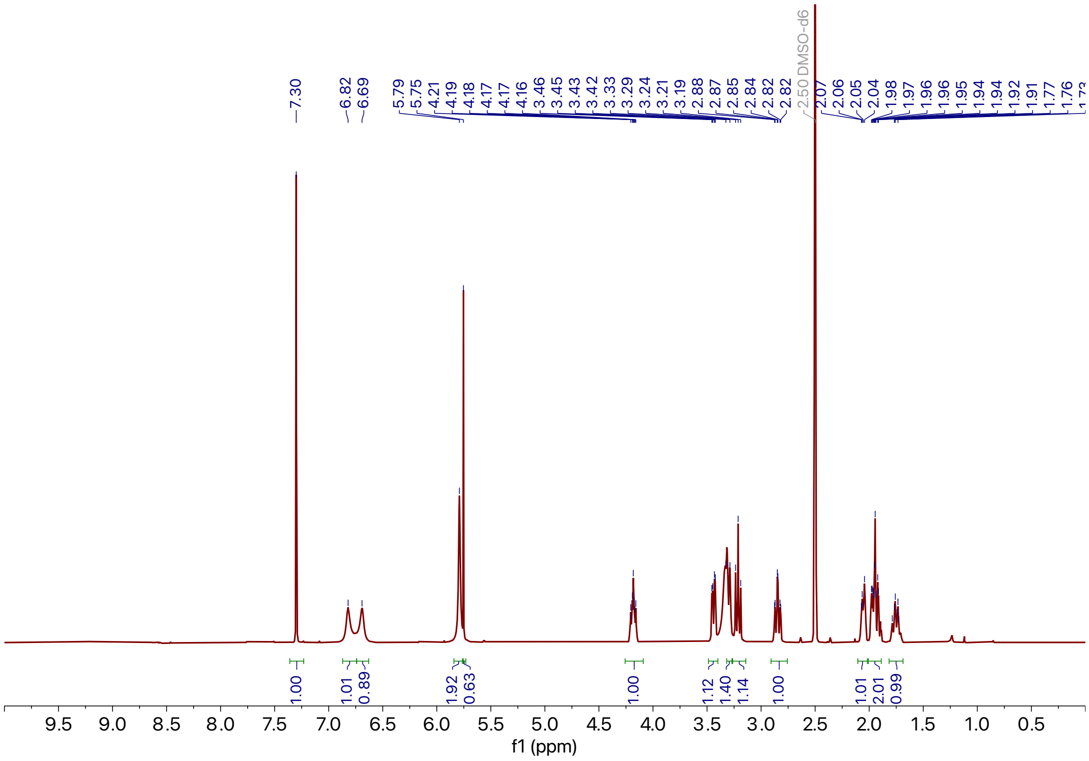

**Supporting Figure 38:** $^{1}$H NMR spectra of **14**

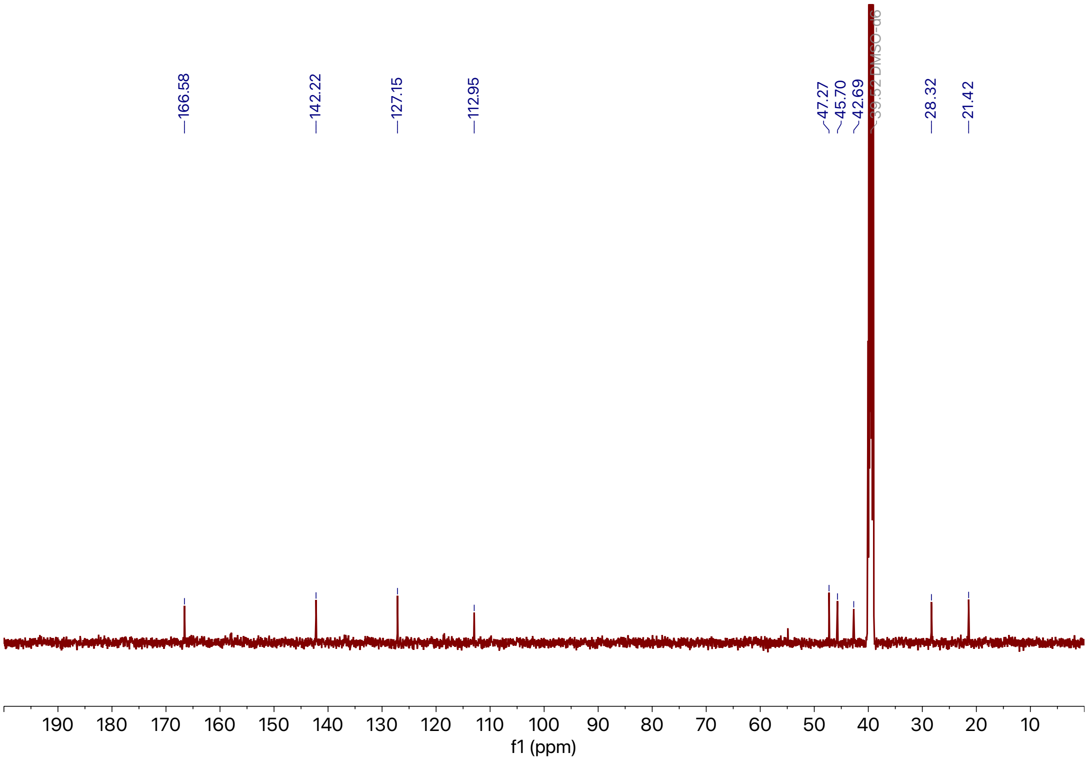

**Supporting Figure 39:** $^{13}$C NMR spectra of **14**

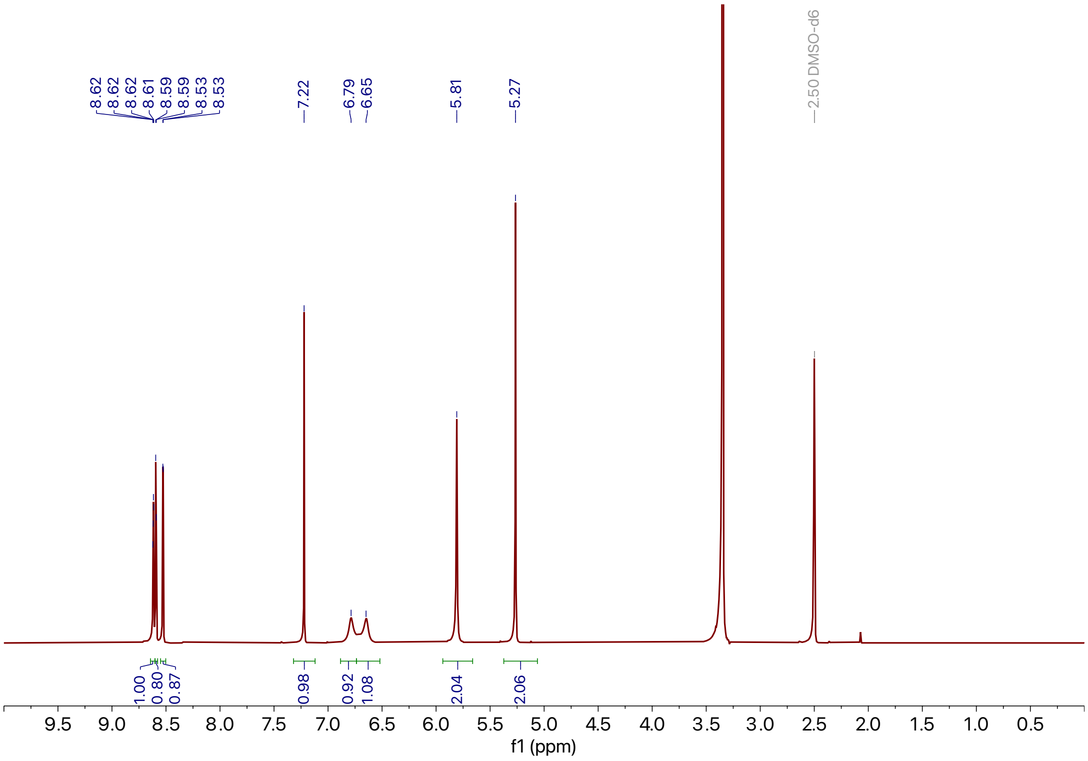

**Supporting Figure 40:** $^{1}$H NMR spectra of **15**

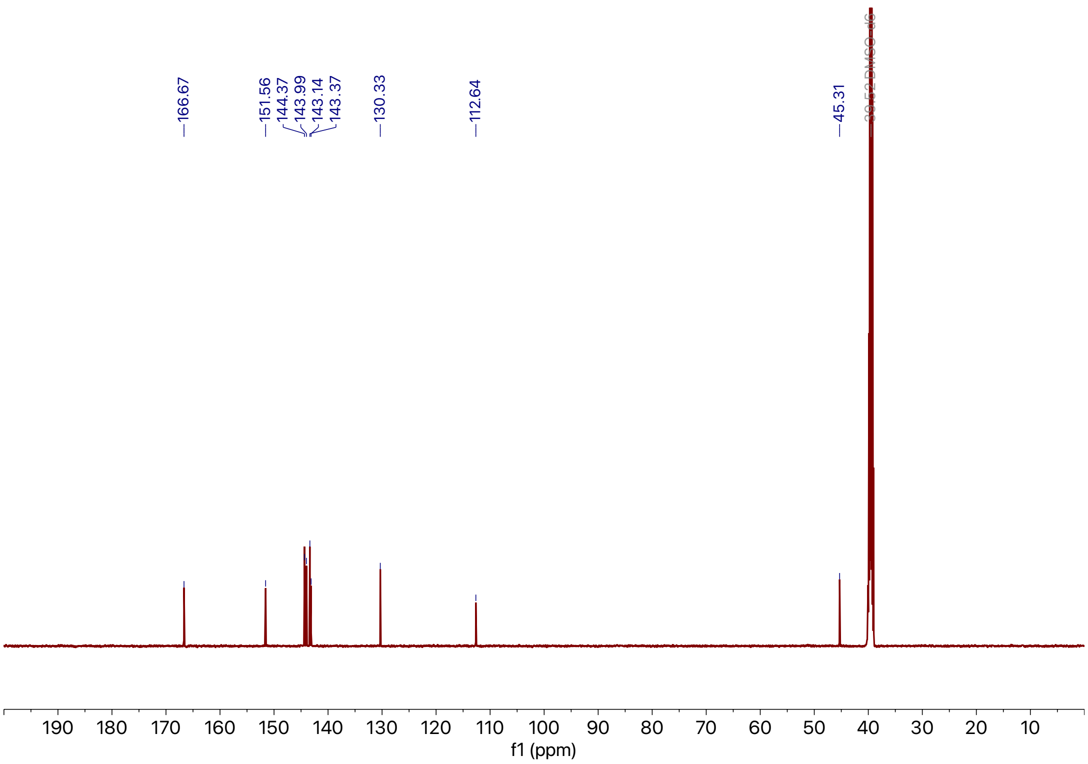

**Supporting Figure 41:** $^{13}$C NMR spectra of **15**

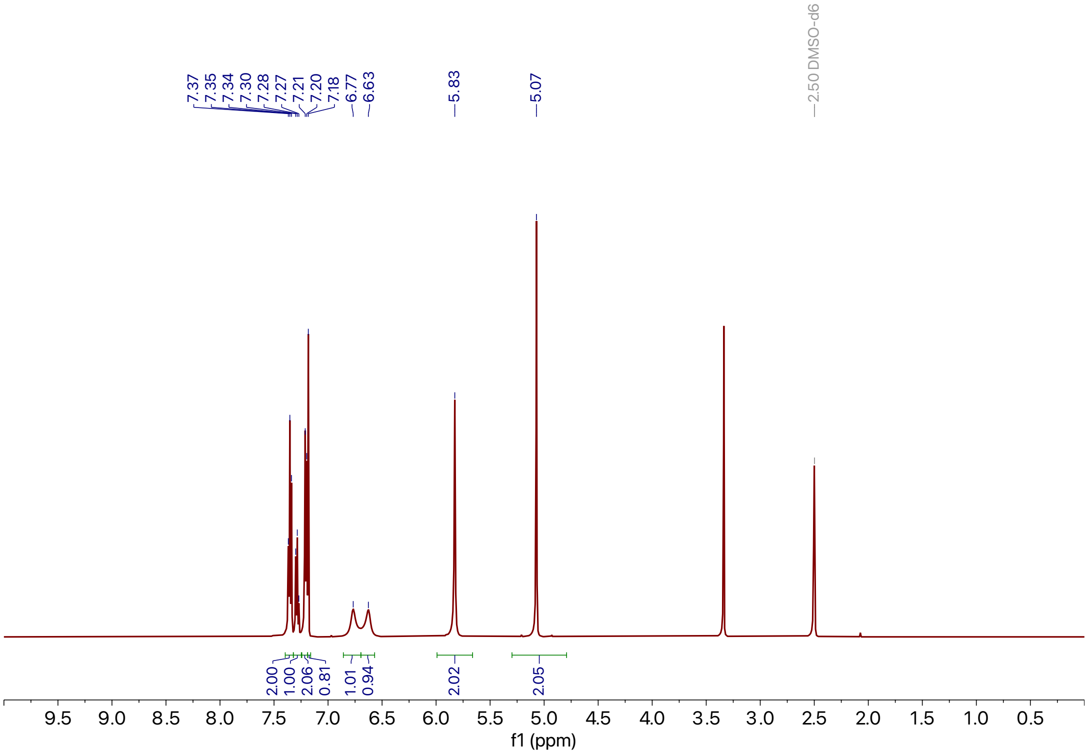

**Supporting Figure 42:** $^{1}$H NMR spectra of **16**

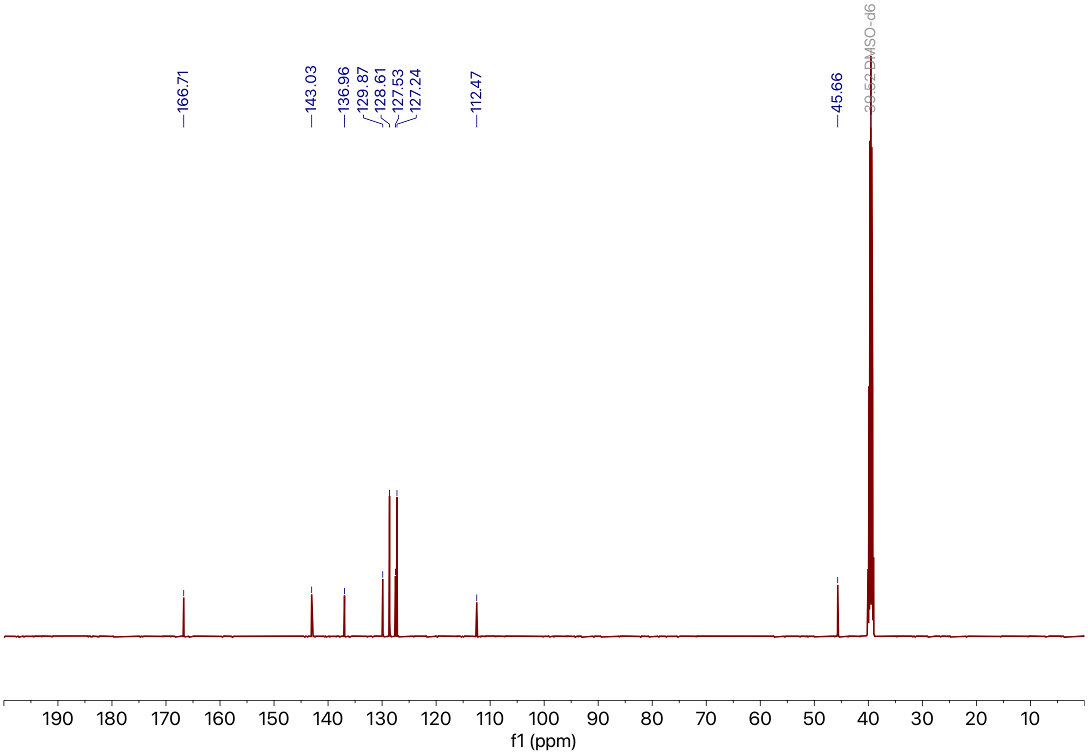

**Supporting Figure 43:** $^{13}$C NMR spectra of **16**

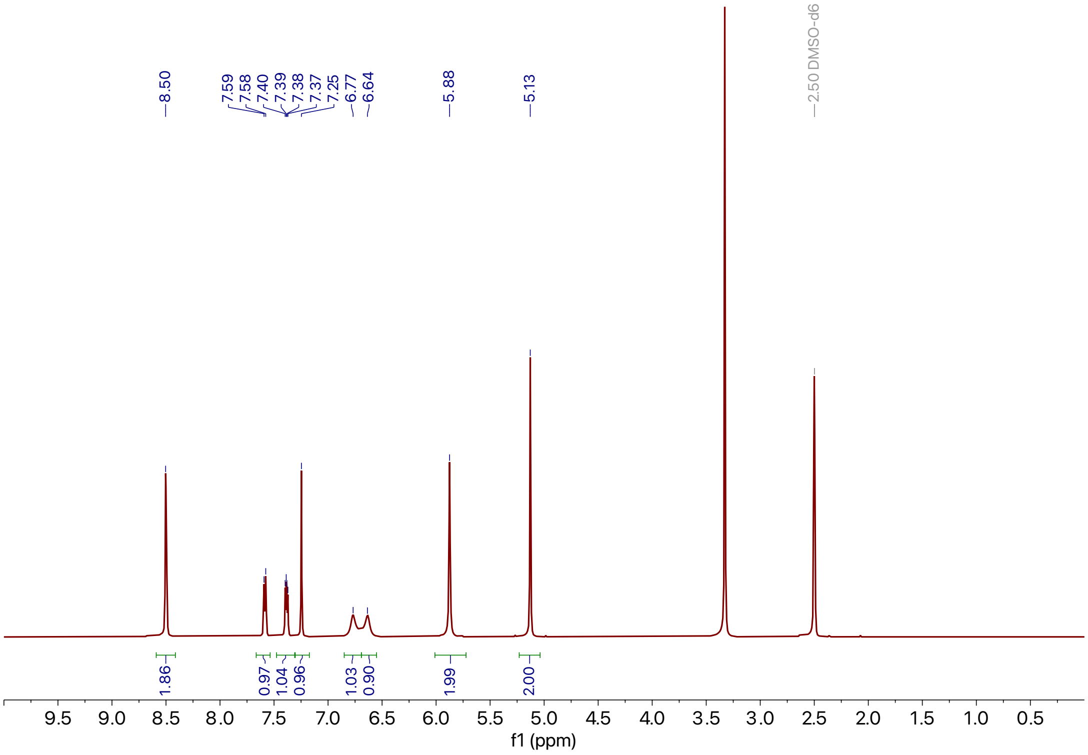

**Supporting Figure 44:** $^{1}$H NMR spectra of **17**

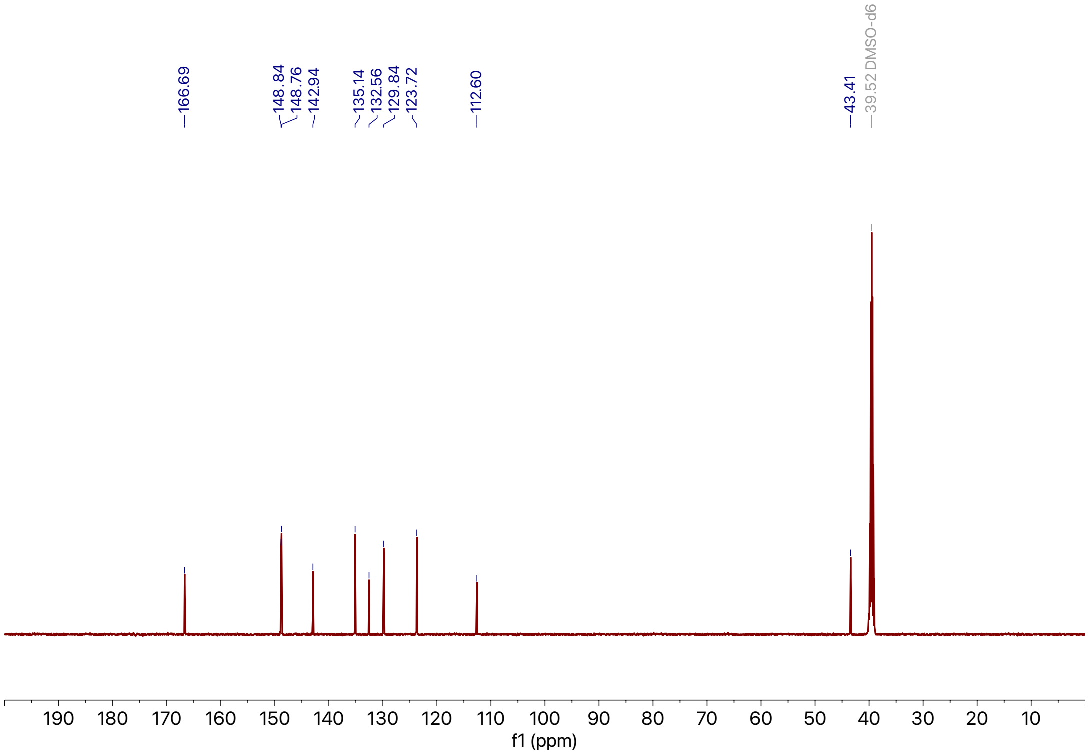

**Supporting Figure 45:** $^{13}$C NMR spectra of **17**

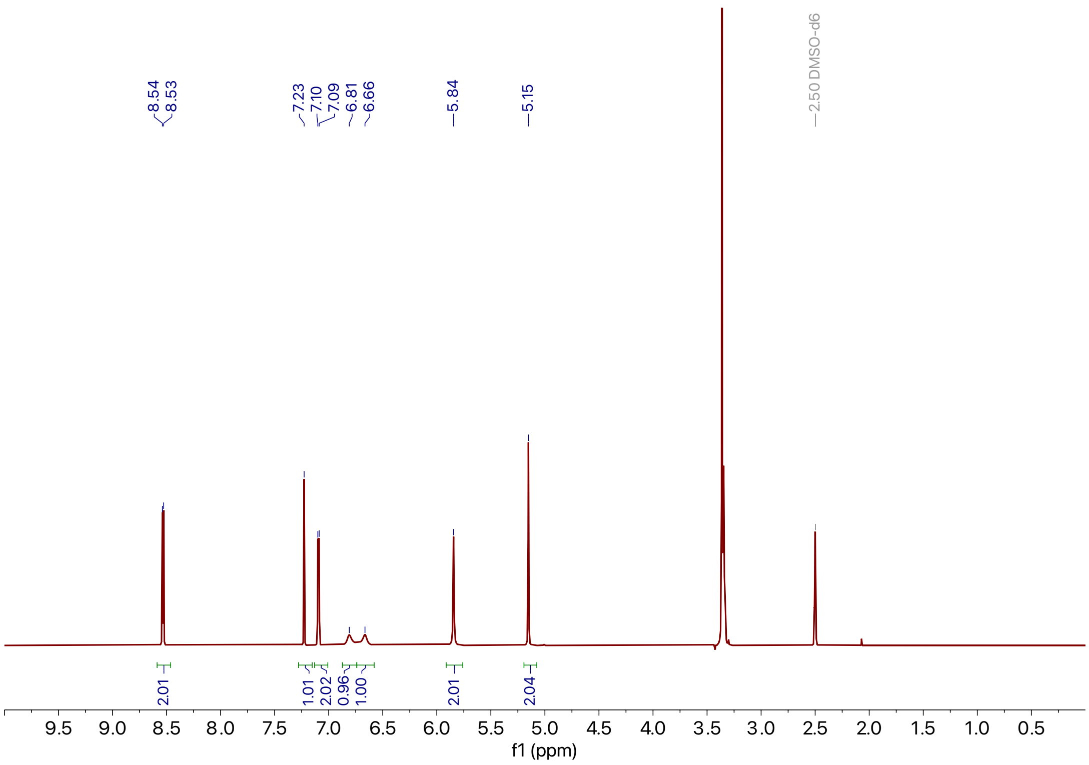

**Supporting Figure 46:** $^{1}$H NMR spectra of **18**

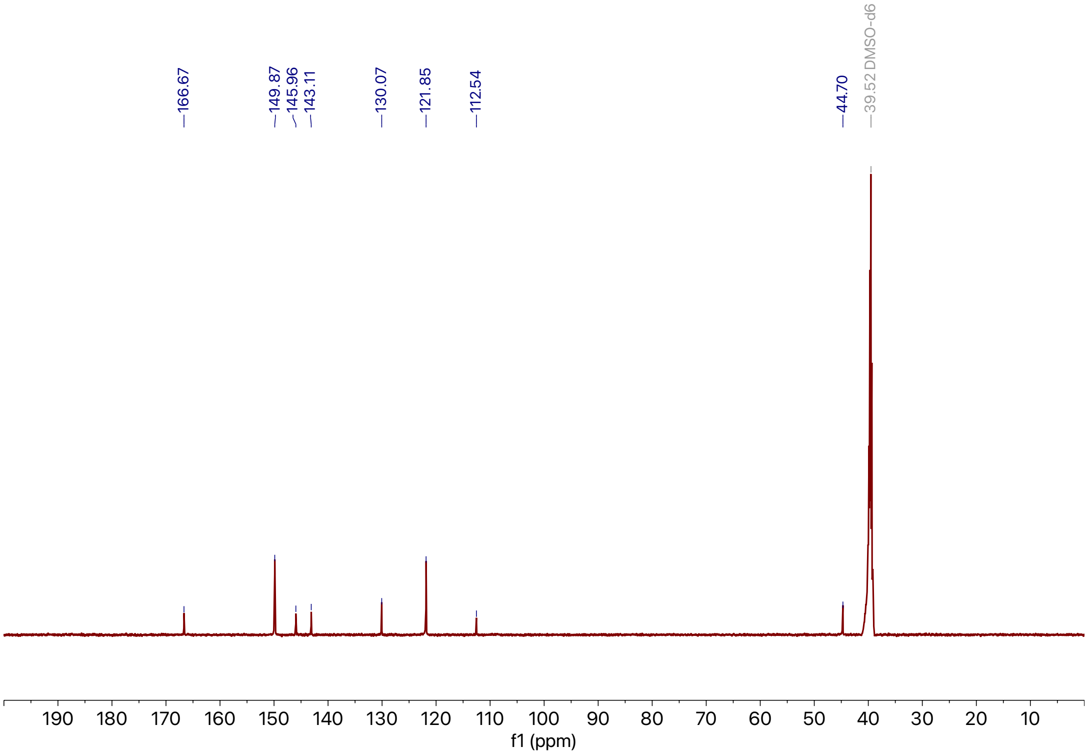

**Supporting Figure 47:** $^{13}$C NMR spectra of **18**

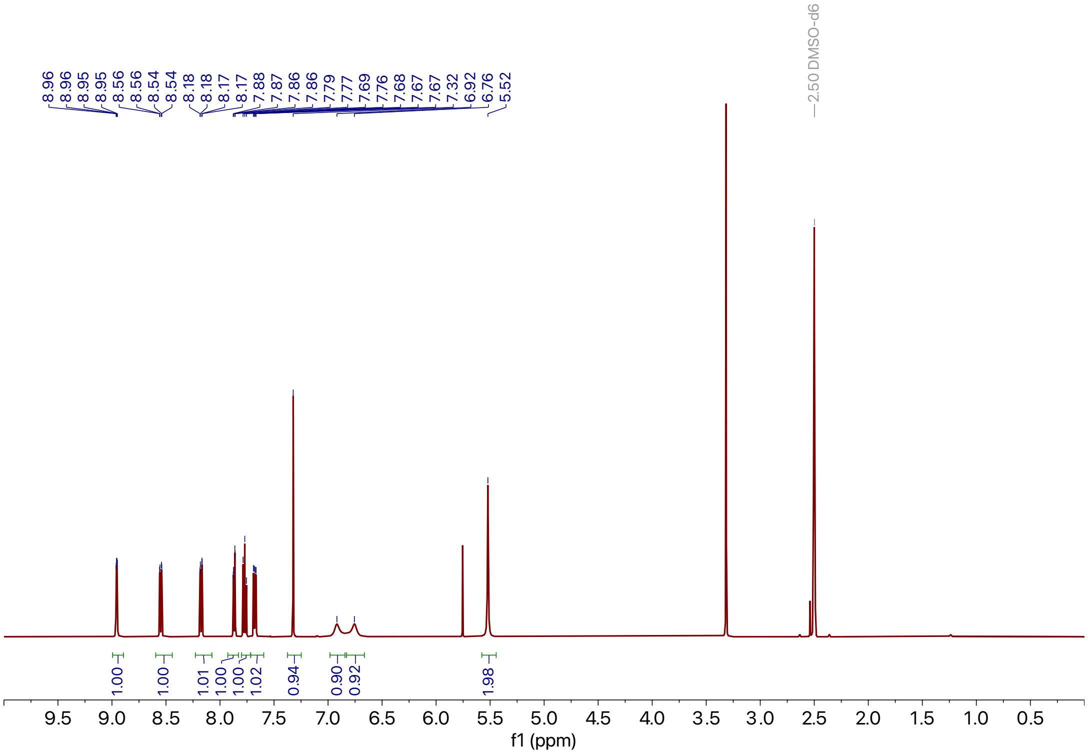

**Supporting Figure 48:** $^{1}$H NMR spectra of **19**

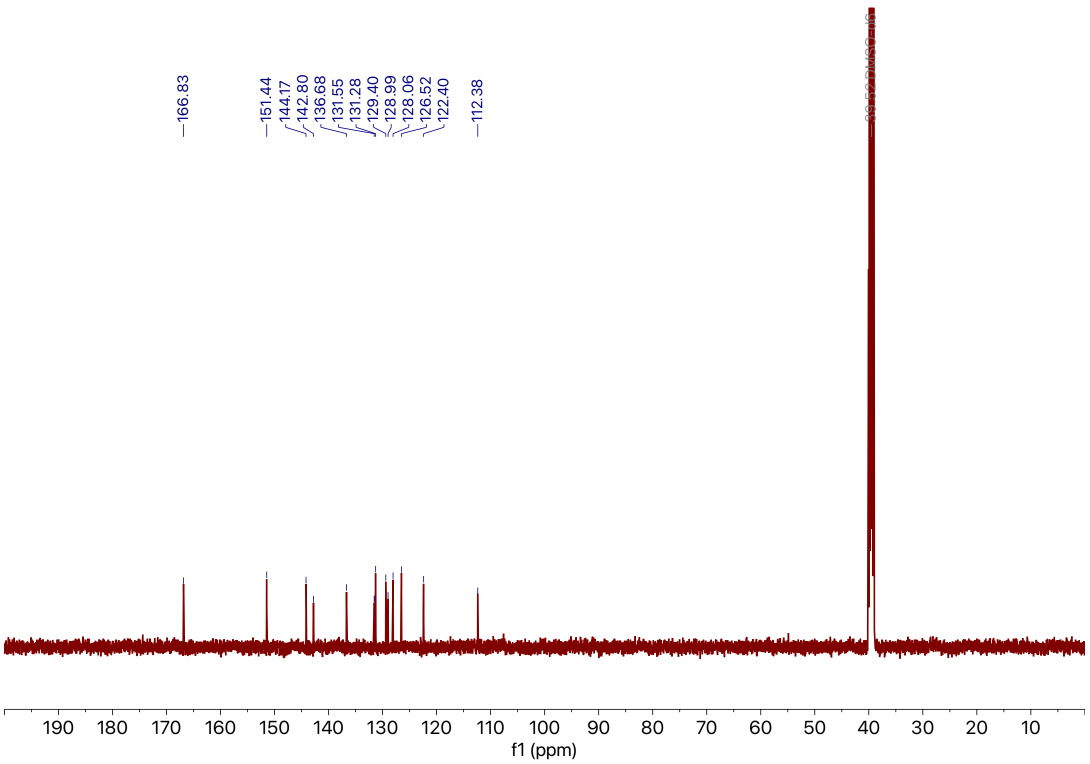

**Supporting Figure 49:** $^{13}$C NMR spectra of **19**

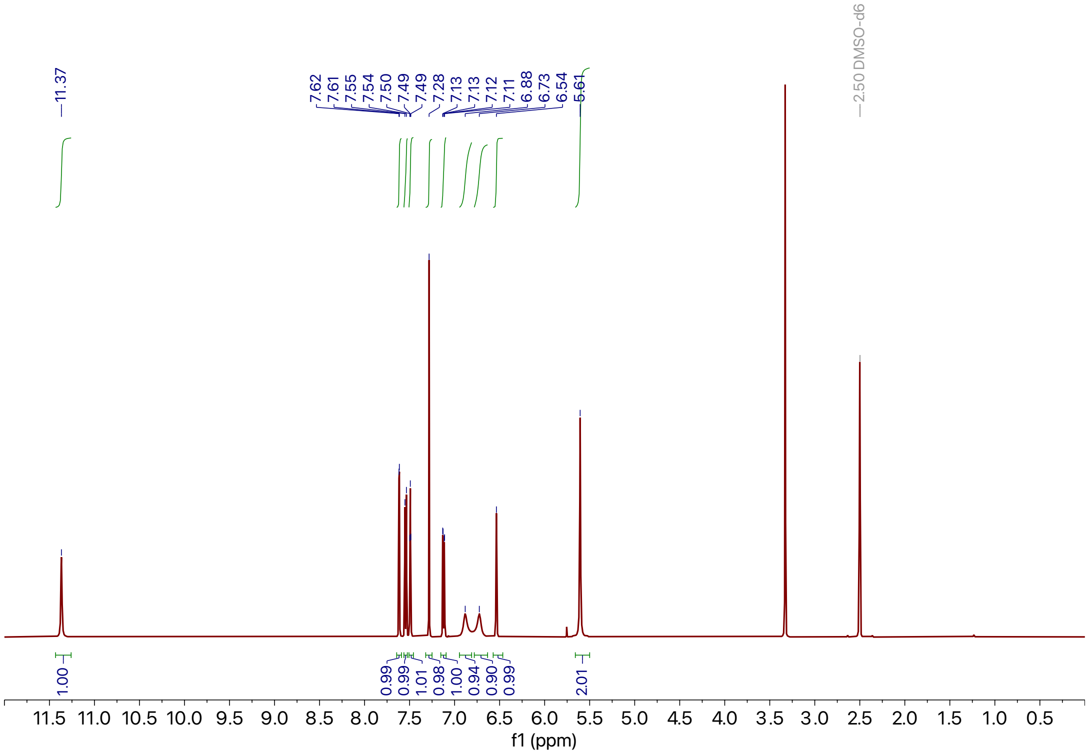

**Supporting Figure 50:** $^{1}$H NMR spectra of **20**

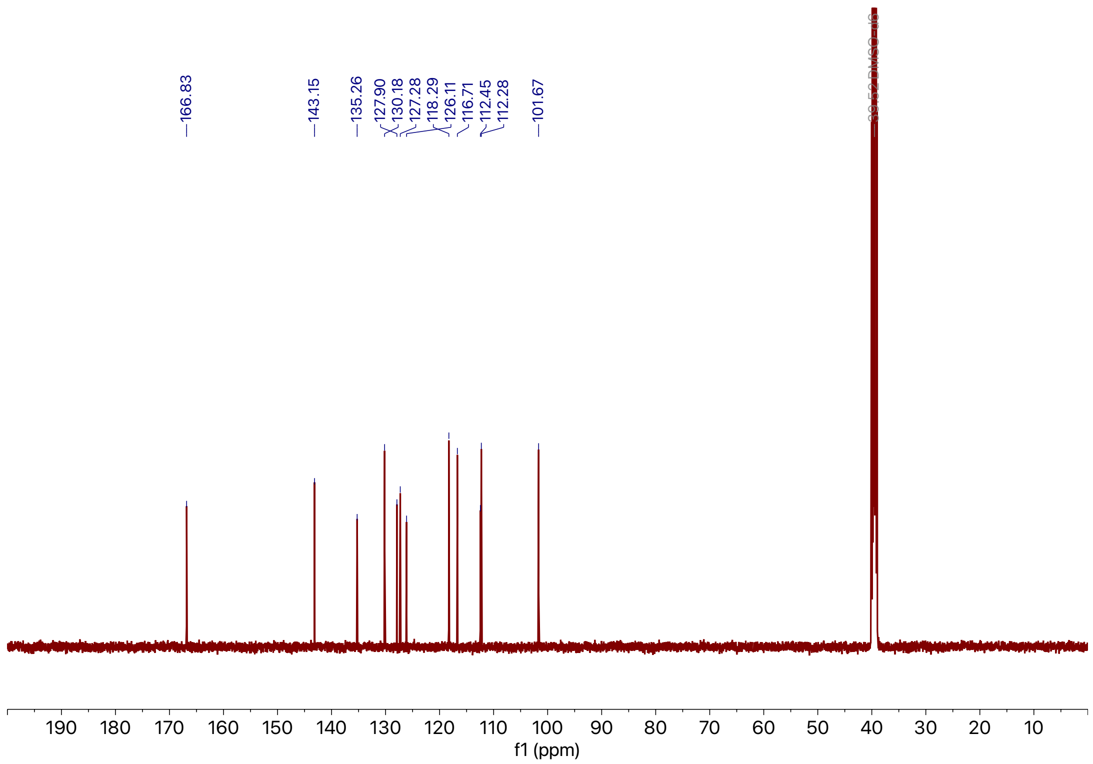

**Supporting Figure 51:** $^{13}$C NMR spectra of **20**

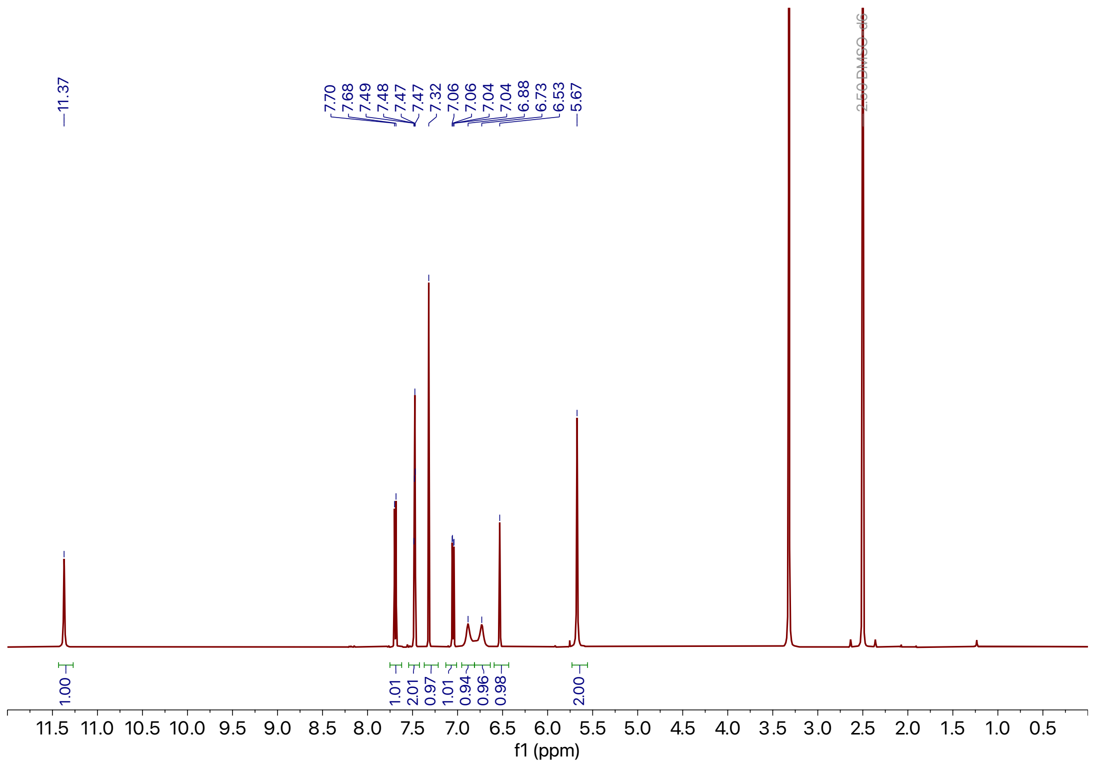

**Supporting Figure 52:** $^{1}$H NMR spectra of **21**

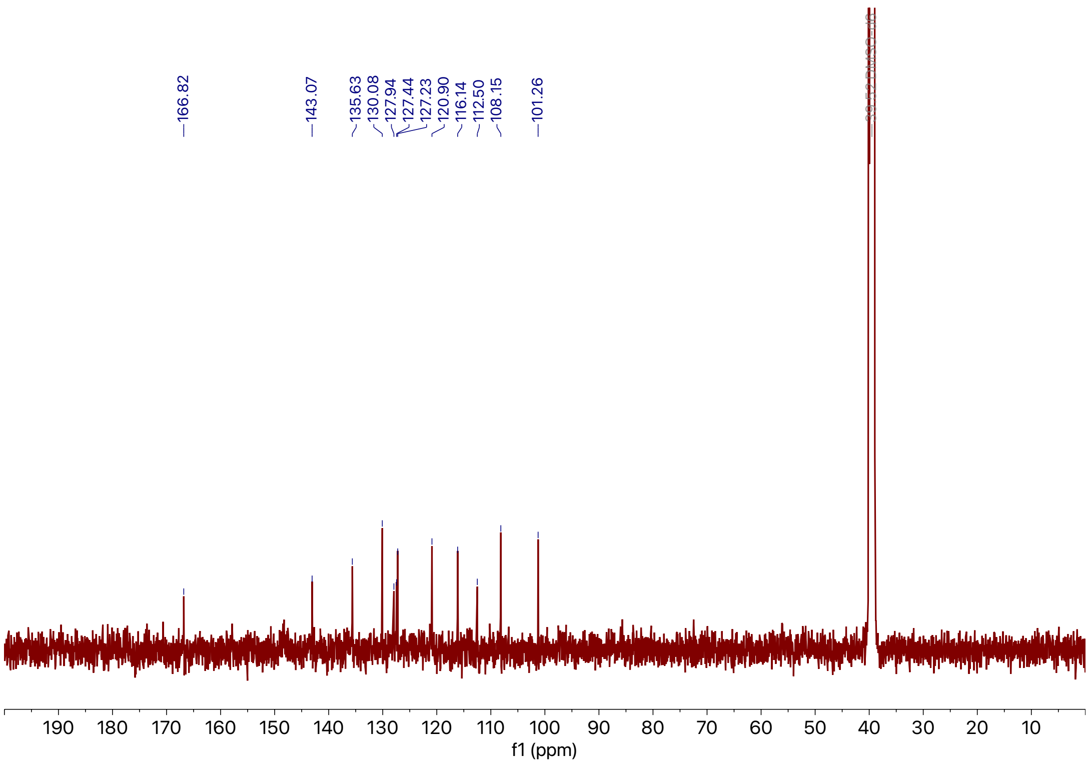

**Supporting Figure 53:** $^{13}$C NMR spectra of **21**

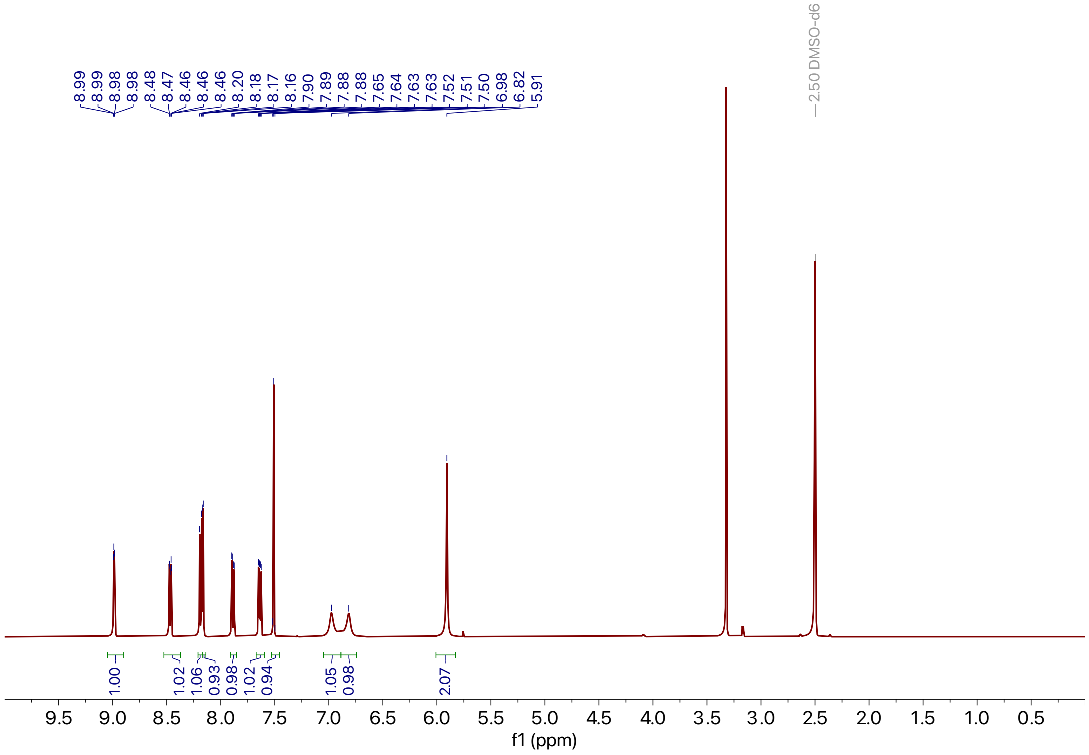

**Supporting Figure 54:** $^{1}$H NMR spectra of **22**

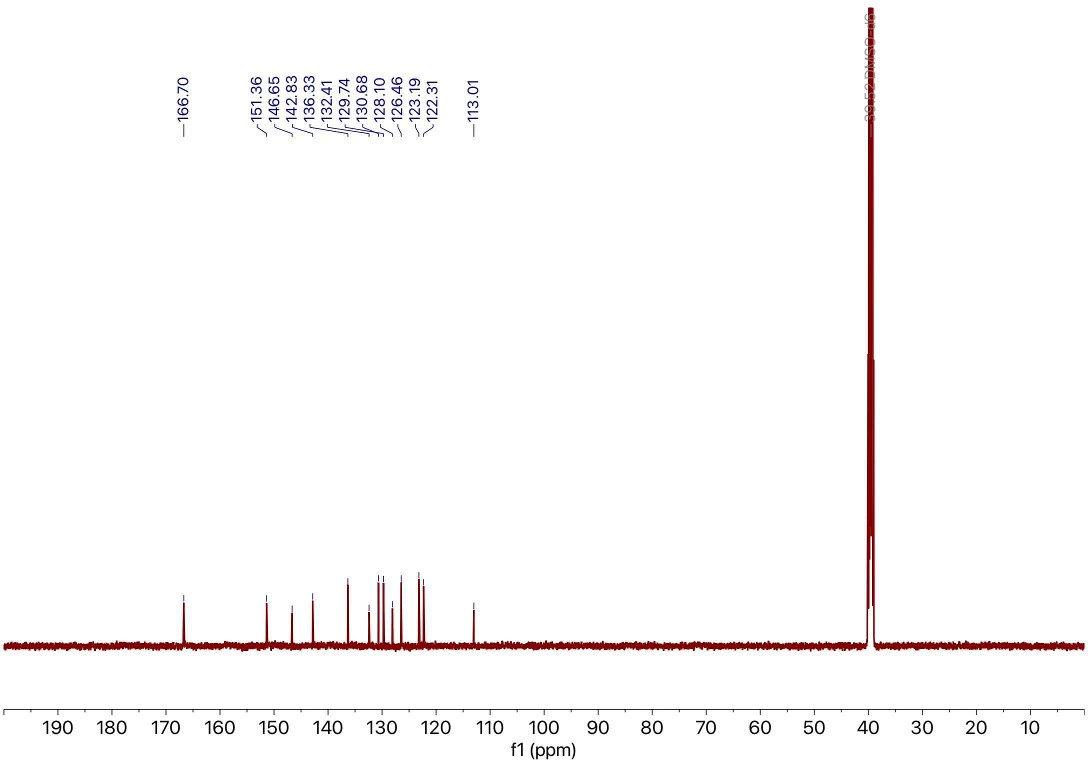

**Supporting Figure 55:** $^{13}$C NMR spectra of **22**

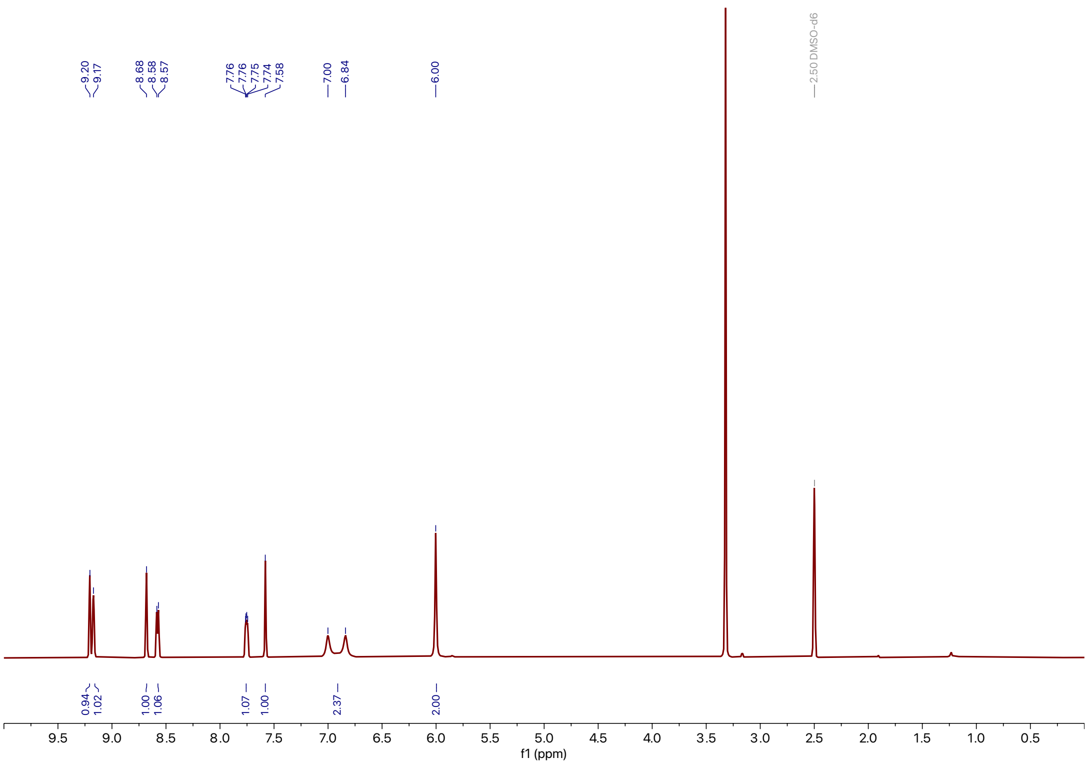

**Supporting Figure 56:** $^{1}$H NMR spectra of **23**

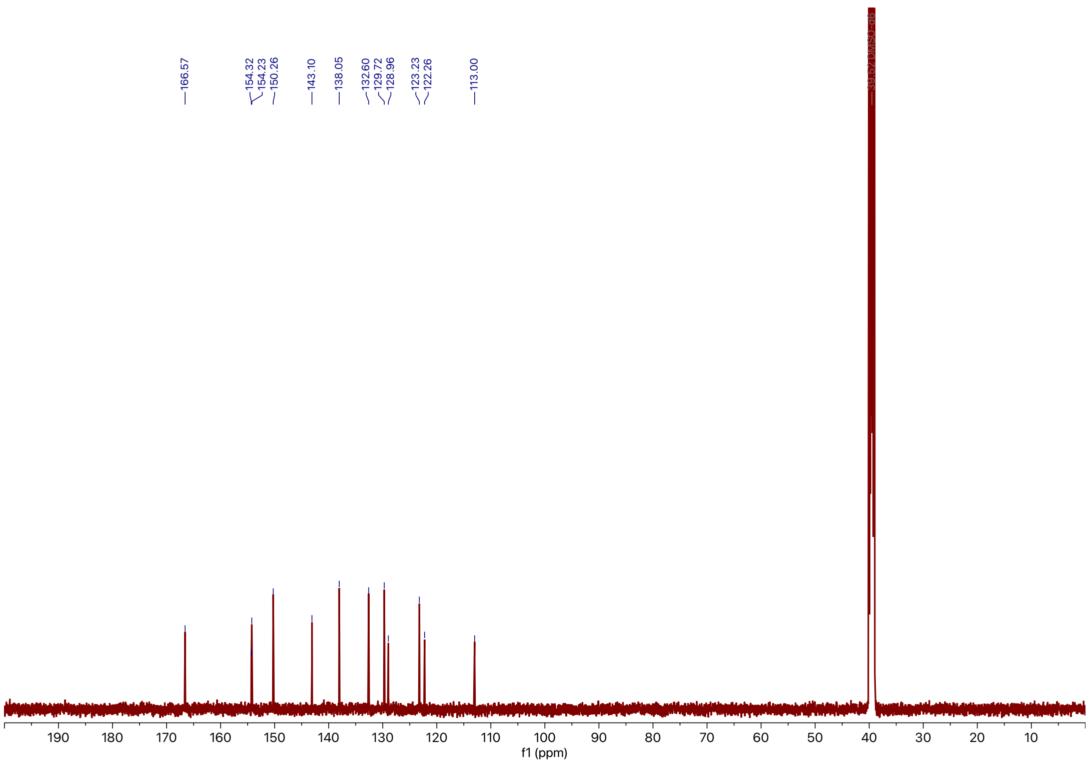

**Supporting Figure 57:** $^{13}$C NMR spectra of **23**

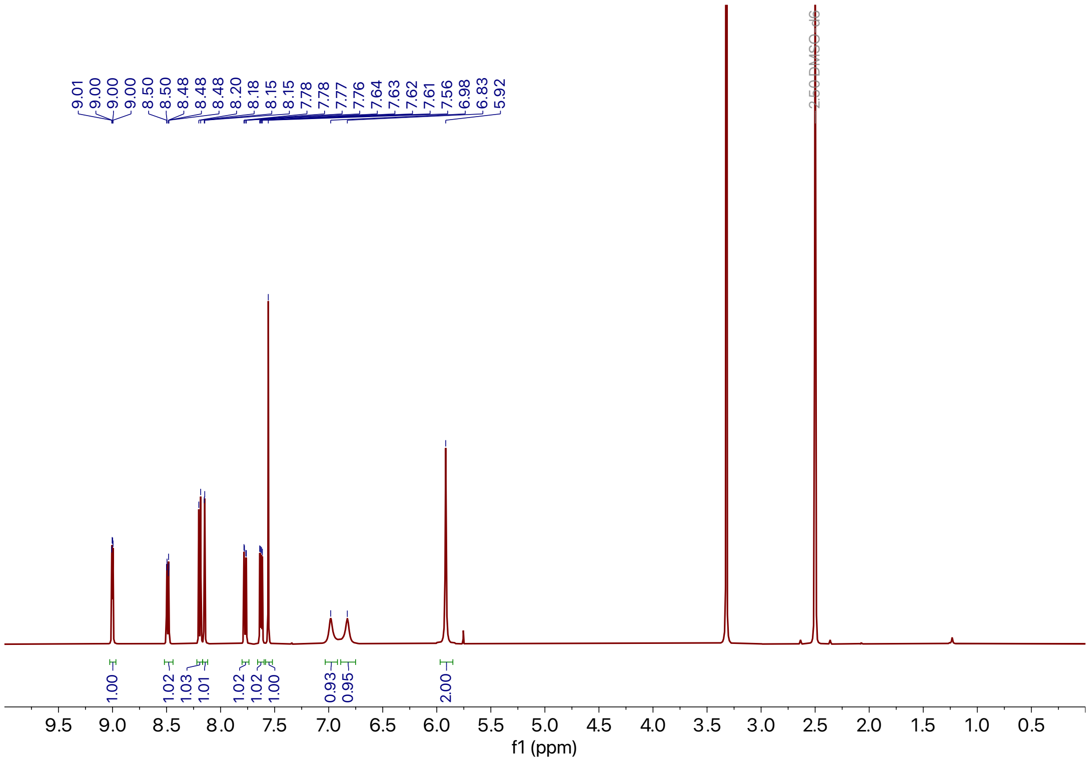

**Supporting Figure 58:** $^{1}$H NMR spectra of **24**

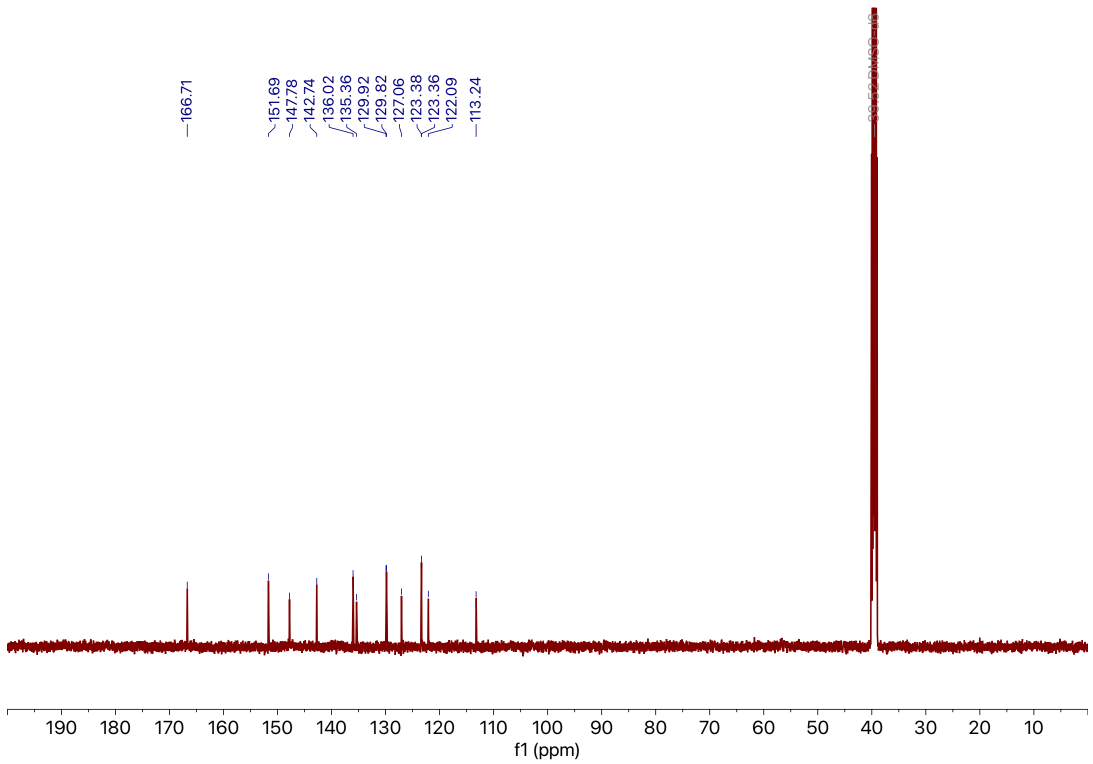

**Supporting Figure 59:** $^{13}$C NMR spectra of **24**

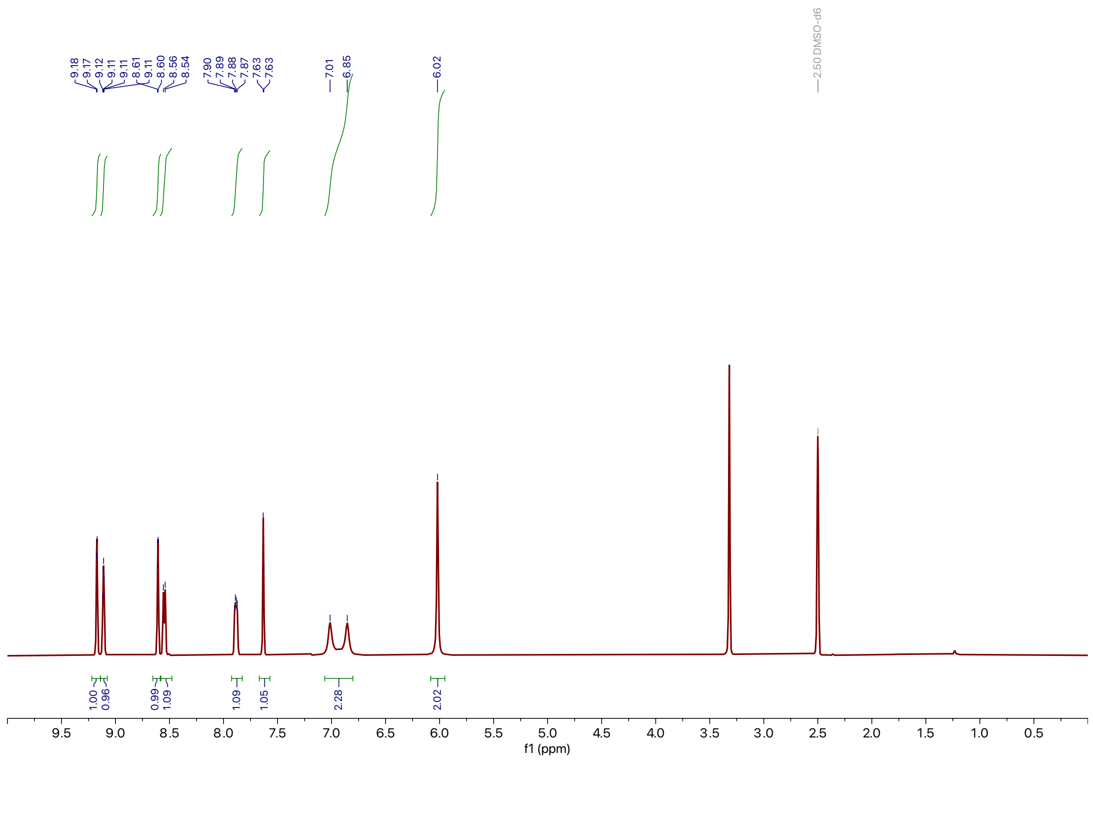

**Supporting Figure 60:** $^{1}$H NMR spectra of **25**

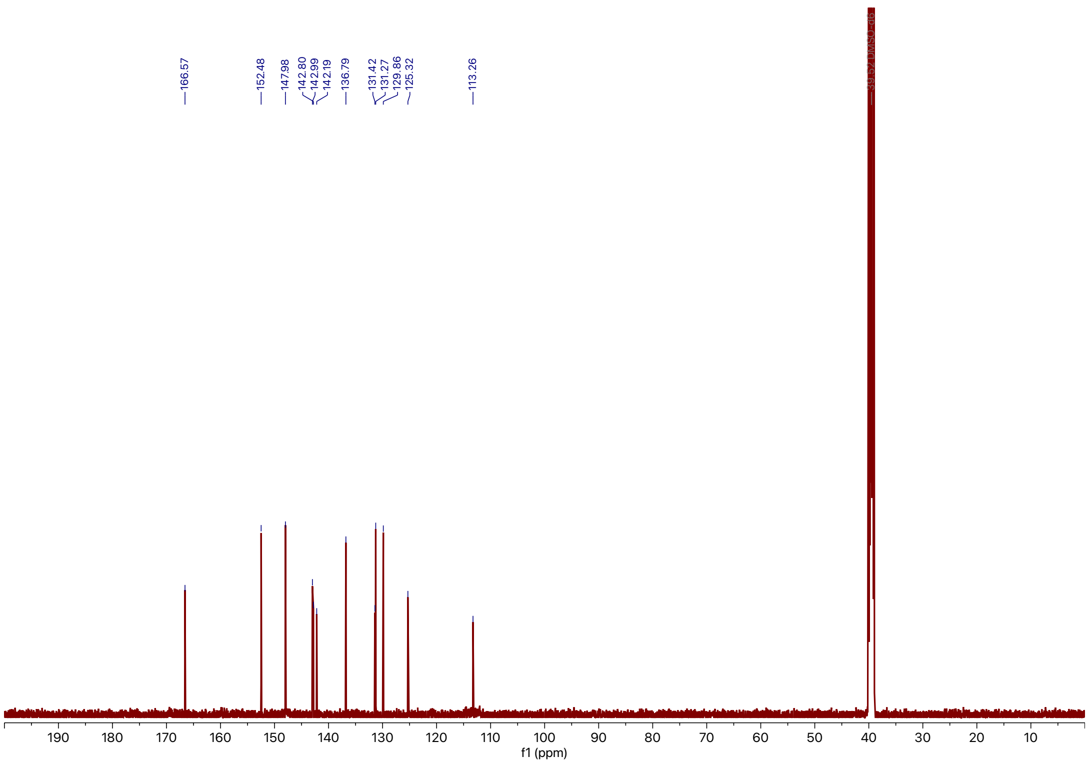

**Supporting Figure 61:** $^{13}$C NMR spectra of **25**

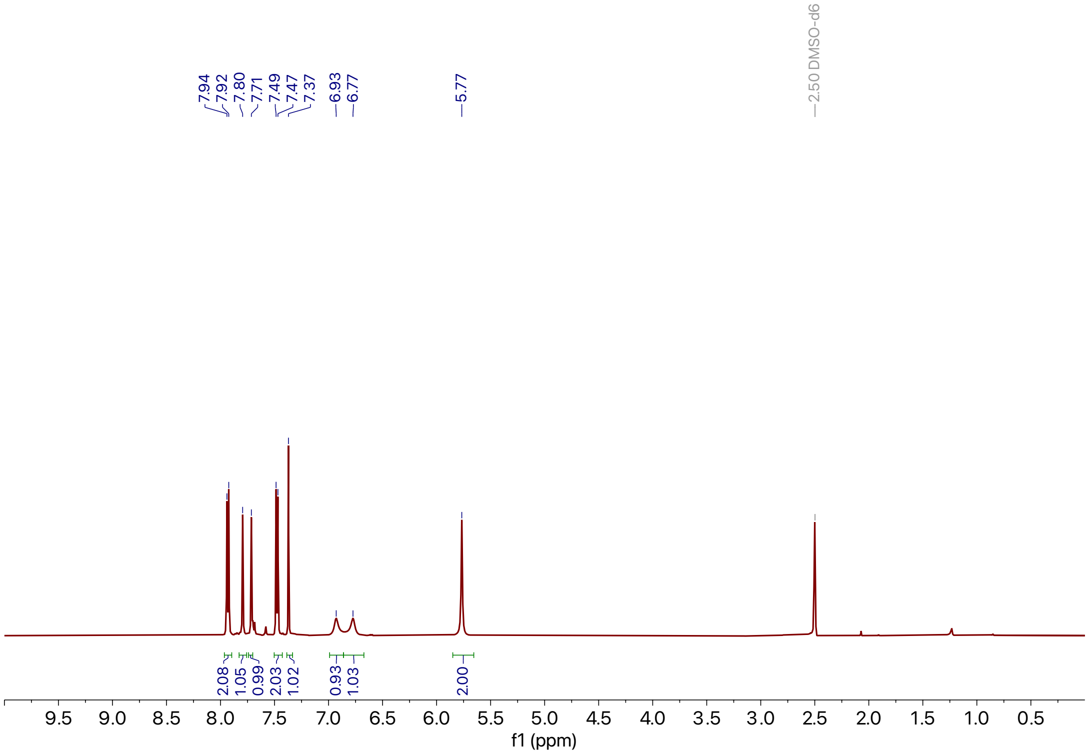

**Supporting Figure 62:** $^{1}$H NMR spectra of **26**

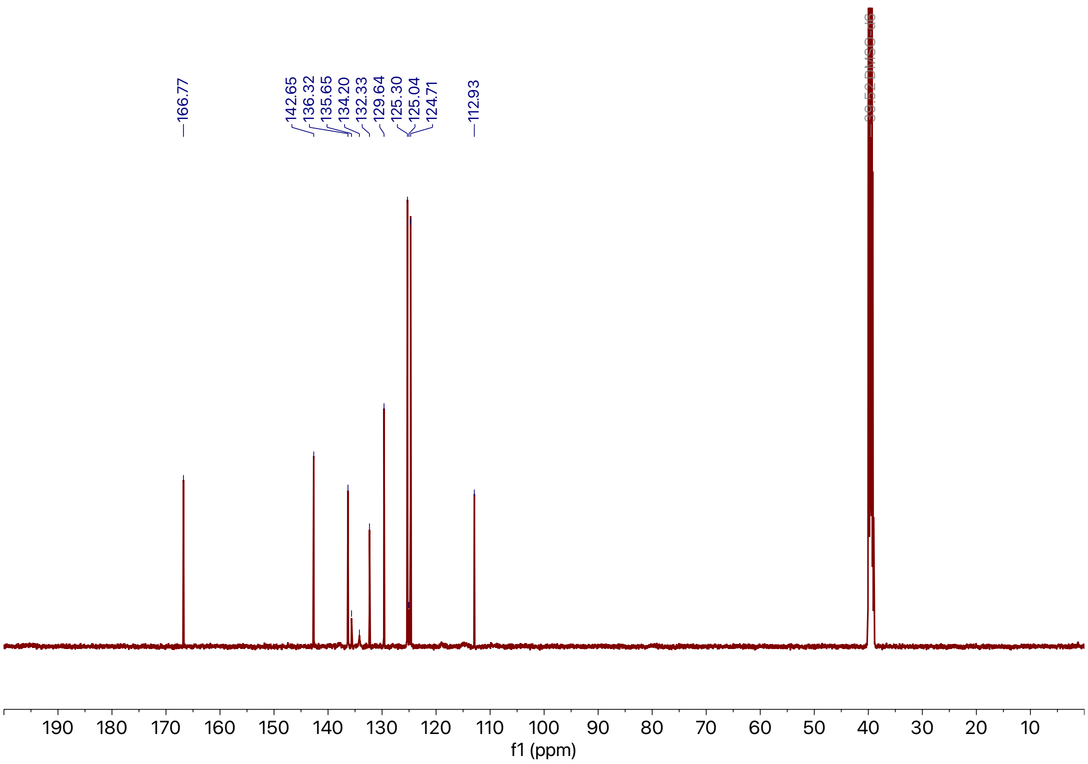

**Supporting Figure 63:** $^{13}$C NMR spectra of **26**

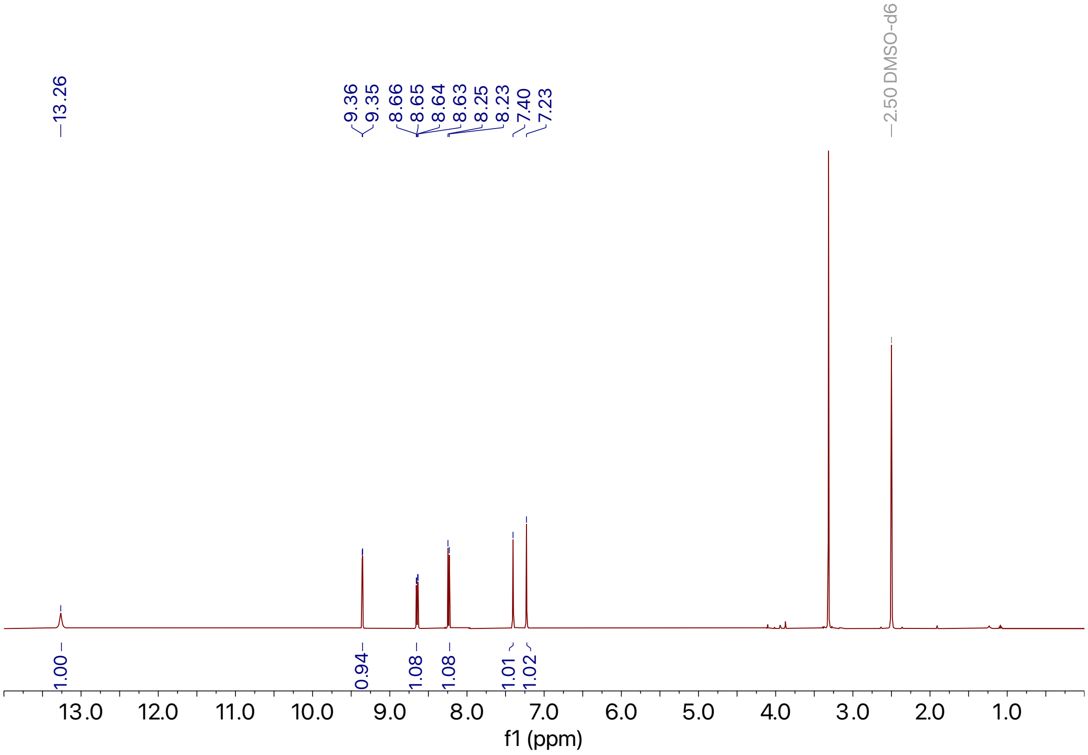

**Supporting Figure 64:** $^{1}$H NMR spectra of **S1**

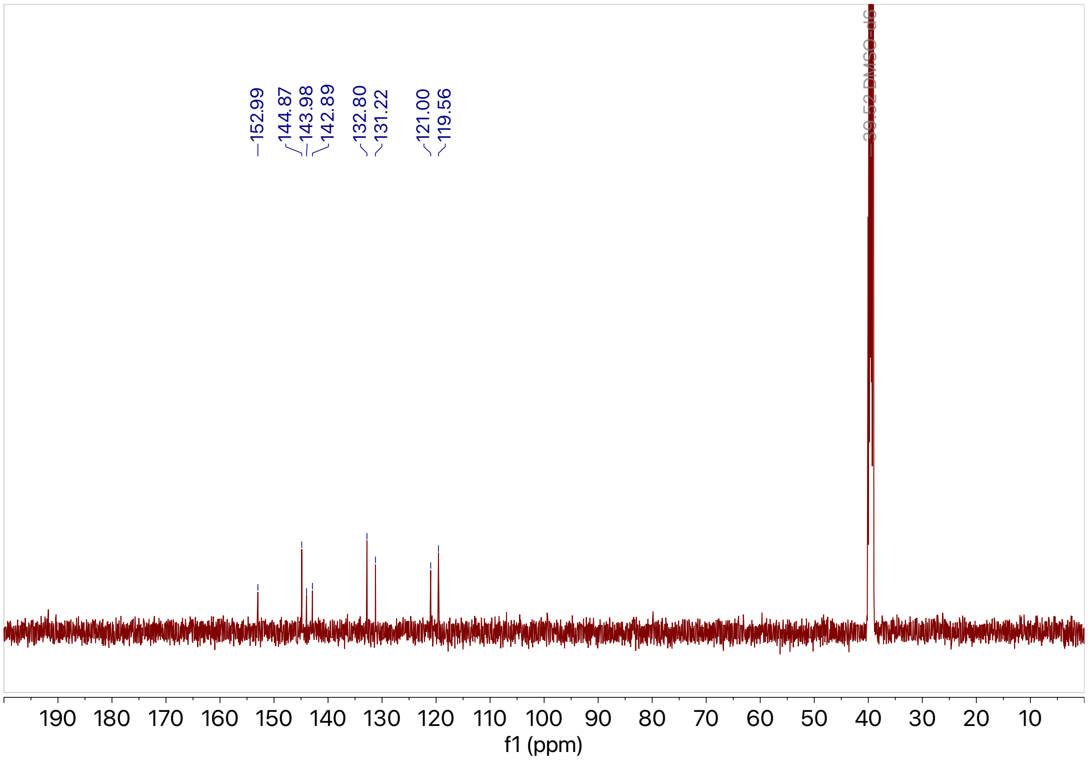

**Supporting Figure 65:** $^{13}$C NMR spectra of **S1**

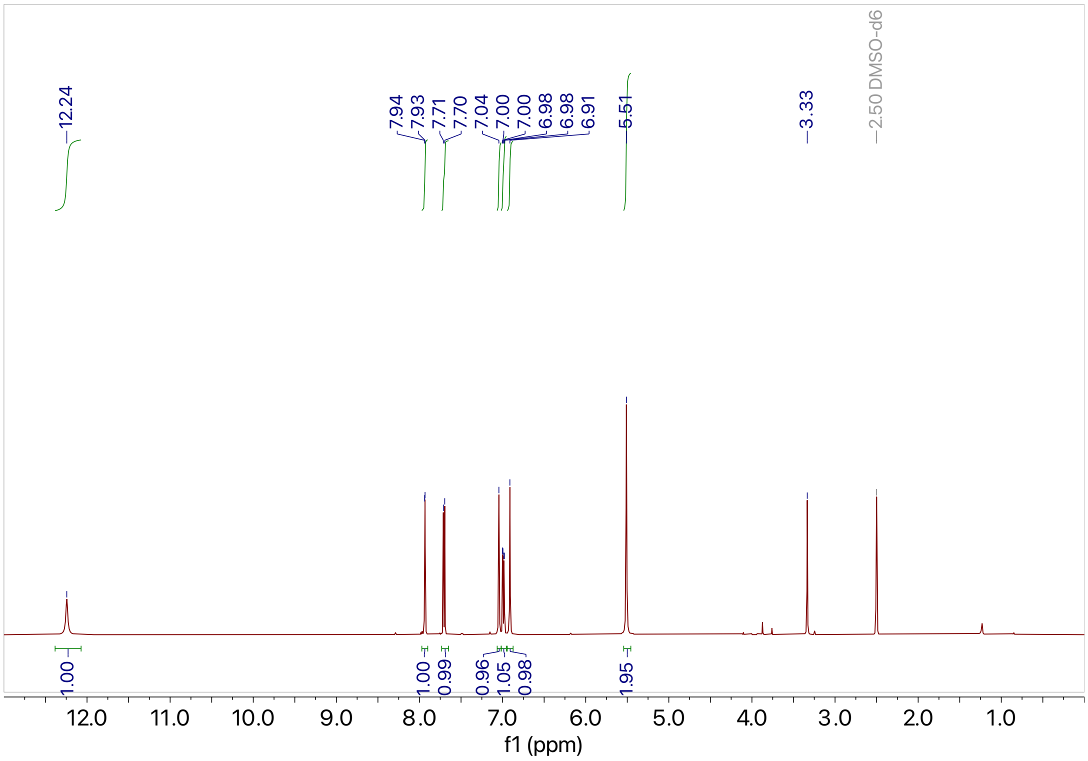

**Supporting Figure 66:** $^{1}$H NMR spectra of **S2**

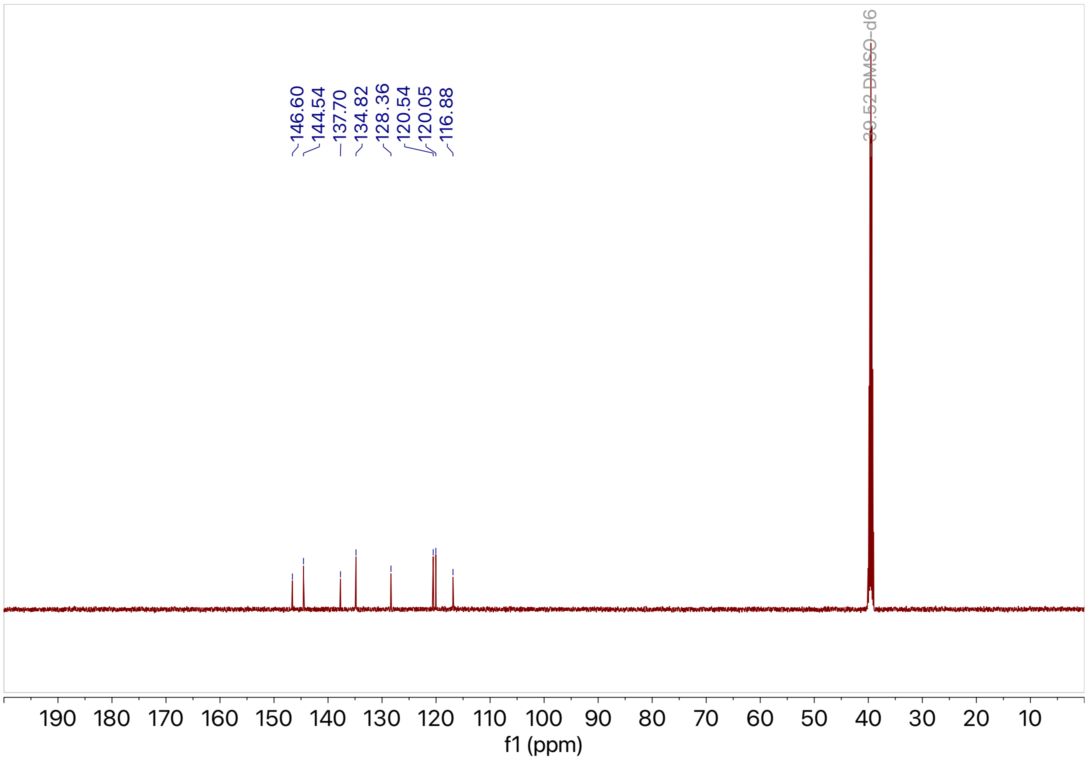

**Supporting Figure 67:** $^{13}$C NMR spectra of **S2**

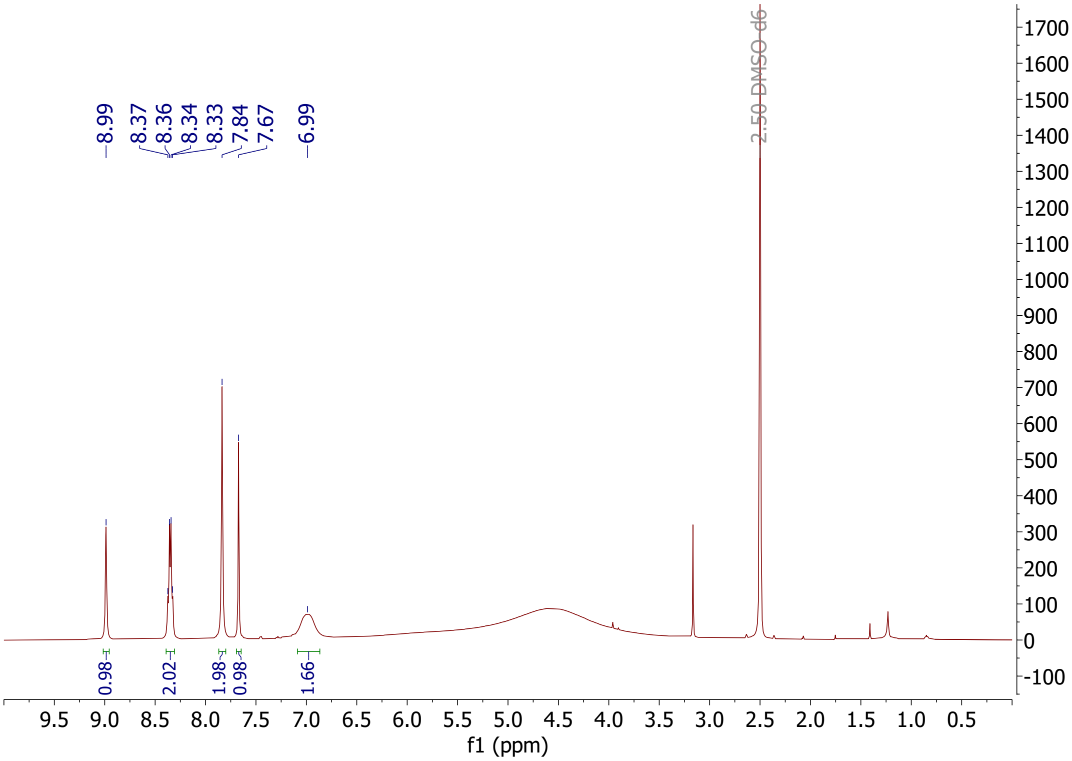

**Supporting Figure 68:** $^{1}$H NMR spectra of **27**

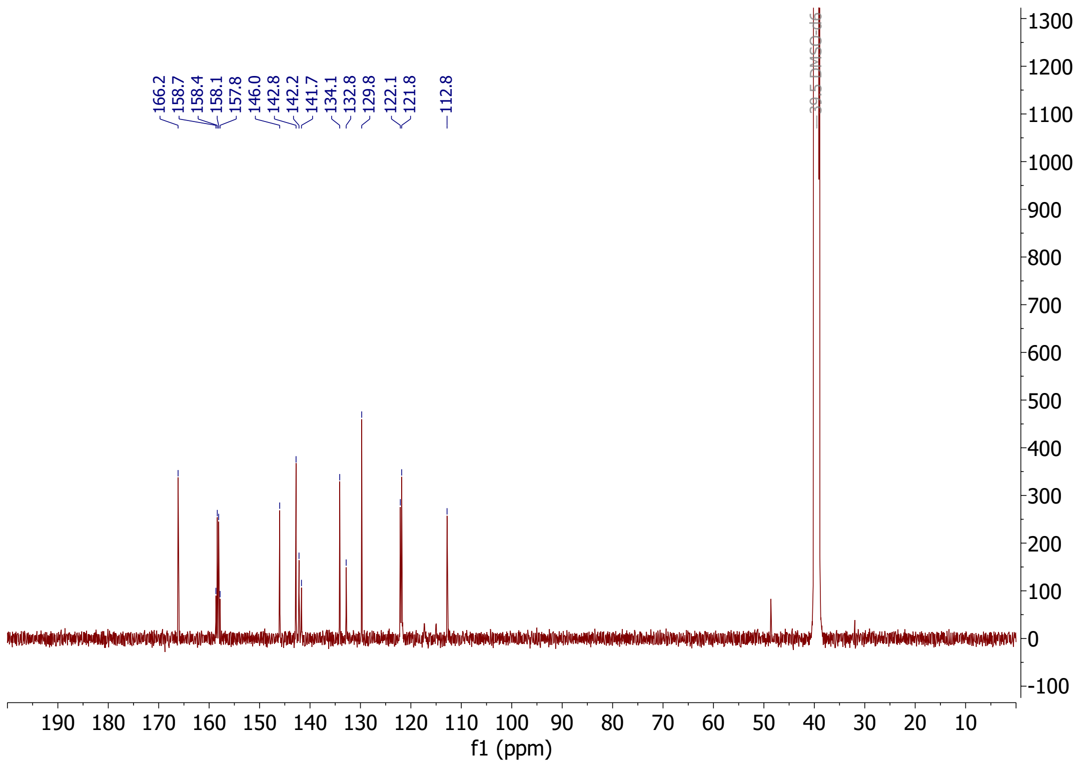

**Supporting Figure 69:** $^{13}$C NMR spectra of **27**

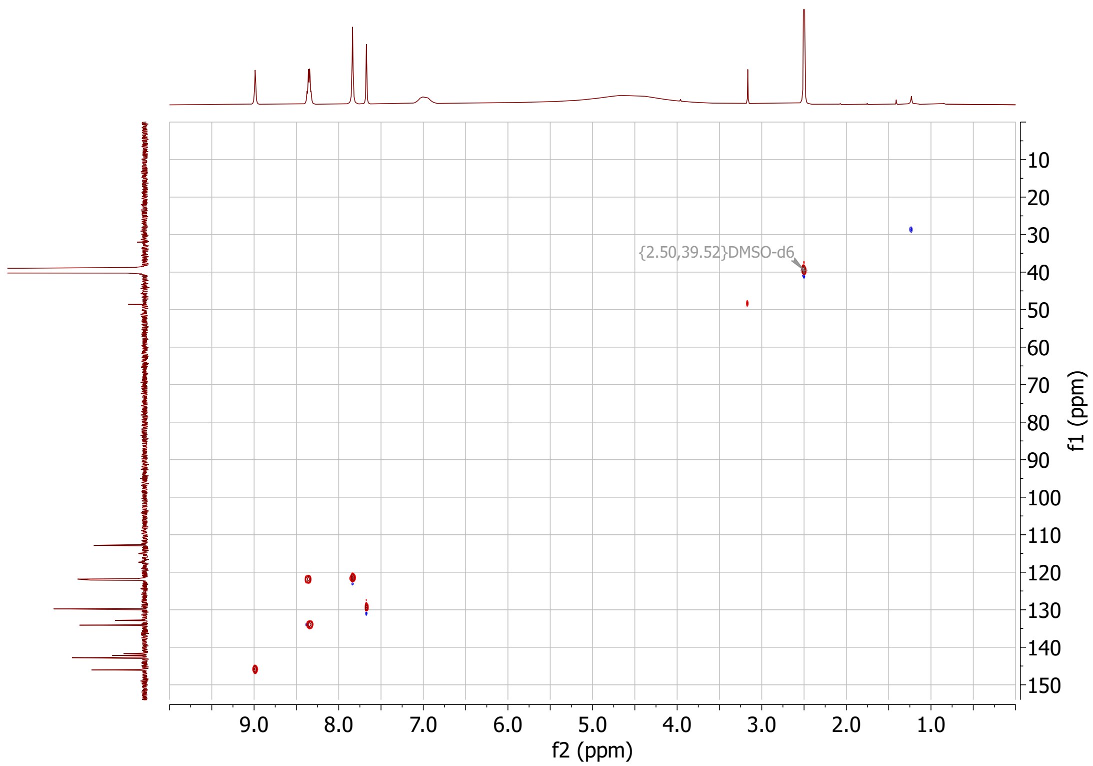

**Supporting Figure 70:** HSQC spectra of **27**

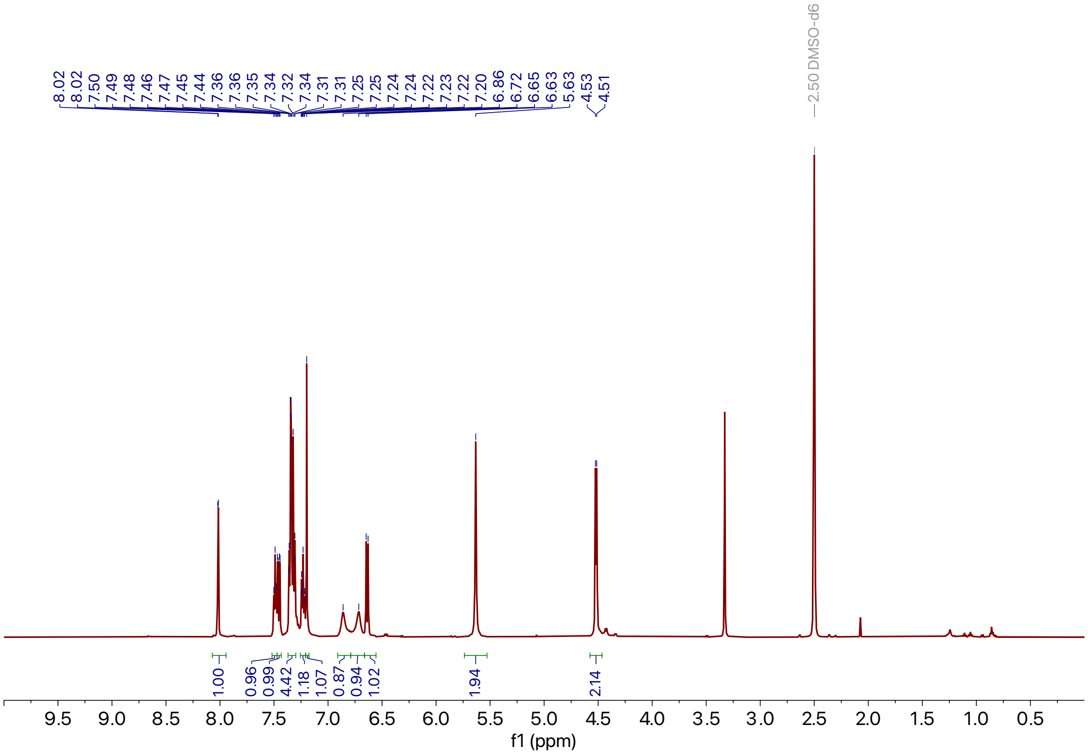

**Supporting Figure 71:** $^{1}H$ NMR spectra of **28**

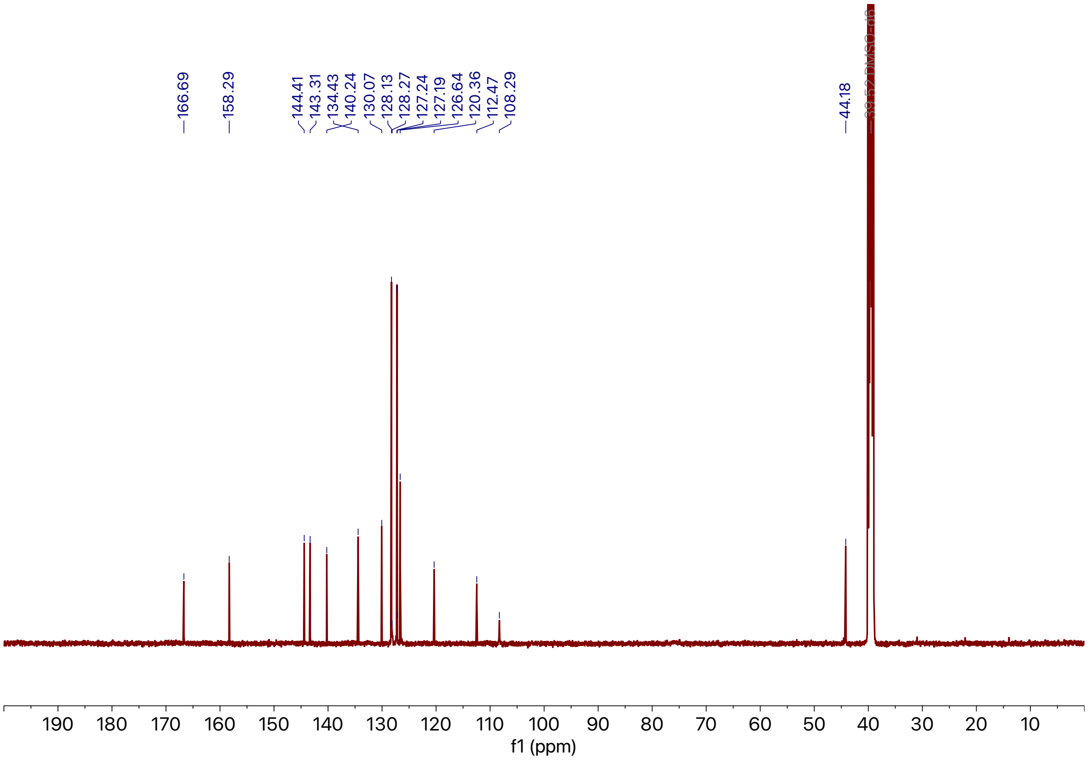

**Supporting Figure 72:** $^{13}C$ NMR spectra of **28**

#### References

[1] A. R. Ferré-D'Amaré and J. A. Doudna, Nucleic acids research **24**, 977 (1996).

[2] C. P. Jones and A. R. Ferré-D'Amaré, The EMBO journal **33**, 2692 (2014).

[3] J. J. Trausch, J. G. Marcano-Velázquez, M. M. Matyjasik, and R. T. Batey, Chemistry & biology **22**, 829 (2015).

[4] A. J. McCoy, R. W. Grosse-Kunstleve, P. D. Adams, M. D. Winn, L. C. Storoni, and R. J. Read, Journal of applied crystallography **40**, 658 (2007).

[5] D. Liebschner, P. V. Afonine, M. L. Baker, G. Bunkóczi, V. B. Chen, T. I. Croll, B. Hintze, L.-W. Hung, S. Jain, A. J. McCoy, et al., Acta Crystallographica Section D: Structural Biology **75**, 861 (2019).

[6] C. P. Jones, G. Piszczek, and A. R. Ferré-D'Amaré, Microcalorimetry of Biological Molecules: Methods and Protocols , 75 (2019).

[7] B. Tran, P. Pichling, L. Tenney, C. M. Connelly, M. H. Moon, A. R. Ferré-D'Amaré, J. S. Schneekloth, and C. P. Jones, Cell chemical biology **27**, 1241 (2020).

[8] J. L. Childs-Disney, X. Yang, Q. M. Gibaut, Y. Tong, R. T. Batey, and M. D. Disney, Nature Reviews Drug Discovery **21**, 736 (2022).

[9] P. Tiwary and M. Parrinello, Physical review letters **111**, 230602 (2013).

[10] P. Tiwary and M. Parrinello, The Journal of Physical Chemistry B **119**, 736 (2015).

[11] D. Wang and P. Tiwary, The Journal of Chemical Physics **154** (2021).

[12] T. Berger, Wiley Encyclopedia of Telecommunications (2003).

[13] J. M. L. Ribeiro, P. Bravo, Y. Wang, and P. Tiwary, The Journal of chemical physics **149** (2018).

[14] D. P. Kingma and M. Welling, arXiv preprint arXiv:1312.6114 (2013).

[15] T. D. Goddard, C. C. Huang, E. C. Meng, E. F. Pettersen, G. S. Couch, J. H. Morris, and T. E. Ferrin, Protein Science **27**, 14 (2018).

[16] M. R. Tucker, S. Piana, D. Tan, M. V. LeVine, and D. E. Shaw, The Journal of Physical Chemistry B **126**, 4442 (2022).

[17] V. T. Lim, D. F. Hahn, G. Tresadern, C. I. Bayly, and D. L. Mobley, F1000Research **9** (2020).

[18] L. J. Kinnaman, R. M. Roller, and C. S. Miller, Journal of Chemical Education **95**, 888

(2018).

[19] W. G. Hoover, Physical review A **31**, 1695 (1985).

[20] M. Parrinello and A. Rahman, Physical review letters **45**, 1196 (1980).

[21] Y. Jiao, B.-T. Xin, Y. Zhang, J. Wu, X. Lu, Y. Zheng, W. Tang, and X. Zhou, European Journal of Medicinal Chemistry **90**, 170 (2015).
